# BAP1 Haploinsufficiency Predicts a Distinct Immunogenic Class of Malignant Peritoneal Mesothelioma

**DOI:** 10.1101/243477

**Authors:** Raunak Shrestha, Noushin Nabavi, Yen-Yi Lin, Fan Mo, Shawn Anderson, Stanislav Volik, Hans H. Adomat, Dong Lin, Hui Xue, Xin Dong, Robert Shukin, Robert H. Bell, Brian McConeghy, Anne Haegert, Sonal Brahmbhatt, Estelle Li, Htoo Zarni Oo, Antonio Hurtado-Coll, Ladan Fazli, Joshua Zhou, Yarrow McConnell, Andrea McCart, Andrew Lowy, Gregg B. Morin, Tianhui Chen, Mads Daugaard, S. Cenk Sahinalp, Faraz Hach, Stephane Le Bihan, Martin E. Gleave, Yuzhuo Wang, Andrew Churg, Colin C. Collins

## Abstract

**Background:** Malignant Peritoneal Mesothelioma (PeM) is a rare and fatal cancer that originates from the peritoneal lining of the abdomen. Standard treatment of PeM is limited to cytoreductive surgery and/or chemotherapy, and no effective targeted therapies for PeM exist. Some immune checkpoint inhibitor studies of mesothelioma have found positivity to be associated with a worse prognosis.

**Methods:** To search for novel therapeutic targets for PeM, we performed a comprehensive integrative multi-omics analysis of the genome, transcriptome, and proteome of 19 treatment-naïve PeM, and in particular we examined *BAP1* mutation and copy-number status and its relationship to immune checkpoint inhibitor activation.

**Results:** We found that PeM could be divided into tumors with an inflammatory tumor microenvironment and those without, and that this distinction correlated with haploinsufficiency of *BAP1*. To further investigate the role of *BAP1*, we used our recently developed cancer driver gene prioritization algorithm, HIT’nDRIVE, and observed that PeM with *BAP1* haploinsufficiency form a distinct molecular subtype characterized by distinct gene expression patterns of chromatin remodeling, DNA repair pathways, and immune checkpoint receptor activation. We demonstrate that this subtype is correlated with an inflammatory tumor microenvironment and thus is a candidate for immune checkpoint blockade therapies.

**Conclusions:** Our findings reveal *BAP1* to be a potential, easily trackable prognostic and predictive biomarker for PeM immunotherapy that refines PeM disease classification. *BAP1* stratification may improve drug response rates in ongoing phase-I and II clinical trials exploring the use of immune checkpoint blockade therapies in PeM in which *BAP1* status is not considered. This integrated molecular characterization provides a comprehensive foundation for improved management of a subset of PeM patients.

## Background

Malignant mesothelioma is a rare but aggressive cancer that arises from internal membranes lining of the pleura and the peritoneum. While the majority of mesotheliomas are pleural in origin, the incidence of peritoneal mesothelioma (PeM) accounts for approximately 20-30% of all mesothelioma cases in the United States of America, and possibly higher in other countries [1]. Occupational asbestos exposure is a significant risk factor in the development of pleural mesothelioma (PM). However, epidemiological studies suggest that unlike PM, asbestos exposure plays a far smaller role in the etiology of PeM tumors [2]. More importantly, the incidence of PeM is skewed towards young women of childbearing ages rather than in old patients [1] making PeM a malignancy often associated with many years of life lost.

Previous studies in mesotheliomas has revealed that over 60% of mesotheliomas harbor *BRCA1* associated protein 1 (*BAP1*) inactivating mutation or copy number loss, making *BAP1* the most commonly altered gene in this malignancy [3–7]. BAP1 is a tumor suppressor and deubiquitinase, localized to the nucleus, known to regulate chromatin remodeling and maintain genome integrity [8, 9]. Furthermore, BAP1 localized in endoplasmic reticulum regulate calcium (Ca^2+^) flux to promote apoptosis [10]. Thus the combined reduced BAP1 nuclear and cytoplasmic activity results in the accumulation of DNA-damaged cells and greater susceptibility to development of malignancy. In addition, inactivating mutations of neurofibromin 2 (*NF2*) and cyclin dependent kinase inhibitor 2A (*CDKN2A*) are also relatively common, while other mutations are rare. Previous studies in PeM [11–18] have only focused on genomic information, therefore the downstream consequences of these genomic alterations is not well understood. Genome information coupled with transcriptome and proteome information is more likely to be successful in revealing potential therapeutic modalities.

Mesothelioma is typically diagnosed in the advanced stages of the disease. A combination of cytoreductive surgery (CRS) and hyperthermic intraperitoneal chemotherapy (HIPEC), sometimes followed by normothermic intraperitoneal or systemic chemotherapy (NIPEC) has recently emerged as a first-line treatment for PeM [19]. However, even with this regime, complete cytoreduction is hard to achieve and death ensues for many patients. Actionable molecular targets for PeM critical for precision oncology remains to be defined. Immune checkpoint blockade therapy in PM has recently gained traction [7, 20] given that 20-40% of PM cases are reported to show an inflammatory phenotype [21]. However, the role of immunostaining for PD-L1, the usual approach to predicting a response to immunotherapy for other tumor types, is controversial in PM, since positive stating has generally been associated with a worse prognosis, and it is unclear what marker should be used to predict tumors that may respond to immunotherapy.

Although, clinical trials typically lump PeM and PM together for immune checkpoint blockade [22–26], even less is known about PeM and immunotherapy. Thus, there has been no attempt to stratify PeM patients. In this study, we performed an integrated multi-omics analysis of the genome, transcriptome, and proteome of 19 PeM, predominantly of epithelioid subtype, and correlated these with tumor inflammation.

## Methods

### Patient Cohort

We assembled a cohort of 19 PeM from 18 patients (**Table 1**) undergoing CRS at Vancouver General Hospital (Vancouver, Canada), Mount Sinai Hospital (Toronto, Canada), and Moores Cancer Centre (San Diego, California, USA) (**Additional file 2: **Table S1****). We obtained 19 fresh-frozen primary treatment-naïve PeM tumor tissue and adjacent benign tissues or whole blood from the 18 patients. For one patient, MESO-18, two tumors from distinct sites were available. Immunohistochemical analyses using different biomarkers were evaluated by two independent pathologists (**Additional file 1: **Figures S1-S4****). Both pathologist categorized all 19 tumors as epithelioid PeM with a content of higher than 75% tumor cellularity. To the best of our knowledge this is the largest cohort of PeM subjected to an integrative multi-omics analysis.

**Table 1:** Peritoneal mesothelioma patients recruited for the study.

| **Tumor** | **Gender** | **Age (in years)** | **Asbestos Exposure** | **Subtype** | **WES** | **WTS** | **MS** |
| --- | --- | --- | --- | --- | --- | --- | --- |
| MESO-01 | Male | 59 | Unknown | BAP1-intact | Yes | No | Yes |
| MESO-02 | Female | 73 | Unknown | BAP1-del | Yes | Yes | Yes |
| MESO-03 | Female | 58 | Unknown | BAP1-intact | Yes | No | Yes |
| MESO-04 | Female | 70 | Unknown | BAP1-intact | Yes | No | Yes |
| MESO-05 | Male | 66 | Unknown | BAP1-del | Yes | Yes | Yes |
| MESO-06 | Male | 62 | No | BAP1-del | Yes | Yes | Yes |
| MESO-07 | Female | 34 | Unknown | BAP1-del | Yes | Yes | Yes |
| MESO-08 | Male | 54 | No | BAP1-intact | Yes | Yes | No |
| MESO-09 | Male | 62 | No | BAP1-del | Yes | Yes | Yes |
| MESO-10 | Female | 76 | No | BAP1-del | Yes | Yes | Yes |
| MESO-11 | Female | 46 | No | BAP1-intact | Yes | Yes | Yes |
| MESO-12 | Female | 55 | No | BAP1-intact | Yes | Yes | Yes |
| MESO-13 | Male | 70 | No | BAP1-intact | Yes | Yes | Yes |
| MESO-14 | Female | 54 | No | BAP1-del | Yes | Yes | Yes |
| MESO-15 | Female | 55 | No | BAP1-intact | Yes | No | No |
| MESO-17 | Female | 47 | No | BAP1-del | Yes | Yes | Yes |
| MESO-18A | Female | 61 | No | BAP1-intact | Yes | Yes | Yes |
| MESO-18E | Female | 61 | No | BAP1-intact | Yes | Yes | Yes |
| MESO-19 | Male | 47 | Yes | BAP1-intact | Yes | Yes | No |

WES: Whole Exome Sequencing, WTS: Whole Transcriptome Sequencing, and MS: Mass-spectrometry

### Immunohistochemistry and histopathology

Freshly cut Tissue Micro Array (TMA) sections were analyzed for immunoexpression using Ventana Discovery Ultra autostainer (Ventana Medical Systems, Tucson, Arizona). In brief, tissue sections were incubated in Tris-EDTA buffer (CC1) at 37°C to retrieve antigenicity, followed by incubation with respective primary antibodies at room temperature or 37°C for 60-120 min. For primary antibodies, mouse monoclonal antibodies against CD8 (Leica, NCL-L-CD8-4B11, 1:100), CK5/Cytokeratin 5(Abcam, ab17130, 1:100), BAP1 (SantaCruz, clone C4, sc-28383, 1:50), rabbit monoclonal antibody against CD3 (Abcam, ab16669, 1:100), and rabbit polyclonal antibodies against CALB2/Calretinin (LifeSpan BioSciences, LS-B4220, 1:20 dilution) were used. Bound primary antibodies were incubated with Ventana Ultra HRP kit or Ventana universal secondary antibody and visualized using Ventana ChromoMap or DAB Map detection kit, respectively. All stained slides were digitalized with the SL801 autoloader and Leica SCN400 scanning system (Leica Microsystems; Concord, Ontario, Canada) at magnification equivalent to x20. The images were subsequently stored in the SlidePath digital imaging hub (DIH; Leica Microsystems) of the Vancouver Prostate Centre. Representative tissue cores were manually identified by two pathologists.

### Whole exome sequencing

DNA was isolated from snap-frozen tumors with 0.2 mg/mL Proteinase K (Roche) in cell lysis solution using Wizard Genomic DNA Purification Kit (Promega Corporation, USA). Digestion was carried out overnight at 55°C before incubation with RNase solution at 37°C for 30 minutes and treatment with protein precipitation solution followed by isopropanol precipitation of the DNA. The amount of DNA was quantified on the NanoDrop 1000 Spectrophotometer and an additional quality check done by reviewing the 260/280 ratios. Quality check were done on the extracted DNA by running the samples on a 0.8% agarose/TBE gel with ethidium bromide.

For Ion AmpliSeq™ Exome Sequencing, 100ng of DNA based on Qubit^®^ dsDNA HS Assay (Thermo Fisher Scientific) quantitation was used as input for Ion AmpliSeq™ Exome RDY Library Preparation. This is a polymerase chain reaction (PCR)-based sequencing approach using 294,000 primer pairs (amplicon size range 225-275 bp), and covers >97% of Consensus CDS (CCDS; Release 12), >19,000 coding genes and >198,000 coding exons. Libraries were prepared, quantified by quantitative PCR (qPCR) and sequenced according to the manufacturer's instructions (Thermo Fisher Scientific). Samples were sequenced on the Ion Proton System using the Ion PI™ Hi-Q™ Sequencing 200 Kit and Ion PI™ v3 chip. Two libraries were run per chip for a projected coverage of 40M reads per sample.

### Somatic variant calling

Torrent Server (Thermo Fisher Scientific) was used for signal processing, base calling, read alignment, and generation of results files. Specifically, following sequencing, reads were mapped against the human reference genome hg19 using Torrent Mapping Alignment Program. Variants were identified by using Torrent Variant Caller plugin with the optimized parameters for AmpliSeq exome-sequencing recommended by Thermo Fisher. The variant call format (VCF) files from all sample were combined using GATK (3.2-2) [27] and all variants were annotated using ANNOVAR [28]. Only non-silent exonic variants including non-synonymous single nucleotide variations (SNVs), stop-codon gain SNVs, stop-codon loss SNVs, splice site SNVs and In-Dels in coding regions were kept if they were supported by more than 10 reads and had allele frequency higher than 10%. To obtain somatic variants, we filtered against dbSNP build 138 (non-flagged only) and the matched adjacent benign or blood samples sequenced in this study. Putative variants were manually scrutinized on the Binary Alignment Map (BAM) files through Integrative Genomics Viewer version 2.3.25 [29].

### Copy number aberration (CNA) analysis

Copy number changes were assessed using Nexus Copy Number Discovery Edition Version 9.0 (BioDiscovery, Inc., El Segundo, CA). Nexus NGS functionality (BAM ngCGH) with the FASST2 Segmentation algorithm was used to make copy number calls (a Circular Binary Segmentation/Hidden Markov Model approach). The significance threshold for segmentation was set at 5x10^-6^, also requiring a minimum of 3 probes per segment and a maximum probe spacing of 1000 between adjacent probes before breaking a segment. The log ratio thresholds for single copy gain and single copy loss were set at +0.2 and -0.2, respectively. The log ratio thresholds for gain of 2 or more copies and for a homozygous loss were set at +0.6 and -1.0, respectively. Tumor sample BAM files were processed with corresponding normal tissue BAM files. Reference reads per CNA point (window size) was set at 8000. Probes were normalized to median. Relative copy number profiles from exome sequencing data were determined by normalizing tumor exome coverage to values from whole blood controls.

### Whole transcriptome sequencing (RNA-seq)

Total RNA from 100μm sections of snap-frozen tissue was isolated using the mirVana Isolation Kit from Ambion (AM-1560). Strand specific RNA sequencing was performed on quality controlled high RIN value (>7) RNA samples (Bioanalyzer Agilent Technologies) before processing at the high throughput sequencing facility core at BGI Genomics Co., Ltd. (The Children's Hospital of Philadelphia, Pennsylvania, USA). In brief, 200ng of total DNAse treated RNA was first treated to remove the ribosomal RNA (rRNA) and then purified using the Agencourt RNA Clean XP Kit (Beckman Coulter) prior to analysis with the Agilent RNA 6000 Pico Chip to confirm rRNA removal. Next, the rRNA-depleted RNA was fragmented and converted to cDNA. Subsequent steps include end repair, addition of an ‘A’ overhang at the 3’ end, ligation of the indexing-specific adaptor, followed by purification with Agencourt Ampure XP beads. The strand specific RNA library prepared using TruSeq (Illumina Catalogue No. RS-122-2201) was amplified and purified with Ampure XP beads. Size and yield of the barcoded libraries were assessed on the LabChip GX (Caliper), with an expected distribution around 260 base pairs. Concentration of each library was measured with real-time PCR. Pools of indexed library were then prepared for cluster generation and PE100 sequencing on Illumina HiSeq 4000. The RNA-seq reads were aligned using STAR (2.3.1z) [30], onto the human genome reference (GRCh38) and the transcripts were annotated based on from Ensembl release 80 gene models. Only reads unique to one gene and which corresponded exactly to one gene structure, were assigned to the corresponding genes by using HTSeq [31]. Normalization of read counts was conducted by DESeq [32]. For detailed description see Supplementary Methods.

### Proteomics analysis using mass spectrometry

Fresh frozen samples dissected from tumor and adjacent normal were individually lysed in 50mM of HEPES pH 8.5, 1% SDS, and the chromatin content was degraded with benzonase. The tumor lysates were sonicated (Bioruptor Pico, Diagenode, New Jersey, USA), and disulfide bonds were reduced with DTT and capped with iodoacetamide. Proteins were cleaned up using the SP3 method [33, 34] (Single Pot, Solid Phase, Sample Prep), then digested overnight with trypsin in HEPES pH 8, peptide concentration determined by Nanodrop (Thermo) and adjusted to equal level. A pooled internal standard control was generated comprising of equal volumes of every sample (10μl of each of the 100μl total digests) and split into 3 equal aliquots. The labeling reactions were run as three TMT 10-plex panels (9+IS), then desalted and each panel divided into 48 fractions by reverse phase HPLC at pH 10 with an Agilent 1100 LC system. The 48 fractions were concatenated into 12 superfractions per panel by pooling every 4th fraction eluted resulting in a total 36 overall samples. These samples were analyzed with an Orbitrap Fusion Tribrid Mass Spectrometer (Thermo Fisher Scientific) coupled to EasyNanoLC 1000 using a data-dependent method with synchronous precursor selection (SPS) MS3 scanning for TMT tags. Based on ProteomeDiscoverer 2.1.1.21 (Thermo Fisher Scientific), we selected peptide-spectrum match (PSM) results with q-value <= 0.05, and extract proteins from both high and medium-confidence level after false discovery rate filtering for protein identification and quantification results. For detailed description see Supplementary Methods.

### Prioritization of driver genes using HIT’nDRIVE

Non-silent somatic mutation calls, CNA gain or loss, and gene-fusion calls were collapsed in gene-patient alteration matrix with binary labels. Gene-expression values were used to derive expression-outlier gene-patient outlier matrix using Generalized Extreme Studentized Deviate (GESD) test. STRING ver10 [35] protein-interaction network was used to compute pairwise influence value between the nodes in the interaction network. We integrated these genome and transcriptome data using HIT’nDRIVE algorithm [36]. Following parameters were used: *α* = 0.9, *β* = 0.6, *γ* = 0.8. We used IBM-CPLEX as the Integer Linear Programming (ILP) solver.

### Stromal and immune score

We used two sets of 141 genes (one each for stromal and immune gene signatures) as described in [37]. We used ‘inverse normal transformation’ method to transform the distribution of expression data into the standard normal distribution. The stromal and immune scores were calculated, for each sample, using the summation of standard normal deviates of each gene in the given set.

### Enumeration of tissue-resident immune cell types using mRNA expression profiles

CIBERSORT software [38] was applied to the RNA-seq gene-expression data to estimate the proportions of 22 immune cell types (B cells naive, B cells memory, Plasma cells, T cells CD8, T cells CD4 naive, T cells CD4 memory resting, T cells CD4 memory activated, T cells follicular helper, T cells gamma delta, T cells regulatory (Tregs), NK cells resting, NK cells activated, Monocytes, Macrophages M0, Macrophages M1, Macrophages M2, Dendritic cells resting, Dendritic cells activated, Mast cells resting, Mast cells activated, Eosinophils and Neutrophils) using LM22 dataset provided by CIBERSORT platform. Genes not expressed in any of the PeM tumor samples were removed from the LM22 dataset. The analysis was performed using 1000 permutation. The 22 immune cell types were later aggregated into 9 distinct groups.

## Results

### Landscape of somatic mutations in PeM

To investigate the landscape of somatic gene mutations in PeM, we performed high-coverage whole-exome sequencing of 19 tumors and 16 matched normal samples (**Additional file 2: **Table S1****). We achieved a mean coverage of 180x for cancerous samples and 96x for non-cancerous samples (**Additional file 2: **Table S2****). We identified 346 unique non-silent mutations affecting 202 unique genes (**Additional file 1: **Figure S5** and Additional file 2: **Table S3****). We observed an average of 0.015 protein-coding non-silent mutations per Mb per tumor sample.

We first identified driver genes of PeM using our recently developed computational algorithm HIT’nDRIVE [36]. Briefly, HIT’nDRIVE measures the potential impact of genomic aberrations on changes in the global expression of other genes/proteins which are in close proximity in a gene/protein-interaction network. It then prioritizes those aberrations with the highest impact as cancer driver genes. HIT’nDRIVE prioritized 25 unique driver genes in 15 PeM samples for which matched genome and transcriptome data were available (**Fig. 1 and Additional file 2: **Table S4****). Six genes (*BAP1, BZW2, ABCA7, TP53, ARID2*, and *FMN2*) were prioritized as drivers, harboring single nucleotide changes.

**Fig. 1.**
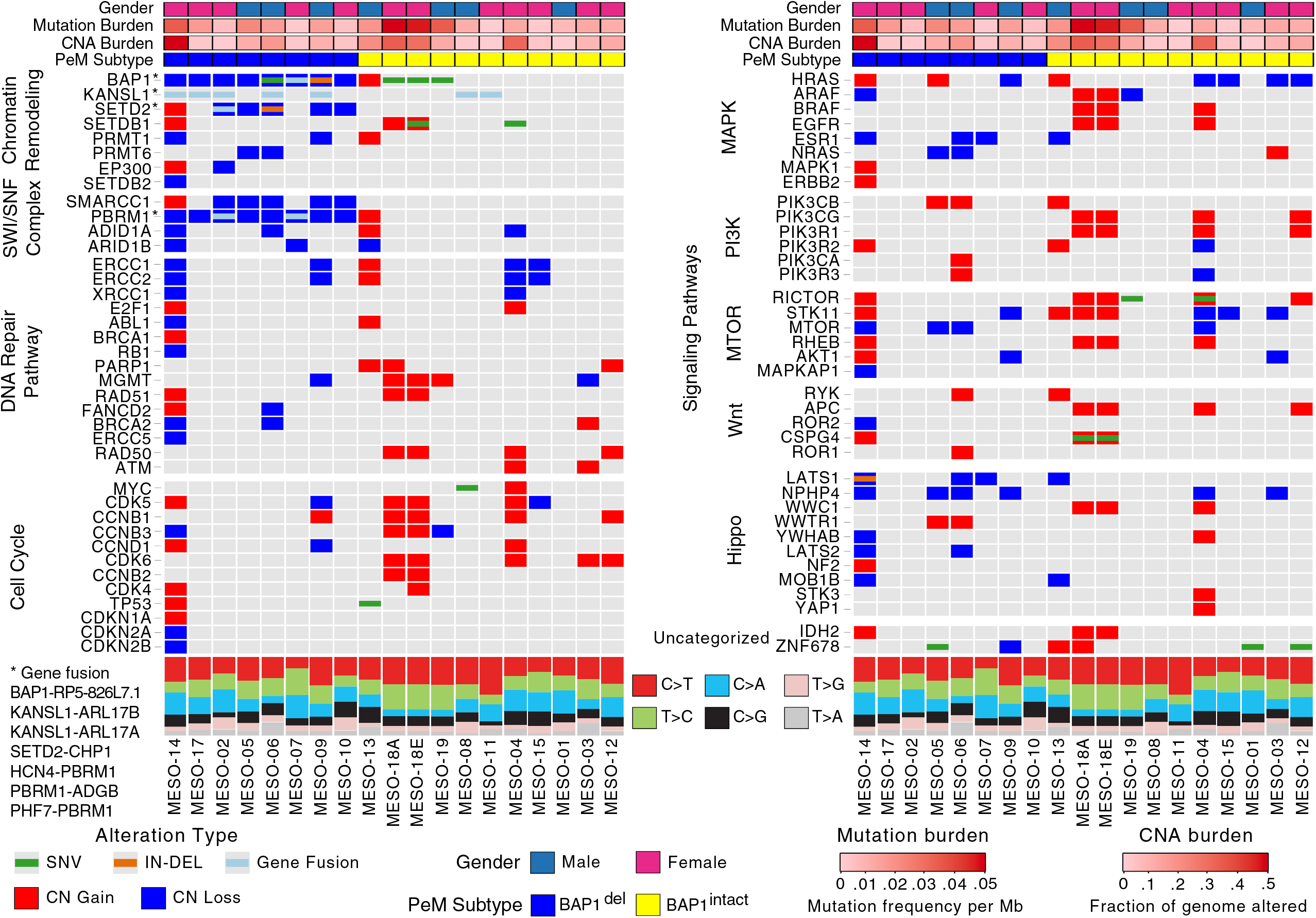
Integrated Molecular Comparison of Somatic Alterations across peritoneal mesothelioma subtypes. Somatic alterations status in PeM subtypes grouped by important cancer-pathways-chromatin remodeling, SWI/SNF complex, DNA Repair pathway, Cell cycle, MAPK, PI3K, MTOR, Wnt, and Hippo pathways. Somatic mutation status, copy-number status, gene-fusion, distribution of substitution mutation types, mutation burden, and copy-number aberration burden are indicated.

*BAP1* was the most frequently mutated gene (5 out of 19 tumors) in PeM. Among the five *BAP1* mutated cases, two cases (MESO-06 and MESO-09) were predicted to have inactivated BAP1, whereas despite *BAP1* mutation in three cases (MESO-18A/E and MESO-19) their mRNA transcripts were expressed in high levels (**Fig. 2C and Additional file 1: **Figures S6-S7****). We identified that all variants of *BAP1* (except a 42bp deletion in MESO-09) were expressed at the RNA level (**Additional file 2: **Table S16****). In addition, we identified mutations in genes such as *TP53, SETD2, SETDB1*, and *LATS1* each present in just a single case (**Fig. 1**).

**Fig. 2.**
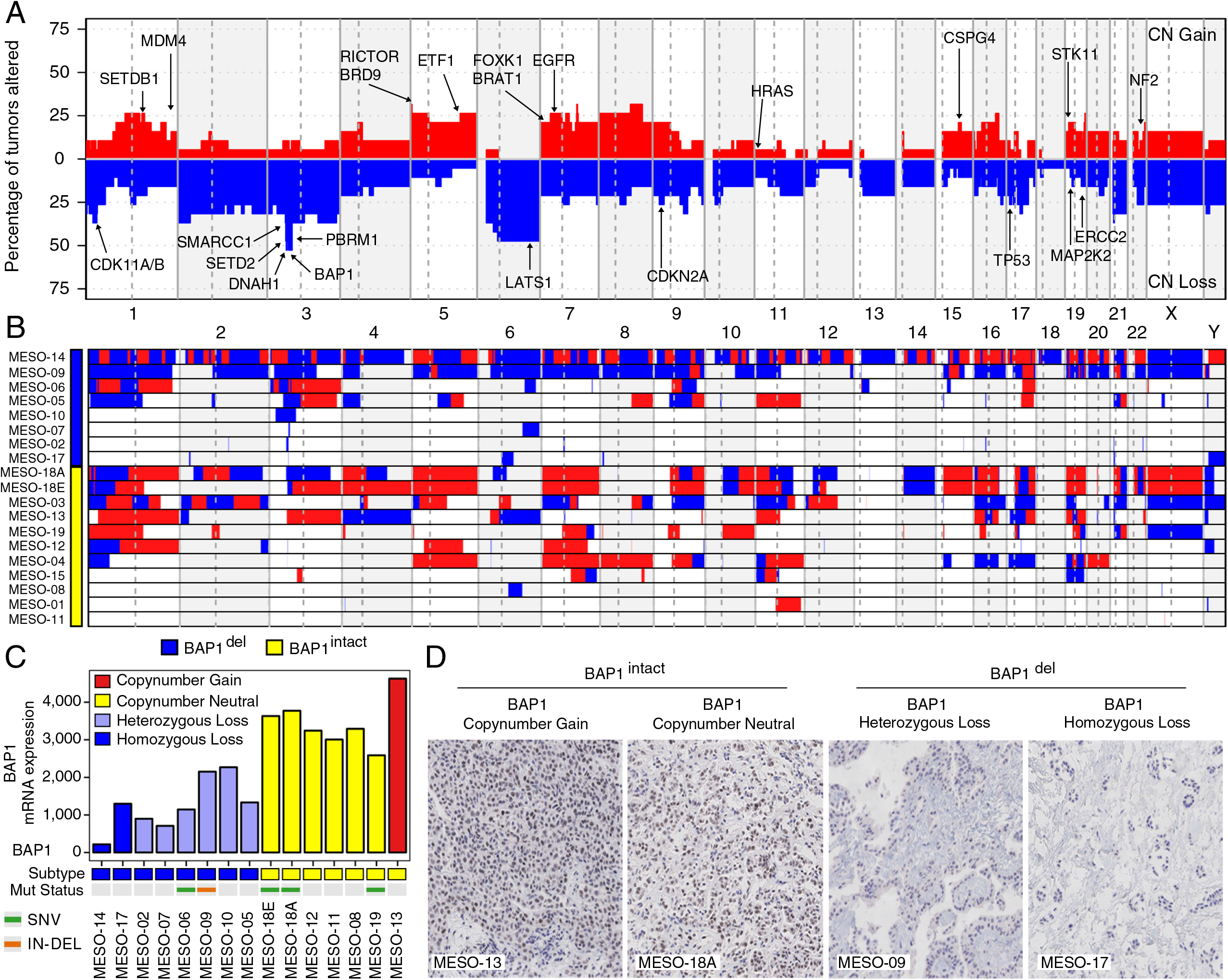
Landscape of copy number aberrations in PeM. (A) Aggregate copy-number alterations by chromosome regions in PeM. Important genes with copy-number changes are highlighted. (B) Sample-wise view of copy-number alterations in PeM. (C) mRNA expression pattern of *BAP1* across all PeM samples. (D) Detection of *BAP1* nuclear protein expression in PeM tumors by immunohistochemistry (Photomicrographs magnification - 20x).

### Copy number landscape in PeM

The aggregate somatic copy number aberration (CNA) profile of PeM is shown in **Fig. 2A-B**. We observed a total of 1,281 CNA events across all samples (**Additional file 2: **Table S5****). On an average, 10% of the protein-coding genome was altered per PeM. Interestingly, CNA burden in PeM was strongly correlated (R = 0.74) with its mutation burden (**Additional file 1: **Figure S9****).

Using HIT’nDRIVE, we identified genes in chromosome 3p21, *BAP1, PBRM1*, and *SETD2*, as key driver genes of PeM (**Fig. 1 and Additional file 2: **Table S4****). This region was also identified as significantly recurrent focal CNAs using GISTIC [39] algorithm (**Additional file 1: **Figure S9****). Chromosome 3p21 was deleted (homozygous or heterogzgous) in almost half of the tumors (8 of 19) in the cohort. Here, we call tumors with 3p21 (or *BAP1*) loss as *BAP1*^del^ and the rest of the tumors with 3p21 (or *BAP1*) copy-number intact as BAP1^intact^. Interestingly, *BAP1* mRNA transcripts in *BAP1*^del^ tumors were expressed at lower levels as compared to those in *BAP1*^intact^ tumors (*p*-value = 3x10^-4^) (**Fig. 2C**). We validated this using immunohistochemical (IHC) staining demonstrating lack of BAP1 nuclear staining in the tumors with *BAP1* homozygous deletion (**Fig. 2D**). Tumors with *BAP1* heterozygous loss still displayed BAP1 nuclear staining (**Additional file 1: **Figure S10****). We observed three *BAP1* mutated cases (MESO-18A/E and MESO-19) among *BAP1*^intact^ tumors. *BAP1* mRNA transcripts in these three tumors, were expressed at high levels (**Fig. 2C**). Furthermore, we found DNA copy loss of 3p21 locus to include four concomitantly deleted cancer genes - *BAP1, SETD2, SMARCC1*, and *PBRM1*, consistent with[5]. Copy-number status of these four genes was significantly correlated with their corresponding mRNA expression (**Additional file 1: **Figure S11****), suggesting that the allelic loss of these genes is associated with decreased transcript levels. These four genes are chromatin modifiers, and *PBRM1* and *SMARCC1* are part of SWI/SNF complex that regulates transcription of a number of genes.

### The global transcriptome and proteome profile of PeM

To segregate transcriptional subtypes of PeM, we performed total RNA-seq (Illumina HiSeq 4000) and its quantification of 15 PeM tumor samples for which RNA were available. Using principal-component analyses, we found that tumor samples in *BAP1*^intact^ and *BAP1*^del^ subtypes have distinct transcriptomic patterns; however, a few samples showed an overlapping pattern (**Additional file 1: **Figure S16A****).

We performed mass spectrometry (Fusion Orbitrap LC/MS/MS) with isobaric tagging for expressed peptide identification and its corresponding protein quantification using Proteome Discoverer for processing pipeline for 16 PeM tumors and 7 matched normal tissues. We identified 8242 unique proteins in 23 samples analyzed. We were surprised BAP 1 protein was however not detected in our MS experiment, likely due to inherent technical limitations with these samples and/or processing. Quality control analysis of in solution Hela digests also have very low BAP1 with only a single peptide observed in occasional runs. Unlike in transcriptome profiles, the proteome profiles of tumor samples in *BAP1*^intact^ and *BAP1*^del^ subtypes did not group into distinct clusters (**Additional file 1: **Figure S16B****).

Next we identified differentially expressed genes and proteins between *BAP1*^intact^ and *BAP1*^del^ subtypes (see Supplemental Methods). As expected, *BAP1, PBRM1* and *SMARCA4, SMARCD3* were among the top-500 differentially expressed genes. Many other important cancer-related genes were differentially expressed such as *CDK20, HIST1H4F, ERCC1, APOBEC3A, CDK11A, CSPG4, TGFB1, IL6, LAG3*, and *ATM*.

To identify the pathways dysregulated by the differentially expressed genes between the PeM subtypes, we performed geneset enrichment analysis (see Supplementary Methods). Intriguingly, we observed high concordance between pathways dysregulated by the two sets (mRNA and protein expression data) of top-500 differentially expressed genes and proteins (**Fig. 3A-B**). The unsupervised clustering of pathways revealed two distinct clusters for *BAP1*^del^ and *BAP1*^intact^ tumors. This indicates that the enriched pathways, between the patient groups, are also differentially expressed. *BAP1*^del^ tumors demonstrated elevated levels of RNA and protein metabolism as compared to *BAP1*^intact^ tumors. Many genes involved in chromatin remodeling and DNA damage repair were differently expressed between the groups (**Additional file 1: **Figures S20-S21****). Our data suggests that *BAP1*^del^ tumors have repressed DNA damage response pathways. Genes involved in cell-cycle and apoptotic pathways were observed to be highly expressed in *BAP1*^del^ patients. Furthermore, glucose and fatty-acid metabolism pathways were repressed in *BAP1*^del^ as compared to *BAP1*^intact^. More interestingly, we observed a striking difference in immune-system associated pathways between the PeM subtypes. Whereas *BAP1*^del^ tumors demonstrated strong activity of cytokine signaling and the innate immune system; MHC-I/II antigen presentation system and Adaptive immune system were active in *BAP1*^intact^ tumors.

**Fig. 3.**
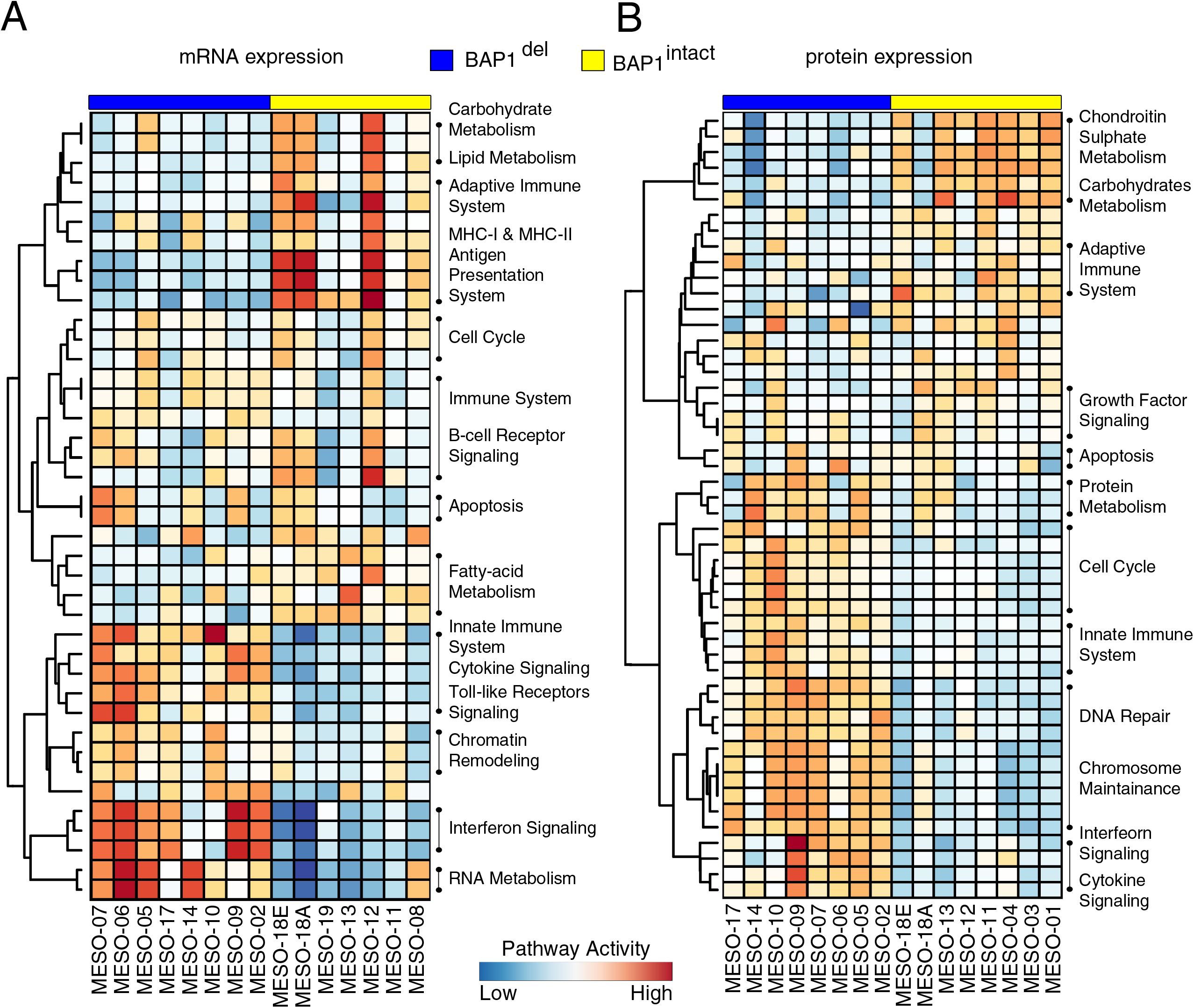
Transcriptome and proteome profile of PeM. Pathways enrichment of top-500 differentially expressed genes between PeM subtypes obtained using (A) mRNA expression and (B) protein expression. The colors on the heatmap shows pathway activity of the respective signaling pathways.

### *BAP1*^del^ subtype is correlated with tumor inflammation characterized by immune checkpoint receptor activation

Prompted by this finding, we next analyzed whether PeM were infiltrated with leukocytes. To assess the extent of leukocyte infiltration, we computed an expression (RNA-seq and protein) based score (**see Methods**) using the immune-cell and stromal markers proposed by [37]. We discovered that the immune marker gene score was strongly correlated with stromal marker gene score (**Fig. 4A**) suggesting possible leukocyte infiltration in PeM from the tumor microenvironment. Furthermore, using CIBERSORT [38] software, we computationally estimated leukocytes representation in the bulk tumor transcriptome. We observed massive infiltration of T cells in majority of the PeM (**Fig. 4B**). A subset of PeM had massive infiltration of B cells in addition to T cells. Interestingly, when we group the PeM by their *BAP1* aberration status, there was a marked difference in the proportion of infiltrated plasma cells, natural killer (NK) cells, mast cells, and B cells cells between the groups. Whereas the proportions of plasma cells, NK cells and B cells were less in the *BAP1*^del^ tumors, there was more infiltration of mast cells and T cells in *BAP1*^del^ tumors as compared to *BAP1*^intact^ tumors. We performed TMA IHC staining of CD3 and CD8 antibody on PeM tumors. We observed that *BAP1*^del^ PeM were positively stained for both CD3 and CD8 confirming infiltration of T cells in *BAP1*^del^ PeM (**Fig. 4C and Additional file 1: **Figures S22-S23****). Combined, this strongly indicates that PeM could be divided into tumors with an inflammatory tumor microenvironment and those without, and that this distinction correlated with *BAP1* haploinsufficiency.

**Fig. 4.**
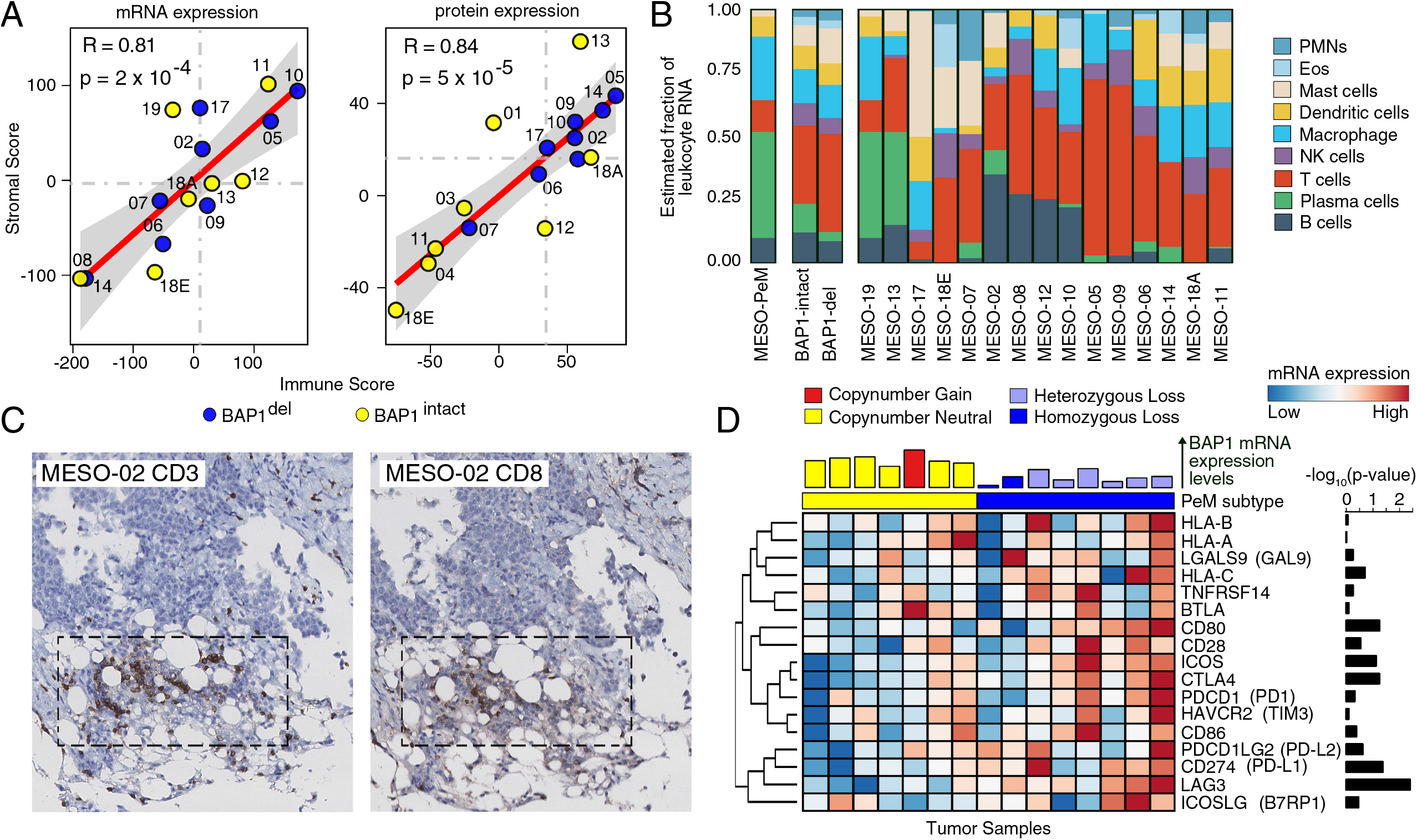
Immune cell infiltration in PeM. (A) Correlation between immune score and stromal score derived for each tumor sample using mRNA expression and protein expression. (B) Estimated relative mRNA fractions of leukocytes infiltrated in PeM tumors based on CIBERSORT analysis. (C) CD3 and CD8 immunohistochemistry showing immune cell infiltration on *BAP1*^del^ PeM (Photomicrographs magnification - 20x). (D) mRNA expression differences in immune checkpoint receptors - *LAG3, PD1, CTLA4, CD28, ICOS, BTLA*, and *HAVCR2* between PeM subtypes. Other genes in the figures are interacting receptors of the immune checkpoint markers mentioned above. The bar plot of the top of the heatmap indicates *BAP1* mRNA expression levels. The colors on the bar indicates *BAP1* copy-number status. The bar plot on the right represents the negative log10 of Wilcoxon signed-rank test *p*-value of individual immune checkpoint receptors computed between PeM subtypes. The expression levels are log2 transformed and mean normalized.

Finally, we surveyed PeM for expression of genes involved in immune checkpoint pathways. A number of immune checkpoint receptors were highly expressed in *BAP1*^del^ tumors relative to *BAP1*^intact^ tumors. These included *PDCD1* (*PD1*), *CD274* (*PD-L1*), *CD80, CTLA4, LAG3*, and *ICOS* (**Fig. 4D and Additional file 1: **Figure S30****) for which inhibitors are either clinically approved or are at varying stages of clinical trials. Notably, differential gene expression pattern of *LAG3, ICOS*, and *CTLA4* between the PeM subtypes suggests potential opportunities for immune checkpoint blockade beyond conventional PD1/PD-L1. Moreover, a number of MHC genes, immuno-inhibitor genes as well as immuno-stimulator genes were differentially expressed between *BAP1*^del^ and *BAP1*^intact^ tumors (**Additional file 1: **Figure S24****). Furthermore, we analyzed whether the immune checkpoint receptors were differentially expressed in tumors with and without 3p21 loss in pleural mesotheliomas (PM) from The Cancer Gene Atlas (TCGA) project [7]. Unlike in PeM, we did not observe a significant difference in immune checkpoint receptor expression between the PM groups (i.e. *BAP1*^del^ and *BAP1*^intact^) (**Additional file 1: **Figure S25****). These findings suggest that *BAP1*^del^ PeM tumors could potentially be targeted with immune-checkpoint inhibitors while PM tumors may less likely to respond.

## Discussion

In this study, we present a comprehensive integrative multi-omics analysis of malignant peritoneal mesotheliomas. Even though this is a rare disease we managed to amass a cohort of 19 tumors. Prior studies of mesotheliomas, performed using a single omic platform, have established *BAP1* inactivation as a key driver event in mesotheliomas. Our novel contribution to PeM is that we provide evidence from integrative multi-omics analyses that *BAP1* haploinsufficiency (*BAP1*^del^) forms a distinct molecular subtype of PeM. This subtype is characterized by distinct expression patterns of genes involved in chromatin remodeling, DNA repair pathway, and immune checkpoint activation. Moreover, *BAP1*^del^ subtype is correlated with inflammatory tumor microenvironment. Our results suggest that *BAP1*^del^ tumors might be prioritized for immune checkpoint blockade therapies. Thus *BAP1* is likely both prognostic and predictive biomarker for PeM enabling better disease stratification and patient treatment. Further corroborating our findings, *BAP1* status has been recently shown to be correlated with perturbed immune signaling in PM [7].

Loss of *BAP1* is known to alter chromatin architecture exposing the DNA to damage, and also impairing the DNA-repair machinery^8,38^. The DNA repair defects thus drive genomic instability and dysregulate tumor microenvironment[41]. DNA repair deficiency leads to the increased secretion of cytokines, including interferons that promote tumor-antigen presentation, and trigger recruitment of T lymphocytes to destroy tumor cells. As a response, tumor cells evade this immune-surveillance by increased expression of immune checkpoint receptors. The results presented here also indicate that PeM are infiltrated with immune-cells from the tumor microenvironment. Moreover, the *BAP1*^del^ subtype displays elevated levels of immune checkpoint receptor expression which strongly suggests the use of immune checkpoint inhibitors to treat this subtype of PeM. However, in a small subset of PM tumors in TCGA dataset, the loss of *BAP1* did not elevate expression of immune checkpoint marker genes. This warrants further investigation on the characteristics of these groups of PM.

The main challenge in mesothelioma treatment is that, all current efforts made towards testing new therapy options are limited to using therapies that have been proven successful in other cancer types, without a good knowledge of underlying molecular mechanisms of the disease. As a result of sheer desperation, some patients have been treated even though no targeted therapy for mesothelioma has been proven effective as yet. For example, a number of clinical trials exploring the use of immune checkpoint blockade (anti-PD1/PD-L1 or anti-CTLA4) in PM and/or PeM patients are currently under progress. The results of the first few clinical trials report either very low response rate or no benefit to the patients [22–24, 26, 42]. Notably, *BAP1* copy-number or mutation status were not assessed in these studies. Our study warrants further investigation of immune checkpoint molecules targeting beyond conventional PD1/PD-L1. We hypothesize based on this evidence presented that response rates for immune checkpoint blockade therapies in clinical trials for PeM will improve when patients are segregated by their *BAP1* copy-number status.

### Additional files

#### Additional file 1

Figure S1-S30. **Figure S1** Pathology of Peritoneal Mesothelioma. **Figure S2** Pathology of Peritoneal Mesothelioma. **Figure S3** TMA slides of PeM IHC stained for CK5. **Figure S4** TMA slides of PeM IHC stained for CALB2. **Figure S5** Summary of non-silent somatic mutation landscape of PeM. **Figure S6** Effect of BAP1 somatic mutation on resulting amino-acid chain. **Figure S7** Effect of BAP1 somatic mutation on resulting amino-acid chain. **Figure S8** Common somatic mutated genes in mesothelioma patients cohorts. **Figure S9** Landscape of copy number aberrations in PeM. **Figure S10** TMA slides of PeM IHC stained for BAP1. **Figure S11** Co-deletion of four cancer-associated genes in chromosomal region 3p21. **Figure S12** Consensus clustering of copy-number segment mean profiles of PeM. **Figure S13** Comparison of SCNA profile of peritoneal and pleural mesothelioma. **Figure S14** Gene Fusions in PeM. **Figure S15** Gene Fusions in PeM. **Figure S16** Transcriptome and proteome profile of PeM. **Figure S17** Consensus clustering of mRNA expression profiles of PeM. **Figure S18** Correlation between mRNA and protein expression profiles. **Figure S19** Consensus clustering of mRNA expression profiles of PeM. **Figure S20** Significant differentially expressed genes/proteins between molecular subtypes of PeM. **Figure S21** Differentially expressed DNA-repair genes between PeM subtypes. **Figure S22** TMA slides of BAP1del PeM IHC stained for CD3 and CD8. **Figure S23** TMA slides of BAP1intact PeM IHC stained for CD3 and CD8. **Figure S24** mRNA expression pattern of genes involved in immune system between PeM subtypes. **Figure S25** Immune cell infiltration in Pleural Mesothelioma. **Figure S26** Distribution of the variant allele frequency of somatic mutations in PeM. **Figure S27** Consensus clustering of long noncoding RNA (lncRNA) expression profiles of PeM. **Figure S28** Consensus clustering of micro RNA (miRNA) expression profiles of PeM. **Figure S29** Differentially expressed genes obtained when MESO-18A and/or MESO-18E are removed from the analysis. **Figure S30** Bar-plot of mRNA expression of different immune checkpoint receptors mentioned in Fig.4D.

#### Additional file 2

Table S1-S16. **Table S1**. Clinical information associated with the PeM patients. **Table S2**. Quality control statistics for WES data. **Table S3**. Somatic Mutation in PeM tumors. **Table S4**. Driver genes prioritized by HIT'nDRIVE algorithm. **Table S5**. Copy number segmentation profiles of PeM tumors. **Table S6**. Significant copy number changed regions prioritized by GISTIC. **Table S7**. Significant differential copy number aberrated regions between PeM and PM. **Table S8**. Predicted gene fusion events in PeM tumors using deFuse algorithm. **Table S9**. List of gene fusion events validated using Sanger sequencing. **Table S10**. Differentially expressed transcripts between PeM molecular subtypes. **Table S11**. Differentially expressed proteins between PeM molecular subtypes. **Table S12**. Correlation between CNA-mRNA and CNA-Protein. **Table S13**. Differentially expressed CORUM complex measured using mRNA expression levels. **Table S14**. Differentially expressed CORUM complex measured using protein expression levels. **Table S15**. lncRNA correlated or anticorrelated with different immune checkpoint receptors (LAG3, ICOS, CD274 (PD-L1), CTLA4, and CD80) in the PeM samples analyzed in the study. **Table S16**. The description of mutations in BAP1 observed using whole exome sequencing (DNA-seq) and corresponding BAP1 expressed variants detected using whole transcriptome sequencing (RNA-seq).

## Supporting information

## Abbreviations

BAM: Binary alignment map
BAP1: BRCA1 associated protein 1
CDKN2A: cyclin dependent kinase inhibitor 2A
CNA: Copy number aberration
CRS: Cytoreductive surgery
GESD: Generalized extreme studentized deviate
HIPEC: Hyperthermic intraperitoneal chemotherapy
IHC: Immunohistochemical
ILP: Integer Linear Programming
NF2: Neurofibromin 2 (NF2)
NIP EC: Normothermic intraperitoneal chemotherapy
PCR: Polymerase chain reaction
PeM: Peritoneal mesothelioma
PM: Pleural mesothelioma
PSM: Peptide-spectrum match
qPCR: Quantitative PCR
SETD2: SET domain containing 2
SNV: Single nucleotide variation
SPS: Synchronous precursor selection
TMA: Tissue microarray
VCF: Variant call format

## Declarations

### Acknowledgement

We acknowledge the contribution and support of Drs. Jessica McAlpine, Anna Tinker, and Jeff Simko as well as Emily Taylor (Mount Sinai Hospital), Donald Donaldson, Rose Schweigert, Matthew Sturgen, Sarah Padilla at Vancouver Coastal Health for supporting and coordinating the procurement of PeM tumors. We are thankful for the support of regulatory and ethics bodies: Margaret Luk, Wylo Kyle, Jacqueline Lee, Zahra Karim, Nandita Chowdhury, Jessica Gagliardi, Suzanne Richardson, and Sheila O'Donoghue at Vancouver General Hospital; Chantal Lackan at Mount Sinai Hospital; Ida Deichaite and Oudone Sisanachandeng at Moores Cancer Centre. Thanks to Shane Colborne and Dr. Christopher Hughes for help and suggestions regarding mass spectrometry experiments and data analyses. The authors thank all members of the Collins', Wang’s, Hach’s, and Sahinalp’s labs for helpful suggestions. The results published here are in part based upon data generated by the TCGA Research Network: http://cancergenome.nih.gov/.

## Availability of data and materials

The whole-exome and whole-transcriptome sequencing data from this study is available in the European Genome-phenome Archive (EGA; https://ega-archive.org/) under accession number EGAS00001002820. The proteome data from mass spectrometry is available in the PRIDE Archive (https://www.ebi.ac.uk/pride/archive/) under accession number PXD008873.

## Author Contributions

R.S., N.N., S.LB., Y.W. A.C., and C.C.C. conceived the study. R.S. and N.N. performed data analysis and wrote the manuscript. N.N. and S.LB. managed the project. Y.L., F.M., S.A., S.V., H.H.A, R.H.B., and J.Z. performed data analysis. N.N., Ro.S., B.M., A.H., S.B., H.H.A, and G.B.M. performed the specimen processing, quality control, sequencing and mass-spectrometry experiments, and validation experiments. E.L., HZ.O., A.H., and L.F. constructed TMAs, performed IHC experiments, and TMA scoring. A.C. and HZ.O. reviewed the tissue slides. N.N., D.L., H.X., and X.D., constructed patient derived mouse xenograft. Y.M., A.M., and A.L. contributed clinical specimens and clinical data. M.D., T.C., S.C.S., F.H., S.LB., M.E.G., Y.W., A.C., and C.C.C. supervised the project, contributed scientific insights, and edited the manuscript.

## Ethics approval and consent to participate

This study was approved by the Institutional Review Board of the University of British Columbia and the Vancouver Coastal Health (REB Number. H1500902 and V15-00902). All samples and information were collected with written and signed informed consent from the participating patients.

## Consent for publication

Not applicable.

