## Supplementary material for "BAP1 Haploinsufficiency Predicts a Distinct Immunogenic Class of Malignant Peritoneal Mesothelioma"

Fig S30 Bar-plot of mRNA expression of different immune checkpoint receptors mentioned in Fig.4D. 50

### 1 Supplemental Methods

#### 1.1 Cohort Description and Pathology Evaluation

Primary untreated peritoneal mesothelioma (PeM) tumors and matched benign samples were obtained from cancer patients undergoing cytoreductive surgeries following protocols approved by the Clinical Research Ethics Board of the Vancouver General Hospital (Vancouver, BC, Canada), Mount Sinai Hospital (Toronto, ON, Canada), and Moores Cancer Centre (San Diego, CA, USA). This study was approved by the Institutional Review Board of the University of British Columbia and Vancouver Coastal Health (REB No. H15-00902 and V15-00902). All patients signed a formal consent form approved by the respective institutional ethics board. Material transfer agreements between institutions were fully executional prior to sample acquisition.

The PeM cohort included fresh frozen tumors from PeM patients who underwent cytoreductive surgeries prior to therapy, spanning from 2013-2016. Demographics (age and gender), histopathologic information, and asbestos exposure and tobacco use information were also collected from patient's medical records.

Histologic parameters and pathological scoring of tumors confirming PeM was established by three independent pathologists. Hematoxylin and eosin (H&E) and immunostained formalin-fixed paraffin-embedded (FFPE) slides were reviewed by at least two specialized pathologists to diagnose PeM and its subtype. H&E staining was used to determine the highest tumor cellularity ($\geq 75\%$) from sections for sequencing.

The surgical resections were snap frozen and processed at respective institutions. The tumors have a companion normal tissue specimen (either adjacent normal tissue or peripheral blood previously extracted for germline DNA control). Each tumor specimen was approximately $1cm^3$ in size and weighed between 100-300 mg. Specimen were shipped overnight on dry ice that maintained an average temperature of less than -80$^o$C. Upon receipt, the tissues were sectioned into 5 slices for Deoxyribonucleic acid (DNA), ribonucleic acid (RNA), and protein extraction as well as construction of tissue microarrays (TMA).

#### 1.2 Construction of tissue microarrays (TMA)

FFPE tissue blocks were retrieved from the archives of the Department of Pathology, Vancouver General Hospital (Vancouver, Canada). H&E stained slides from each block were reviewed by two pathologists to identify tumor areas. TMAs were constructed with 1 mm diameter tissue cores from representative tumor areas from FFPE blocks. Cores were transferred to a paraffin block using a semi-automated tissue array instrument (Pathology Devices TMArrayer, San Diego, CA). Duplicate tissue cores were taken from each specimen, resulting in a composite TMA block. Reactive mesothelial tissues from pleura were also included as benign controls. Following construction, $4\mu$m thick sections were cut for H&E and immunohistochemical staining.

#### 1.3 Exome sequencing

DNA was isolated from snap-frozen tumors with 0.2 mg/mL Proteinase K (Roche) in cell lysis solution using Wizard Genomic DNA Purification Kit (Promega Corporation, USA). Digestion was carried out overnight at 55$^{\circ}$C before incubation with RNase solution at 37$^{\circ}$C for 30 minutes and treatment with protein precipitation solution followed by isopropanol precipitation of the DNA. The amount of DNA was quantified on the nanodrop and an additional quality check done by reviewing the 260/280 ratios. Quality check was done on the extracted DNA by running the samples on a 0.8% agarose/TBE gel with ethidium bromide.

For Ion AmpliSeq™ Exome Sequencing, 100ng of DNA based on Qubit® dsDNA HS Assay (Thermo Fisher Scientific) quantitation was used as input for Ion AmpliSeq™ Exome RDY Library Preparation. This is a polymerase chain reaction (PCR)-based sequencing approach using 294,000 primer pairs (amplicon size range 225-275 bp), and covers >97% of Consensus CDS (CCDS; Release 12), >19,000 coding genes and >198,000 coding exons. Libraries were prepared, quantified by quatitative PCR (qPCR) and sequenced according to the manufacturer's instructions (Thermo Fisher Scientific). Samples were sequenced on the Ion Proton System using the Ion PI™ Hi-Q™ Sequencing 200 Kit and Ion PI™ v3 chip. Two libraries were run per chip for a projected coverage of 40M reads per sample.

#### 1.4 Somatic variant calling

Torrent Server (Thermo Fisher Scientific) was used for signal processing, base calling, read alignment, and generation of results files. Specifically, following sequencing, reads were mapped against the human reference genome hg19 using Torrent Mapping Alignment Program. The mean target coverage ranges from 78.62 to 226.44, thus sequencing depth ranges from 78 to 226X. Variants were identified by using Torrent Variant Caller plugin with the optimized parameters for AmpliSeq exome-sequencing recommended by Life Technologies. The variant call format (VCF) files from all sample were combined using GATK (3.2-2) (DePristo et al. 2011) and all variants were annotated using ANNOVAR (Wang et al. 2010). Only non-silent exonic variants including non-synonymous single nucleotide variations (SNVs), stop-codon gain SNVs, stop-codon loss SNVs, splice site SNVs and In-Dels in coding regions were kept if they were supported by more than 10 reads and had allele frequency higher than 10%. To obtain somatic variants, we filtered against dbSNP build 138 (non-flagged only) and the matched adjacent benign or blood samples sequenced in this study. Putative variants were manually scrutinized on the Binary Alignment Map (BAM) files through Integrative Genomics Viewer (IGV) version 2.3.25 (Thorvaldsdóttir et al. 2013).

#### 1.5 Copy number aberration (CNA) calls

Copy number changes were assessed using Nexus Copy Number Discovery Edition Version 8.0 (BioDiscovery, Inc., El Segundo, CA). Nexus NGS functionality (BAM ng CGH) with the FASST2 Segmentation algorithm was used to make copy number calls (a Circular Binary Segmentation/Hidden Markov Model approach). The significance threshold for segmentation was set at $5X10^{-6}$, also requiring a minimum of 3 probes per segment and a maximum probe spacing of 1000 between adjacent probes before breaking a segment. The log ratio thresholds for single copy gain and single copy loss were set at $+0.2$ and $-0.2$, respectively. The log ratio thresholds for gain of 2 or more copies and for a homozygous loss were set at $+0.6$ and $-1.0$, respectively. Tumor sample BAM files were processed with corresponding normal tissue BAM files. Reference reads per CN point (window size) was set at 8000. Probes were normalized to median. Relative copy number profiles from exome sequencing data were determined by normalizing tumor exome coverage to values from whole blood controls. The germline exome sequences were used to obtain allele-specific copy number profiles and generating segmented copy number profiles. The GISTIC module on Nexus identifies significantly amplified or deleted regions across the genome. The amplitude of each aberration is assigned a G-score as well as a frequency of occurrence for multiple samples. False Discovery Rate q-values for the aberrant regions have a threshold of 0.15. For each significant region, a "peak region" is identified, which is the part of the aberrant region with greatest amplitude and frequency of alteration. In addition, a "wide peak" is determined using a leave-one-out algorithm to allow for errors in the boundaries in a single sample. The "wide peak" boundaries are more robust for identifying the most likely gene targets in the region. Each significantly aberrant region is also tested to determine whether it results primarily from broad events (longer than half a chromosome arm), focal events, or significant levels of both. The GISTIC module reports the genomic locations and calculated q-values for the aberrant regions. It identifies the samples that exhibit each significant amplification or deletion, and it lists genes found in each "wide peak" region.

##### 1.6 Whole transcriptome sequencing (RNA-seq)

Total RNA from 100$\mu$m sections of snap-frozen tissue was isolated using the mirVana Isolation Kit from Ambion (AM-1560). Strand specific RNA sequencing was performed on quality controlled high RIN value ($>$7) RNA samples (Bioanalyzer Agilent Technologies) before processing at the high throughput sequencing facility core at BGI Genomics Co., Ltd. (The Children's Hospital of Philadelphia, Pennsylvania, USA). In brief, 200ng of total RNA was first treated to remove the ribosomal RNA (rRNA) and then purified using the Agencourt RNA Clean XP Kit (Beckman Coulter) prior to analysis with the Agilent RNA 6000 Pico Chip to confirm rRNA removal. Next, the rRNA-depleted RNA was fragmented and converted to cDNA. Subsequent steps include end repair, addition of an 'A' overhang at the 3' end, ligation of the indexing-specific adaptor, followed by purification with Agencourt Ampure XP beads. The strand specific RNA library prepared using TruSeq (Illumina Catalogue No. RS-122-2201) was amplified and purified with Ampure XP beads. Size and yield of the barcoded libraries were assessed on the LabChip GX (Caliper), with an expected distribution around 260 base pairs. Concentration of each library was measured with real-time PCR. Pools of indexed library were then prepared for cluster generation and PE100 sequencing on Illumina HiSeq 4000.

##### 1.7 Transcriptome (RNA-seq) quantification

Using splice-aware aligner STAR (2.3.1z) (Dobin et al. 2013), RNA-seq reads ( 200MB in size) were aligned onto the human genome reference (GRCh38) and exon-exon junctions, according to the known gene model annotation from the Ensembl release 80 (http://www.ensembl.org). Apart from protein coding genes, non-coding RNA types and pseudogenes are further annotated and classified. Based on the alignment and by using gene annotation (Ensembl release 80), gene expression profiles was calculated. Only reads unique to one gene and which corresponded exactly to one gene structure, were assigned to the corresponding genes by using the python tool HTSeq (Anders et al. 2015). Normalization of read counts was conducted by R package DESeq (Anders and Huber 2010), which was designed for gene expression analysis of RNA-seq data across different samples.

##### 1.8 Identification of fusion transcripts and validation

We used the deFuse algorithm (McPherson et al. 2011) to predict rearrangements in RNA sequence libraries. The deFuse fusion transcript prediction calls were further filtered using following criteria: a fusion gene candidate: (1) must be predicted to have arisen from genome rearrangement, rather than via a readthrough event; (2) must be predicted in no more than two sequence libraries; (3) must map unambiguously on both sides of the predicted breakpoints (that is, no multi-mapping reads); (4) must not map entirely to repetitive elements; (5) must be detected in >5 reads (either split or spanning) and (6) must have at least one of the fusion partner transcript expressed.

Prioritized putative gene fusions were verified by designing PCR primers around the predicted fusion sites. Specifically, reverse transcription PCR (RT-PCR) was used to amplify the predicted fusion gene junctions from the same starting RNA material (100ng) as was used for RNA-seq. Two primers (20-22 bp nucleotides) spanning the exon boundary of fused genes were designed using Primer3 (v. 0.4.0) (Untergasser et al. 2012). PCR was performed in 20$\mu$l reactions using Q5 buffer (NEB), 0.2mM dNTPs, 0.4 $\mu$M each primer, 0.12 units Q5 High-Fidelity DNA Polymerase (NEB) and 2 $\mu$l of the RT reaction. The PCR reaction was carried out with the following program: 95$^{\circ}$C, 30 seconds, followed by 30 cycles of 95$^{\circ}$C for 10 seconds, 57$^{\circ}$C for 20 seconds and 72$^{\circ}$C for 10 seconds. Resulting PCR products, ranging in size from 150bp to 250bp, were purified using AMPure beads (Agencourt) and sequenced using Sanger sequencing to verify fusion junctions.

#### 1.9 Proteomics analysis using mass spectrometry

Fresh frozen samples dissected from tumor and adjacent normal were individually lysed in 50mM of HEPES pH 8.5, 1% SDS, and the chromatin content was degraded with benzonase. The tumor lysates were sonicated (Bioruptor Pico, Diagenode, New Jersey, USA), and disulfide bonds were reduced with DTT and capped with iodoacetamide. Proteins were cleaned up using the SP3 method (Hughes et al. 2014, 2016) (Single Pot, Solid Phase, Sample Prep), then digested overnight with trypsin in HEPES pH 8, peptide concentration determined by Nanodrop (Thermo) and adjusted to equal level. A pooled internal standard control was generated comprising of equal volumes of every sample (10$\mu$l of each of the 100$\mu$l total digests) and split into 3 equal aliquots. The labeling reactions were run as three TMT 10-plex panels (9+IS), then desalted and each panel divided into 48 fractions by reverse phase HPLC at pH 10 with an Agilent 1100 LC system. The 48 fractions were concatenated into 12 superfractions per panel by pooling every 4th fraction eluted resulting in a total 36 overall samples.

These samples were analyzed with an Orbitrap Fusion Tribrid Mass Spectrometer (Thermo Fisher Scientific) coupled to EasyNanoLC 1000 using a data-dependent method with synchronous precursor selection MS3 scanning for TMT tags. A short description follows; more detailed overview is in (Hughes et al. 2016). Briefly, an in house packed reverse phase column run with a 2 hour low pH acetonitrile gradient (5-40% with 0.1% formic acid) was used to separate and introduce peptides into the MS. Survey scans covering m/z 350-1500 were acquired in profile mode at a resolution of 120,000 (at m/z 200) with S-Lens RF Level of 60%, a maximum fill time of 50 milliseconds, and Automatic Gain Control (AGC) target of 4x$10^5$. For MS2, monoisotopic precursor selection was enabled with triggering charge state limited to 2-5, threshold 5x$10^3$ and 10 ppm dynamic exclusion for 60 seconds. Centroided MS2 scans were acquired in in the ion trap in Rapid mode after CID fragmentation with a maximum fill time of 20 milliseconds and 1 m/z isolation quadrupole isolation window, c ollision energy of 30%, activation Q of 0.25, injection for all available parallelizable time turned ON, and an AGC target value of 1x$10^4$. For MS3, fragment ions were isolated from a 400-1200 m/z precursor range, ion exclusion of 20 m/z low and 5 m/z high, isobaric tag loss exclusion for TMT, with a top 10 precursor selection. Acquisition was in profile mode with the Orbitrap after HCD fragmentation (NCE 60%) with a maximum fill time of 90 milliseconds, 50,000 m/z resolution, 120-750 m/z scan range, an AGC target value of 1x$10^5$, and all available parallelizable ON. The total allowable cycle time was set to 4 seconds.

##### 1.10 Peptide identification and protein quantification

Qualitative and quantitative proteomics analysis was done using ProteomeDiscoverer 2.1.1.21 (Thermo Fisher Scientific). To maintain consistency with transcriptome annotation, we used Ensembl GRCh38.87 human reference proteome sequence database for proteome annotation. Sequest HT 1.3 was used for Peptide Spectral Matches (PSM), with parameters specified as trypsin enzyme, two missed cleavages allowed, minimum peptide length of 6, precursor mass tolerance 10 ppm, and a fragment mass tolerance of 0.6 Da. We allowed up to 4 variable modifications per peptide from the following categories: acetylation at protein terminus, methionine oxidation, and TMT label at N-terminal residues and the side chains of lysine residues. In addition, carbamidomethylation of cysteine was set as a fixed modification. PSM results were filtered using q-value cut off of 0.05 to control for FDR determined by Percolator. Identified proteins from both high and medium-confidence level after FDR-filtering were included in the final stage to provide protein identification and quantification results. Reporter ions from MS3 scans were quantified with an integration tolerance of 20ppm with the most confident centroid. Proteins were further filtered to include only those found with minimum one peak in all samples. Proteome Discoverer processed data was exported for further statistical analysis.

##### 1.11 Mutational signature analysis

We used deconstructSigs (Rosenthal et al. 2016), a multiple regression approach to statistically quantify the contribution of mutational signature for each tumor. The 30 mutational signatures were obtained from the COSMIC mutational signature database (Alexandrov et al. 2015) (https://cancer.sanger.ac.uk/cosmic/signatures). Both silent and non-silent somatic mutations were used together to obtain the mutational signatures. In brief, deconstructSigs attempts to recreate the mutational pattern using the trinucleotide mutation context from the input sample that closely resembles each of the 30 mutational signatures from COSMIC mutational signature database. In this process, each mutational signature is assigned a weight normalize between 0 to 1 indicating its contribution (See Rosenthal et al. 2016 for more details). Only those mutational signatures with a weight more than 0.06 were considered for analysis. We further combined (or grouped) these 30 mutational signatures based on the similarity of their mutational features or functional associations.

- Mutation signatures 3, 6, 15, 21, and 26 are related to DNA mismatch repair, so they were grouped together.
- Mutation signatures 5, 12, and 16 exhibits strong transcriptional strand bias for T>C mutations, so they were grouped together.
- Mutational signatures 24 and 29 were grouped together because both exhibits transcriptional strand bias for C>A mutations indicating guanine damage that is most likely repaired by transcription-coupled nucleotide excision repair.

##### 1.12 Prioritization of driver genes using HIT'nDRIVE

Non-silent somatic mutation calls, CNA gain or loss, and gene-fusion calls were collapsed in gene-patient alteration matrix with binary labels. Gene-expression values were used to derive expression-outlier gene-patient outlier matrix using GESD test. STRING ver10 (Szklarczyk et al. 2014) protein-interaction network was used to compute pairwise influence value between the nodes in the interaction network. We integrated these genome and transcriptome data using HIT'nDRIVE algorithm (Shrestha et al. 2017). Following parameters were used: $\alpha$=0.9, $\beta$=0.6, and $\gamma$=0.8. We used IBM-CPLEX as the ILP solver.

##### 1.13 Consensus clustering

We used ConsensusClusterPlus (Wilkerson and Hayes 2010) R-package to perform consensus clustering. We used the following parameters: maximum cluster number to evaluate: 10, number of subsamples: 10000, proportion of items to sample: 0.8, proportion of features to sample: 1, cluster algorithm: hierarchical, distance: pearson.

##### 1.14 Protein attenuation analysis

For every gene/protein profiled for CNA (segment mean), RNA-seq (normalized log2 expression), and MS (normalized log2 expression), we performed the following analysis. For every gene/protein, the Pearson correlation coefficients were calculated for CNA-mRNA expression ($\mathrm{R}_{\mathrm{CNA:mRNA}}$) and CNA-protein expression ($\mathrm{R}_{\mathrm{CNA:protein}}$). The $75^{\mathrm{th}}$ percentile of the difference between the above two correlation coefficients i.e. $R_{\mathrm{diff}} = R_{\mathrm{CNA:mRNA}} - R_{\mathrm{CNA:protein}}$ was found to be approximately 0.45. Therefore those proteins with $R_{\mathrm{diff}} \geq 0.45$ were considered as attenuated proteins.

##### 1.15 Differential expression analysis

To identify differentially expressed genes and proteins between $BAP1^{\mathrm{intact}}$ and $BAP1^{\mathrm{del}}$, we performed Wilcoxon signed-rank test using mRNA and protein expression data independently. We selected genes and proteins with $p$-value $\leq 0.05$ as the significantly differentially expressed genes and proteins. Among these we selected top-500 significantly differentially expressed genes and proteins for further analysis.

##### 1.16 Pathway enrichment analysis

The selected set of genes were tested for enrichment against gene sets of pathways present in Molecular Signature Database (MSigDB) v6.0 (Subramanian et al. 2005) A hypergeometric test based gene set enrichment analysis was used for this purpose (https://github.com/raunakms/GSEAFisher). A cut-off threshold of FDR $< 0.01$ was used to obtain the significantly enriched pathways. Only pathways that are enriched with at least three differentially expressed genes were considered for further analysis. To calculate the pathway activity score, the expression dataset was transformed into standard normal distribution using 'inverse normal transformation' method. This step is necessary for fair comparison between the expression-values of different genes. For each sample, the pathway activity score is the mean expression level of the differentially expressed genes linked to the enriched pathway.

##### 1.17 Stromal and immune score

We used two sets of 141 genes (one each for stromal and immune gene signatures) as described in (Yoshihara et al. 2013). We used 'inverse normal transformation' method to transform the distribution of expression data into the standard normal distribution. The stromal and immune scores were calculated, for each sample, using the summation of standard normal deviates of each gene in the given set.

##### 1.18 Enumeration of tissue-resident immune cell types using mRNA expression profiles

CIBERSORT algorithm (Newman et al. 2015) was applied to the RNA-seq gene-expression data to estimate the proportions of 22 immune cell types (B cells naive, B cells memory, Plasma cells, T cells CD8, T cells CD4 naive, T cells CD4 memory resting, T cells CD4 memory activated, T cells follicular helper, T cells gamma delta, T cells regulatory (Tregs), NK cells resting, NK cells activated, Monocytes, Macrophages M0, Macrophages M1, Macrophages M2, Dendritic cells resting, Dendritic cells activated, Mast cells resting, Mast cells activated, Eosinophils, and Neutrophils) using LM22 dataset provided by CIBERSORT platform. Genes not expressed in any of the PeM tumor samples were removed from the LM22 dataset. The analysis was performed using 1000 permutation. The 22 immune cell types were later aggregated into 9 distinct groups.

##### 1.19 External datasets

TCGA datasets for 16 different cancer-types used in this study were downloaded from the National Cancer Institute-Genomic Data Commons (NCI-GDC; https://portal.gdc.cancer.gov/) on February 2017. For somatic mutation data, non-silent variant calls that were identified by at least three out of four different tools (MUSE, MuTect2, SomaticSniper and VArScan2) were considered. CNA segmented data were further processed using Nexus Copy Number Discovery Edition Version 9.0 (BioDiscovery, Inc., El Segundo, CA) to identify aberrant regions in the genome. In case of the RNA-seq expression data, HTSeq-FPKM-UQ normalized data were used. AACR Project GENIE dataset was downloaded from (SynapseID: syn7222066) on April 2017.

### 2 Supplemental Results

#### 2.1 Landscape of somatic mutations in PeM

To investigate the heterogeneity of somatic gene mutations in VPC-PeM, we performed high-coverage exome sequencing (Ion Proton Hi-Q) of 19 tumors and 16 matched normal samples. We achieved a mean coverage of 180x for cancerous samples and 96x for non-cancerous samples, with at least 43-77% of targeted bases having a coverage of 100x. We identified 346 unique non-silent mutations (313 of which were not previously reported in COSMIC (Forbes et al. 2017)) affecting 202 unique genes. We observed an average of 0.015 protein-coding non-silent mutations per Mb per tumor sample. Patient MESO-18 had the highest mutation burden (0.04 mutations per Mb) and MESO-11 had the least mutation burden (0.001 mutations per Mb) (Fig S5E). The non-silent mutation burden in PeM is low compared to other adult cancers including many abdominal cancers (Fig S5A), with the exception of prostate adenocarcinomas (PRAD), kidney chromophobe carcinomas (KICH), and testicular germ cell tumors (TGCT). Notably, the mutation burden in PeM was fairly similar to PM as well as pancreatic adenocarcinomas (PAAD). We also assessed the mutational process that contribute to alterations in tumors. Analysis of base-level transitions and transversions at mutated sites showed that C>T transitions were predominant in PeM (Fig S5B-D). Using the software deconstructSigs (Rosenthal et al. 2016), we found that mutational signature 1, 5, 12, and 6 were operative in PeM. Interestingly, signature 1 was often correlated with age at diagnosis, and signature 6 was associated with DNA mismatch repair and mostly found in microsatellite instable tumors (Alexandrov et al. 2013).

The *BAP1* missense mutation in MESO-18A/E resulted in a single amino-acid (AA) change in the ubiquitin carboxyl hydrolase domain keeping the rest of the amino acid chain intact. In MESO-06 and MESO-09, a *BAP1* frameshift deletion resulted in a premature stop codon and chain termination. In MESO-09 approximately 91% of *BAP1* amino acid chains were intact, but in MESO-06 only 2% of BAP1 amino acid chains were intact (Fig S6 and Fig S7). In three (15%) tumors, we identified a recurrent R396I mutation in *ZNF678* - a zinc finger protein containing zinc-coordinating DNA binding domains involved in transcriptional regulation.

We compared the mutated genes in our VPC-PeM cohort with publically available datasets (AACR Project GENIE Consortium 2017; Alakus et al. 2015; Bueno et al. 2016) of both PeM and PM (Fig S8). *BAP1* was the only mutated gene common between the three PeM cohorts. Twenty-five genes including tumor suppressors *LATS1*, *TP53*, and chromatin modifiers *SETD2* were common between at least two PeM cohorts. Many mutated genes in VPC-PeM were also previously reported in PM. *BAP1* and *SETD2* were the two mutated genes found common between VPC-PeM and all four PM cohorts.

Our analysis investigating the distribution of variant allele frequencies of the observed variants revealed highly heterogeneous landscape of peritoneal mesothelioma mutations (Fig S26). Variants with allelic frequency ¿ 0.6 in samples MESO-05, MESO-06, MESO-14, MESO-18A/E, and MESO-19 are highly likely to be clonal and rest of the variants are highly likely to be sub-clonal. However, the rest of the samples are highly heterogeneous so it is very hard to classify the variants as clonal or sub-clonal.

#### 2.2 Copy number landscape in PeM

To investigate the somatic CNA profiles of PeM, we derived copy-number calls from exome sequencing data using the software Nexus Copy Number Discovery Edition Version 8.0. We observed a total of 1,281 CNA events across all samples. On an average, 10% of the protein-coding genome was altered per PeM tumor. MESO-14 had the highest CNA burden (42%) whereas MESO-11 had the least (0.01%). Interestingly, both mutation and CNA burden in PeM was strongly correlated ($R = 0.74$) (Fig S9A).

We also compared the CNA burden in protein-coding regions of the VPC-PeM cohort with different adult cancers from TCGA project. Similar to the mutation burden, VPC-PeM tumors were observed at the lower end of the pan-cancer CNA burden spectrum. Only UCEC, PRAD, and PAAD tumors had lower median CNA burden as compared to PeM tumors (Fig S9B). CNA status and mRNA expression for around half of the genes were positively correlated ($R \geq 0.1$) and 16% of the genes had strong correlation ($R \geq 0.5$) (Fig S9D). To identify cancer genes, we compared aberrations in protein-coding genes with data from the Cancer Gene Census (CGC) (Futreal et al. 2004) (https://cancer.sanger.ac.uk/census). Intriguingly, CNA status and mRNA expression for majority of CGC genes were positively correlated.

To identify recurrent focal CNAs in PeM tumors, we used the GISTIC (Mermel et al. 2011) algorithm which yielded 5 regions of focal deletions ($q < 0.05$) including in 3p21 which is characteristic of malignant mesotheliomas (Fig S9C). Furthermore, GISTIC prioritized 8 regions of focal amplification ($q < 0.05$) which included genes such as *BRD9*, *FOXL1*, *EGFR*, and *PDGFA*. Copy-number status of these genes was also significantly correlated with their respective mRNA expression (Fig S9E). Chromosome 1 was the most aberrant region in PeM and chromosomes 13 and 18 were relatively unchanged except for MESO-14.

CNA status of genes associated with a number of key cancer pathways was observed to be different between the PeM subtypes (i.e. $BAP1^{del}$ and $BAP1^{intact}$). We observed many genes involved in chromatin remodeling, SWI/SNF complex and DNA repair pathway to be deleted in $BAP1^{del}$ tumors as compared to $BAP1^{intact}$ tumors. In contrast, we found copy-number gain of many genes in DNA repair pathways (*BRCA2*, *ATM*, *MGMT*, and *RAD51*) and the cell cycle (*MYC*, *CDK5*, *CCNB1*, and *CCND1*) in the $BAP1^{intact}$ tumors. Furthermore, PeM tumors (both $BAP1^{del}$ and $BAP1^{intact}$) harbored CNA events in carcinogenic pathways such as MAPK, PI3K, MTOR, Wnt, and Hippo pathways. Interestingly, *ESR1* copy number deletion is enriched in $BAP1^{del}$ tumors while co-amplification of *EGFR* and *BRAF* were present in three $BAP1^{intact}$ tumors. Notably, we identified copy-number loss of tumor suppressor *LATS1/2* previously associated with mesotheliomas (Bueno et al. 2016) and copy-number gain of *NF2* in one case in $BAP1^{del}$ tumors. Notably, both LATS1/2 and NF2 are key regulators of the Hippo pathway (Li et al. 2014).

Unsupervised consensus clustering of tumor samples based on copy-number segmentation mean values of the 3349 most variable genes identified four tumor sub-groups (Fig S12A-C). We observed that $BAP1^{del}$ and $BAP1^{intact}$ tumors were grouped into distinct clusters. This indicates that $BAP1^{del}$ tumors have distinct copy-number profiles from those of $BAP1^{intact}$ tumors. We identified 692 genes ($p$-value $< 0.01$, Kruskal-Wallis test) with significantly differential CNA genes segments between the clusters. These genes were mapped to eight distinct chromosome loci – 19p, 6q, 1q, and, 13q and were mostly gained in clusters 1 and 3, whereas Xq, 22q, and 7p loss were mostly in clusters 1 and 3 (Fig S12D).

Next, we compared the CNA profile of PeM with PM. We note that PeM and PM tumors displayed substantial overlap (Fig S13A-B). Both PeM and PM tumors harbored characteristic loss of chromosome segments 3p21, and 22q. Copy number loss of CDKN2A (9p21) or NF2 (22q) is observed in more than 40% of PM tumors (Bueno et al. 2016). However, in PeM, loss of CDKN2A and NF2 was observed in only in a single case (MESO-14). We identified 1670 genes (p-value ¡ 0.0005, Wilcoxon signed-rank test) with significant differential CNA segments between PeM and PM. Intriguingly, these 1670 genes mapped to eleven distinct chromosome loci 1q, 3p, 6p, 11p, 11q, 12p, 12q, and 17q that were mostly gained in PM and lost in PeM. Vice versa 22q were mostly lost in PM and gained in PeM (Fig S13A-B).

#### 2.3 Gene fusions in PeM

To identify the presence of gene fusions, we analyzed RNA-seq data in 15 PeM using deFuse algorithm (McPherson et al. 2011). Overall, 82 unique gene fusion events were identified using our filtering criteria, out of which we successfully validated 18 gene fusions using Sanger sequencing (Fig S14 and Fig S15). We observed more gene fusion events in *BAP1*$^{\text{del}}$ tumors as compared to that in *BAP1*$^{\text{intact}}$ tumors (Fig S14A-B).

Notably, *BAP1*, *SETD2*, *PBRM1*, and *KANSL1* were prioritized as a driver gene by HIT'nDRIVE on basis of gene-fusion. Fusions in these genes were mostly found in the *BAP1*$^{\text{del}}$ subtype. *MTG1-SCART1* was the most recurrent gene fusion observed in 7 cases. This was followed by *GKAP1-KIF27* and *KANSL1-ARL17B* (Fig S14C) each of which was identified in 6 different cases. Three unique fusions were present in *PBRM1*, 2 in *KANSL1*, and 1 each in *BAP1* and *SETD2* all of which are involved in chromatin remodeling process (Fig S14D-F).

#### 2.4 The global transcriptome and proteome profile of PeM

To segregate transcriptional subtypes of PeM, we performed total RNA-seq (Illumina HiSeq 4000) and its quantification of 15 PeM tumor samples for which RNA were available (RNA for remaining four tumor samples did not pass the quality control checks). We first performed principal-component analyses and unsupervised consensus clustering of all PeM tumors to determine transcriptomic patterns using genes based on variance among tumor specimens. Consensus clustering revealed two distinct transcriptome sub-groups (Fig S16A). We found *BAP1*$^{\text{intact}}$ and *BAP1*$^{\text{del}}$ have some distinct transcriptomic patterns; however, a few samples showed an overlapping pattern (Fig S17).

We performed mass spectrometry (Fusion Orbitrap LC/MS/MS) with isobaric tagging for expressed peptide identification and its corresponding protein quantification using Proteome Discoverer for processing pipeline for 16 PeM tumors and 7 matched normal tissues (matched normal samples for the remaining tumors were not available). We identified 8242 unique proteins in 23 samples analyzed (we were surprised BAP1 protein was however not detected in our MS experiment, likely due to inherent technical limitations with these samples and/or processing. Quality control analysis of in solution Hela digests also have very low BAP1 with only a single peptide observed in occasional runs). First, we analyzed global matched mRNA-protein expression correlation. Although, 58% (4715 of 8109) of proteins showed positive mRNA-protein correlation (Pearson correlation; $R \geq 0.1$), only 22.7% (1839) of the proteins were strongly correlated with their corresponding mRNA ($R \geq 0.5$) (Fig S18). Expression of 2.4% (194) of proteins strongly negatively correlated with their corresponding mRNA ($R \leq -0.5$). To analyze the proteomic pattern across PeM tumors, we performed principal-component analyses and unsupervised consensus clustering following the same procedure as described above for the transcriptome. Unlike in transcriptome profiles, the proteome profiles of BAP1 PeM tumor sub-types did not group into distinct clusters (Fig S17A, Fig S16A-B, Fig S19A).

To identify the differentially expressed genes (DEG) between *BAP1*$^{\text{intact}}$ and *BAP1*$^{\text{del}}$, we performed Wilcoxon signed-rank test using mRNA and protein expression data independently. We identified 1520 and 466 DEG (with $p$-value $< 0.05$) using mRNA and protein expression data respectively (Fig S20). However, only 53 genes were found common between the two sets of DEG. As expected, *BAP1*, *PBRM1* and *SMARCA4*, *SMARCD3* were among the top-500 DEG. Many other important cancer-related genes were differentially expressed such as *CDK20*, *HIST1H4F*,

*ERCC1*, *APOBEC3A*, *CDK11A*, *CSPG4*, *TGFB1*, *IL6*, *LAG3*, and *ATM*.

##### 2.5 Transcriptional and post-transcriptional mechanisms regulate chromatin remodeling protein-complexes

Next, we aimed to study the extent to which changes in copy number profile affects its corresponding protein expression. For this, we calculated Pearson correlation between CNA-mRNA expression and CNA-protein expression. While, copy number profile of genes, on average, have good agreement with their corresponding mRNA expression, a number of detected proteins had poor correlation with their respective gene's copy number profile. Approximately 25% (1871 of 7462) of proteins were observed to have poor correlation with their genes copy number which we here define as "attenuated proteins" (Fig S16C). Among the attenuated proteins, we identified important chromatin remodeling proteins - PBRM1, SETD2, and SMARCC1. The attenuated proteins also included cancer genes such as NF2, EGFR, APC, PIK3CA, and MAP3K4. We observed that the attenuated proteins were significantly enriched with direct protein-protein interaction partners of the UBC (hypergeometric test $p$-value: $10^{-5}$), BAP1 ($10^{-3}$), and PBRM1 ($10^{-2}$) in STRING v10 interaction network. Notably, geneset enrichment analysis revealed that attenuated proteins are more likely to form a part of a multimeric complex or bind to macromolecules (Fig S16D). These results corroborate previous findings from studies analyzing breast, ovarian and colorectal cancer datasets (Gonçalves et al. 2017). These attenuated proteins were found to be involved in mRNA processing, DNA repair pathway, cell cycle regulation, the immune system, and in carbohydrate and lipid metabolism. Strikingly, we found that DEG between the PeM subtypes are significantly associated with protein attenuation (Chi-Squared test $p$-value: $10^{-4}$ using mRNA expression DEG, $10^{-6}$ using protein expression DEG). These findings suggest that the effects of CNA are attenuated at the protein level via post-transcriptional modification.

To identify large protein complexes containing the attenuated proteins and that are variable (i.e. at least a protein subunit of the complex is differentially expressed) between PeM subtypes, we leveraged a manually curated set of core protein complexes from the CORUM database (Ruepp et al. 2009). These included many protein complexes involved in DNA conformation modification, DNA repair, transcriptional regulation, post-translational modification including ubiquitination. Using our data, we observed that the majority of the protein complexes were highly co-regulated at the protein level rather than at the mRNA level. Notably, we identified SWI/SNF (BAF and PBAF) and HDAC complex which were highly co-regulated. (Fig S16E-G). We found copy-number deletion in many subunits of SWI/SNF complex, mostly in the $BAP1^{\mathrm{del}}$ subtype. About one quarter of proteins in the BAF complex and half of proteins in PBAF were attenuated. PBRM1 was both attenuated at the protein level as well as differentially expressed between PeM subtypes. SMARCB1, and SMARCA4 were also differentially expressed between PeM subtypes in this complex (Fig S16H). We further identified a number of HDAC complex components as highly co-regulated. The complex consisted of Histone deacetylase (HDAC1/2), which regulates expression of a number of genes through chromatin remodeling. About one-third of protein subunits in the complex were attenuated at the protein level. More importantly, HDAC1, CHD4 and ZMYM2 were differentially expressed between PeM subtypes in the protein complex, and different family members of HDAC protein family were highly expressed in the $BAP1^{\mathrm{del}}$ subtype. (Fig S16I). This indicates potential use of HDAC inhibitors to suppress the tumor growth in the $BAP1^{\mathrm{del}}$ subtype. We note that both SWI/SNF and HDAC complexes interact with BAP1. Expression pattern of many subunits of these complexes were either highly correlated or highly anti-correlated with BAP1 expression. (Fig S16E-G). Although mRNA transcripts are transcribed proportional to the changes in copy-number profile of the gene, the corresponding proteins are often stabilized when in complex, and free proteins in excess are usually ubiquitinated and targeted for proteosomal degradation to maintain stoichiometry (Gonçalves et al. 2017).

##### 2.6 lncRNA and miRNA profile of PeM

Using RNA-seq quantification, we identified 6594 long noncoding RNA (lncRNA) and 171 microRNA (miRNA) expressed in at least one of the PeM cases analyzed. Using these data, we performed consensus clustering to identify different clusters of of tumor samples. We obtained four clusters each on both analysis using lncRNA (Fig S27) and miRNA (Fig S28) expression dataset. However, both dataset did not reveal any clear pattern of separation between $BAP1^{\text{del}}$ and $BAP1^{\text{intact}}$ subtypes.

Following the premise that some lncRNA and miRNA regulate the activity of immune checkpoint receptor, we were interested to identify lncRNA and miRNA that strongly correlated with immune checkpoint receptors based on RNA-seq expression. For this, we first selected those immune checkpoint receptors (*LAG3*, *ICOS*, *CD274* (*PD-L1*), *CTLA4*, and *CD80*) that are identified to be differentially expressed between $BAP1^{\text{del}}$ vs. $BAP1^{\text{intact}}$ subtypes. We then performed the Pearson correlation analysis between the above immune checkpoint receptor expression data with that of lncRNA and miRNA expression data. We then selected those lncRNA or miRNA which strongly correlated or anti-correlated (i.e. Pearson correlation coefficient: $R > 0.8$ or $R < -0.8$) with the above immune checkpoint receptor. The Table S15 summarizes the overall results. We obtained 8, 8, 1, 6, and 5 correlated or anti-correlated lncRNA respectively for *LAG3*, *ICOS*, *CD274* (*PD-L1*), *CTLA4*, and *CD80*. The resulting lncRNA are highly likely to regulate the activity of the corresponding immune checkpoint receptors or participate in the downstream signaling activity of the immune checkpoint receptors. However, all of the 171 miRNA analyzed were poorly correlated (i.e. $R < 0.8$ and $R > -0.8$) and did not make it to our filtered list.

##### 2.7 Differentially Expression Analysis: removing MESO-18A/E

To investigate if including both tumor sample from MESO-18 (i.e. MESO-18A and MESO-18E) would skew the results presented in our analysis, we performed the following analysis. We used Wilcoxon signed-rank test to identify differentially expressed genes (DEG) between $BAP1^{\text{del}}$ and $BAP1^{\text{intact}}$ mesotheliomas in the mRNA expression dataset with different sample combination as described below. We performed differential expression analysis

1. using all 15 mesothelioma samples (as presented in the main manuscript),
2. without using sample MESO-18A,
3. without using sample MESO-18E, and
4. without using samples MESO-18A and MESO-18E.

From all the above analysis, we selected top-500 DEGs with p-value $< 0.05$. We then compared the DEGs obtained using dataset

A. with all samples vs. that without MESO-18A (Fig S29A),

B. with all samples vs. that without MESO-18E (Fig S29B), and

C. with all samples vs. that without MESO-18A and MESO-18E (Fig. Fig S29C).

We observed that, there was a large overlap of the genes in each of the above comparison. When MESO-18A or MESO-18E was removed from comparison we obtained 430 and 434 DEG in common with that obtained using all samples. When both MESO-18A and MESO-18E were removed, we obtained 335 DEG in common with that obtained using all samples.

The list of DEG were ranked based on their p-values obtained using Wilcoxon signed-rank test between $BAP1^{\mathrm{del}}$ and $BAP1^{\mathrm{intact}}$ tumors. We analyzed the rank of the DEG that were identified when all samples were used but not detected when MESO-18A and/or MESO-18E were removed. The histogram on the left panel of the Fig S29 (colored in red) summarizes ranked list of DEG. We observed that these genes occupied the bottom portion of the ranked list of DEG.

We also analyzed the rank of the DEG that were detected when MESO-18A and/or MESO-18E were removed but not detected when all samples were used together. The histogram on the right panel of the Fig S29 (colored in blue) summarizes ranked list of DEG. We observed that these genes mostly occupied the bottom portion of the ranked list and very few genes occupied the top-50 or top-100 position of the ranked list of DEG.

Thus this analysis demonstrates that even when we removed MESO-18A and/or MESO-18E, the resulting set of differentially expressed genes do not change to a large degree and may have a negligible impact on our overall findings.

### 3 Supplemental Discussion

Structural alterations in PeM tumors were found to be highly heterogeneous, and occur at a lower rate as compared to most other adult solid cancers. The majority of SNVs and CNAs were typically unique to a patient. However, many of these alterations were non-randomly distributed to critical carcinogenic pathways. We observed many alterations in genes involved in chromatin remodeling, SWI/SNF complex, cell cycle and DNA repair pathway. SWI/SNF complex is an ATP-dependent chromatin remodeling complex known to harbor aberrations in almost one-fifth of all human cancers (Kadoch and Crabtree 2015). Our results show that SWI/SNF complex is differentially expressed between PeM subtypes which further regulates oncogenic and tumor suppressive pathways. Notably, we also identified another chromatin remodeling complex - HDAC complex which is differentially expressed between PeM subtypes. HDAC, known to be regulated by BAP1, is a potential therapeutic target for the *BAP1*$^{\mathrm{del}}$ PeM subtype. Recent in-vitro experiments demonstrated BAP1 loss altered sensitivity of PM as well as uveal melanoma (UM) cells to HDAC inhibition (Landreville et al. 2012; Sacco et al. 2015).

Furthermore, recently, *BAP1* loss has been defined as a distinct molecular subtype of clear cell renal cell carcinoma (ccRCC) and UM (Chen et al. 2016; Peña-Llopis et al. 2012; Robertson et al. 2017). These studies showed that, similar to *BAP1*$^{\mathrm{del}}$ PeM subtype, *BAP1*$^{\mathrm{del}}$ tumors from both ccRCC and UM also have dysregulated chromatin modifiers, impaired DNA repair pathway, and immune checkpoint receptor activation. More recent studies in ccRCC (Miao et al. 2018) and melanoma (Pan et al. 2018) demonstrated that inactivation of PBRM1 (or PBAF complex) predicts response to immune checkpoint blocking therapies. Similarly, DNA repair defects have also been shown to be predictive of response to immune checkpoint blocking therapies (Germano et al. 2017; Le et al. 2015, 2017). This strongly indicates a pan-cancer mechanism of oncogenesis shared among tumors with *BAP1* copy-number loss.

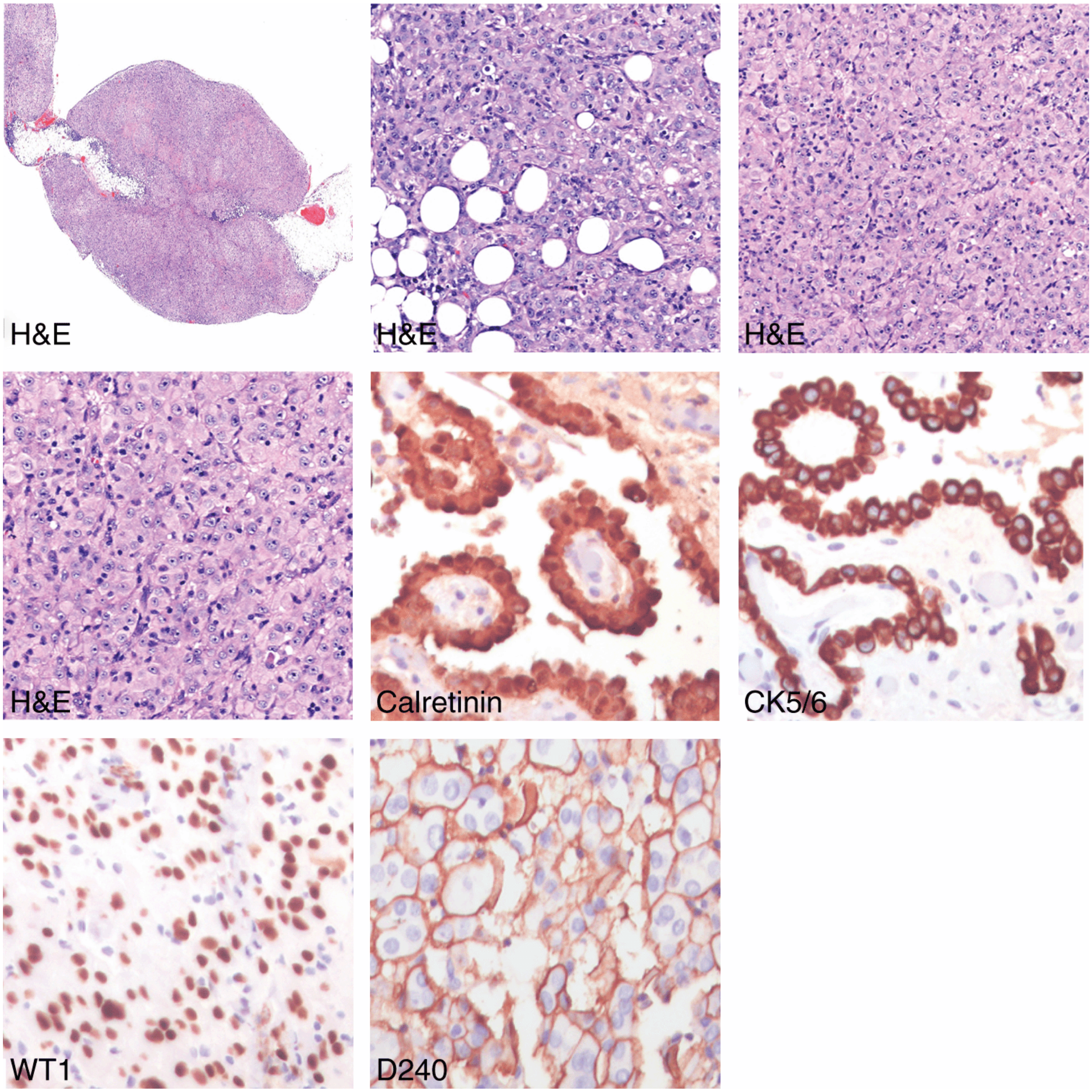

**Fig S1. Pathology of Peritoneal Mesothelioma.** Microphotographs of histological features of Epithelioid PeM from one representative patient (MESO-05). Haematoxylin and eosin (H&E) stained histological sections represent gross morphology from low to high magnification (left to right). Calretinin, CK5/6, WT1, and D240 positive stains show epithelioid subtype.

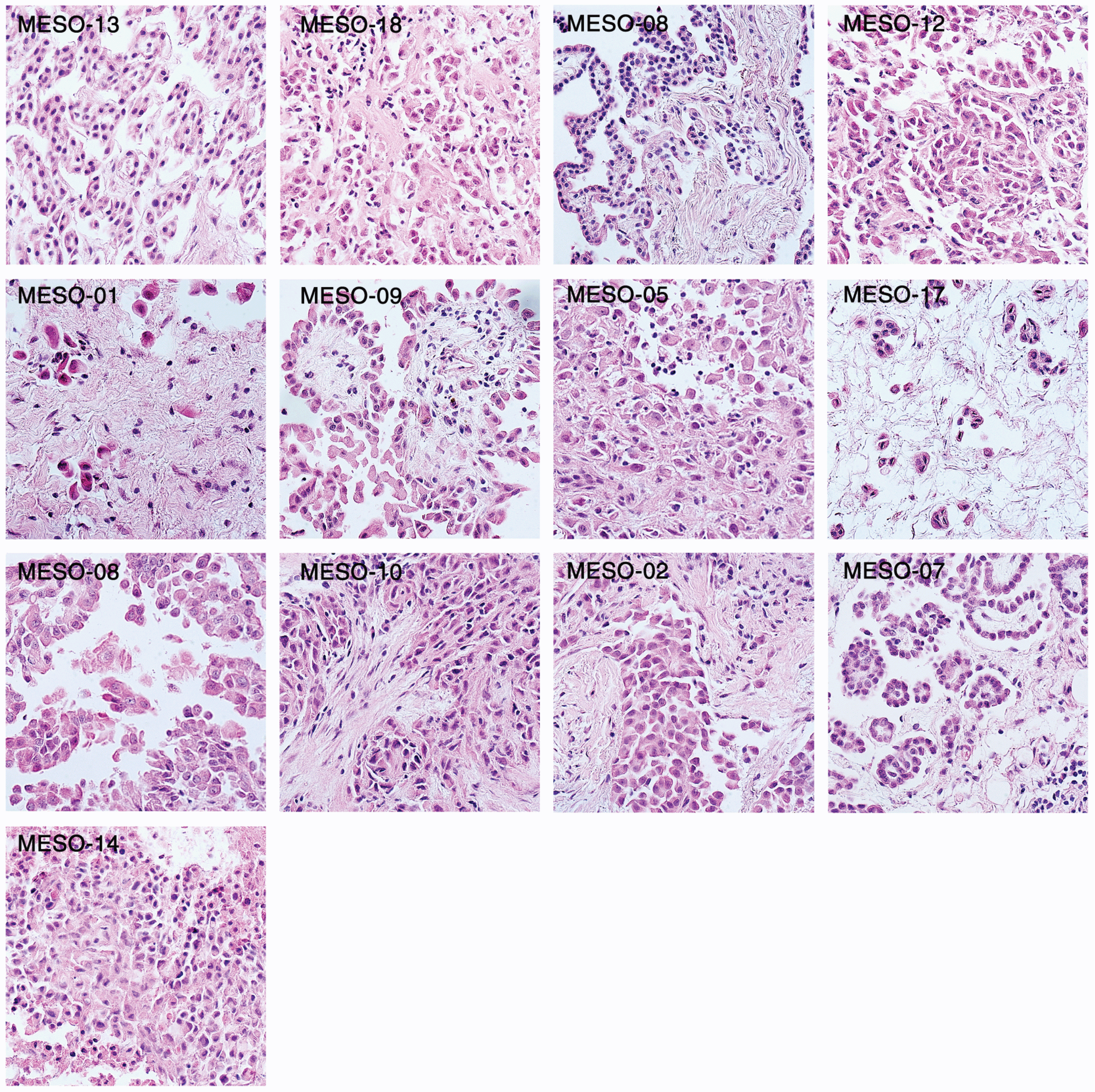

**Fig S2. Pathology of Peritoneal Mesothelioma.** Microphotographs of histological features of Epithelioid PeM stained using Haematoxylin and eosin (H&E).

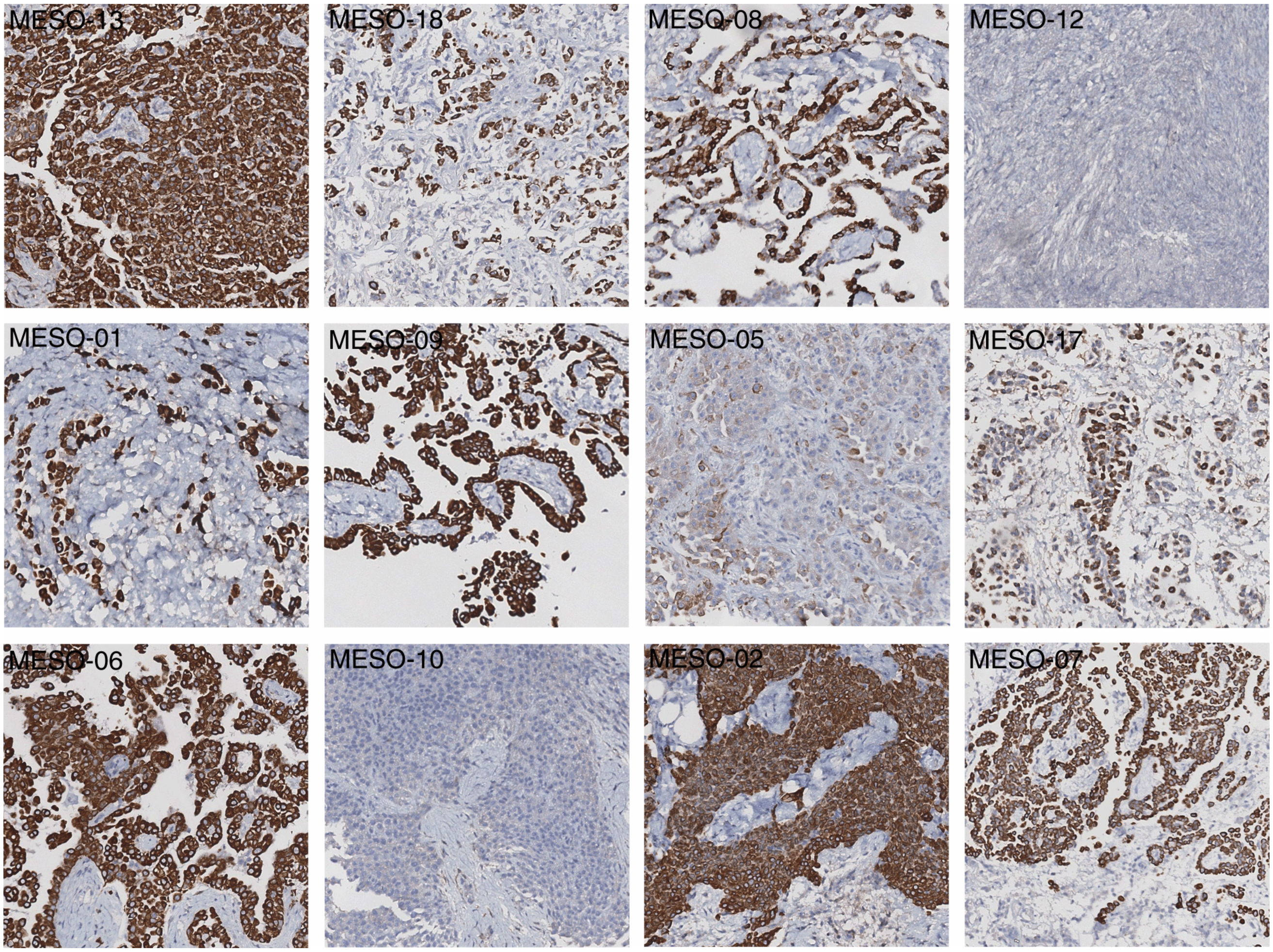

**Fig S3. TMA slides of PeM IHC stained for CK5.** The top two panel of TMA slides represents *BAP1*$^{\text{intact}}$ subtype while the bottom two panel of represents *BAP1*$^{\text{del}}$ subtype.

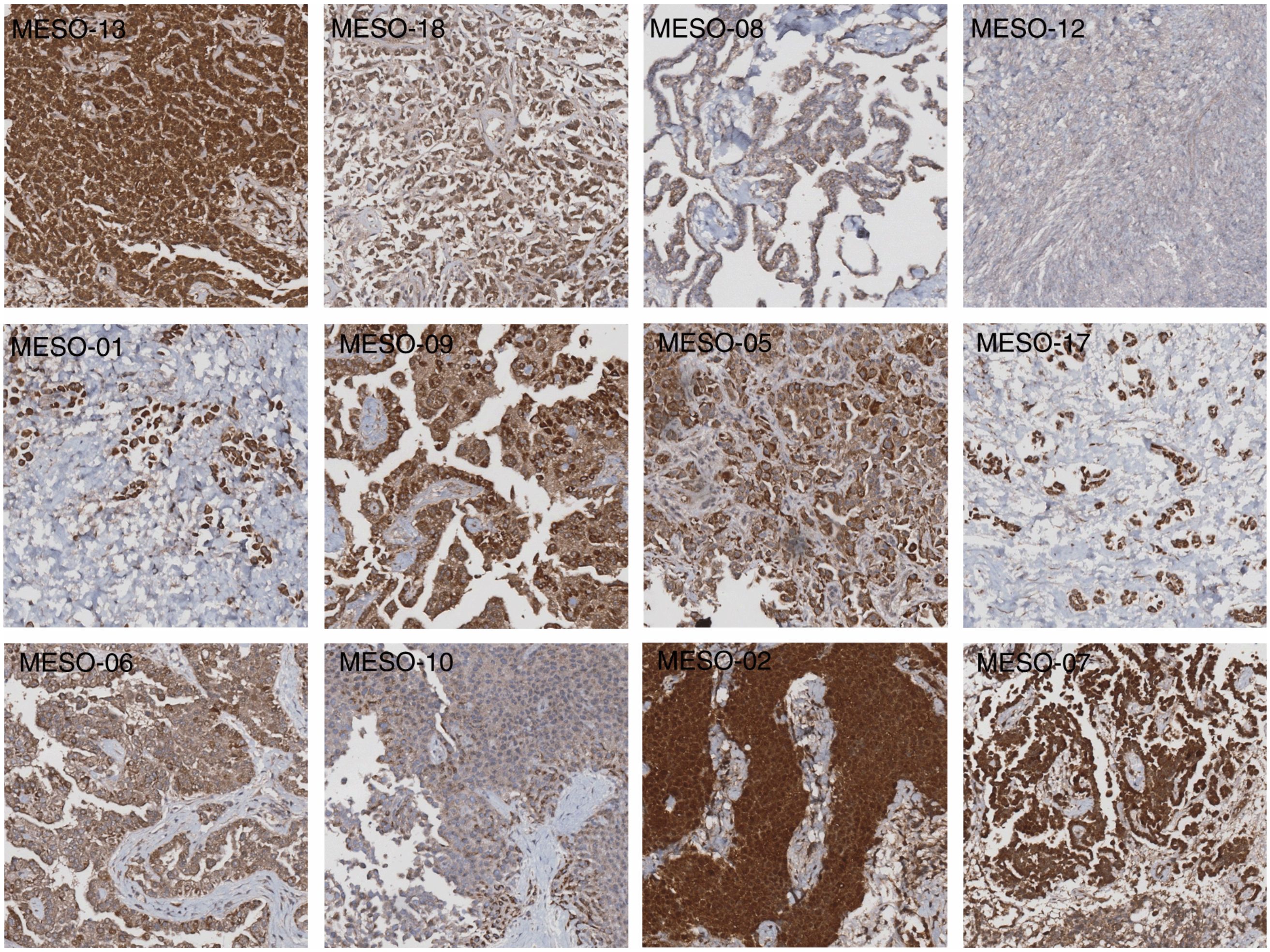

**Fig S4. TMA slides of PeM IHC stained for CALB2.** The top two panel of TMA slides represents *BAP1*$^{\text{intact}}$ subtype while the bottom two panel of represents *BAP1*$^{\text{del}}$ subtype.

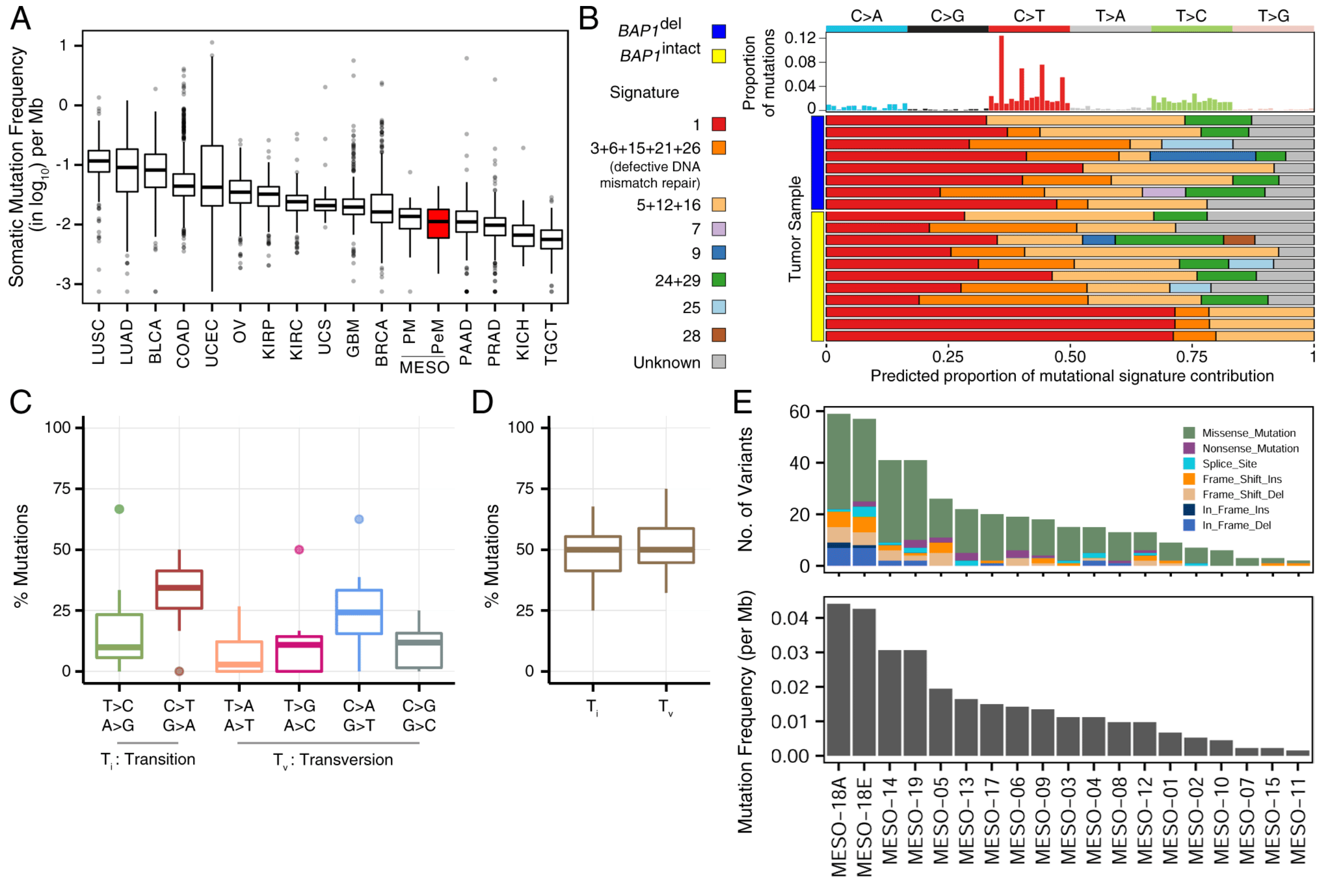

**Fig S5. Summary of non-silent somatic mutation landscape of PeM.** (A) Comparison of somatic mutation rate in protein-coding regions of PeM with different adult cancers obtained from TCGA. (B) Mutational signature present in PeM (top panel). Proportional contribution of different COSMIC mutational signature per tumor sample. (C) Percentage of six different nucleotide base-pair substitutions in all PeM samples. (D) Box-plot comparing transition ($T_i$) and transversion ($T_v$) nucleotide changes. (E) Distribution of somatic mutations in PeM grouped by variant type (top). Rate of non-silent somatic mutation per Mb in PeM (bottom). LUSC: Lung Squamous Cell Carcinoma, LUAD: Lung adenocarcinoma, BLCA: Urothelial Bladder Carcinoma, COAD: Colorectal carcinoma, UCEC: Uterine Corpus Endometrial Carcinoma, OV: Ovarian cancer, KRIP: Kidney renal papillary cell carcinoma, KIRC: Kidney Renal Clear Cell Carcinoma, UCS: Uterine Carcinosarcoma, GBM: Glioblastoma Multiforme, BRCA: Breast Invasive Carcinoma, MESO-PM: Malignant Pleural Mesothelioma, MESO-PeM: Malignant Peritoneal Mesothelioma, PAAD: Pancreatic Adenocarcinoma, PRAD: Prostate Adenocarcinoma, KICH: Kidney Chromophobe, TGCT: Testicular Germ Cell Tumor.

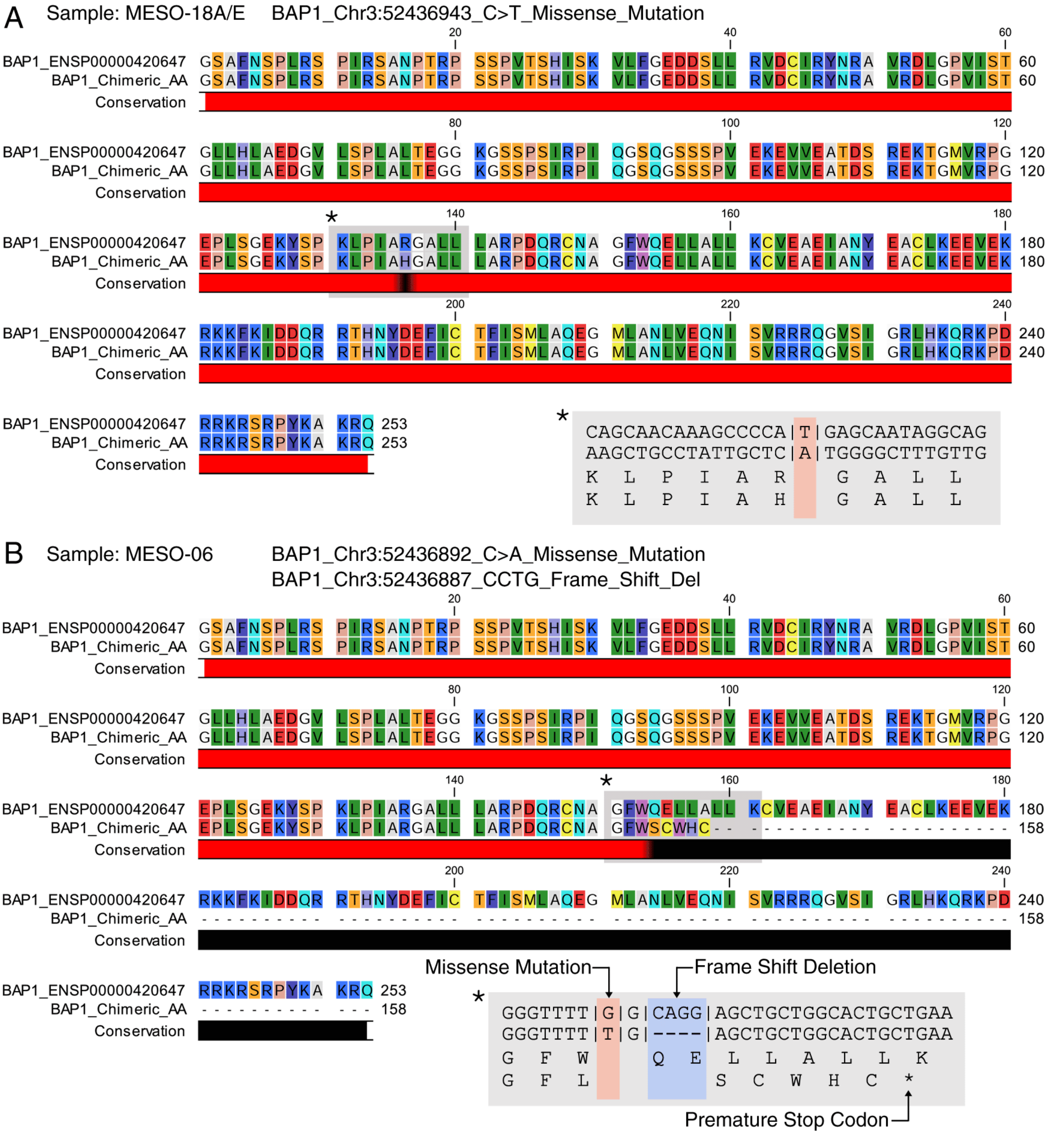

**Fig S6. Effect of *BAP1* somatic mutation on resulting amino-acid chain.** The figure shows alignment of BAP1 chimeric amino-acid sequence to the corresponding reference sequence. The changes in the nucleotide bases and the resulting amino-acid changes are highlighted on the bottom right. (A) Missense mutation in MESO-18A/E. (B) Missense mutation and frame shift deletion in MESO-06

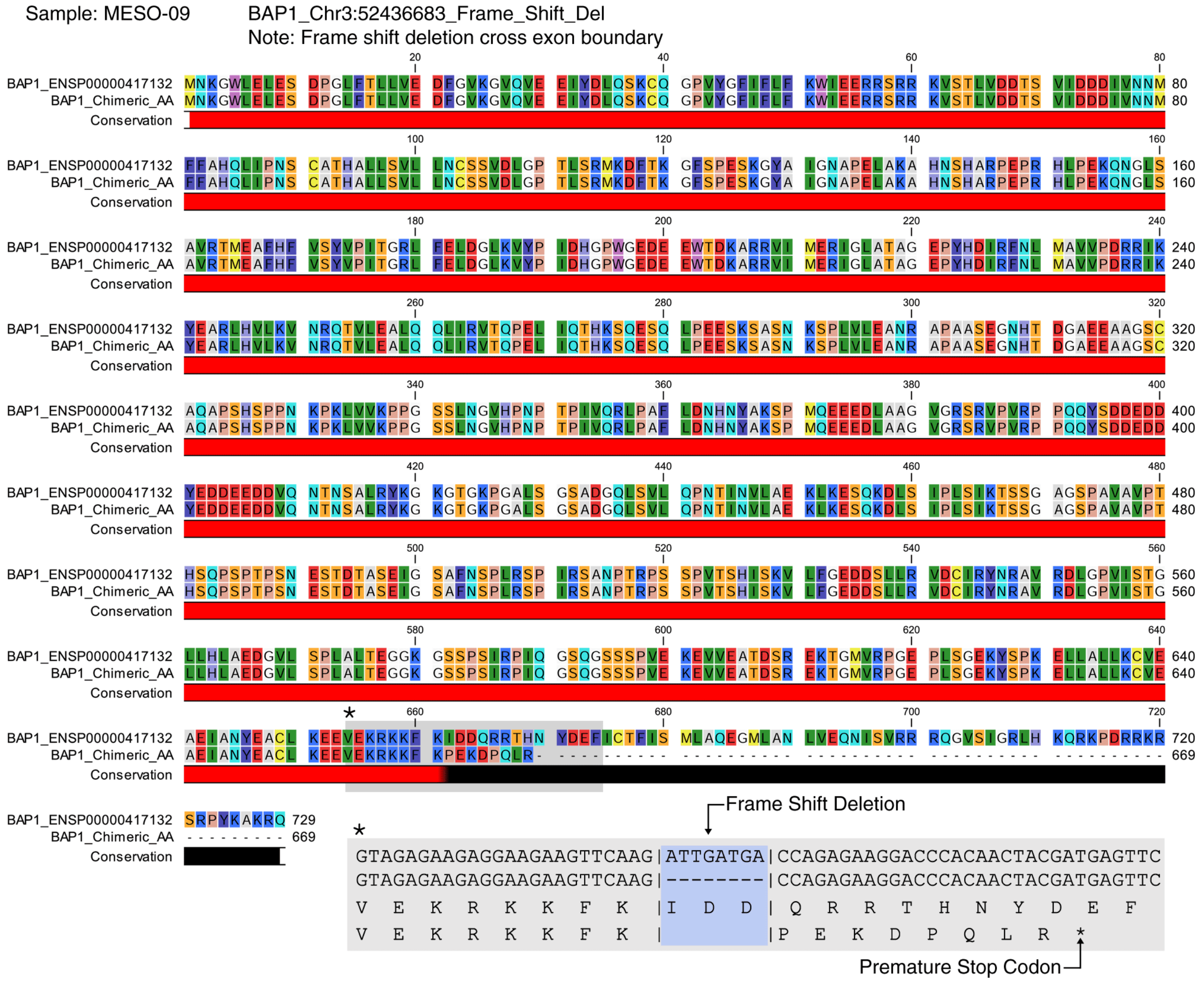

**Fig S7.** **Effect of *BAP1* somatic mutation on resulting amino-acid chain.** The figure shows alignment of BAP1 chimeric amino-acid sequence to the corresponding reference sequence. The changes in the nucleotide bases and the resulting amino-acid changes are highlighted on the bottom right.

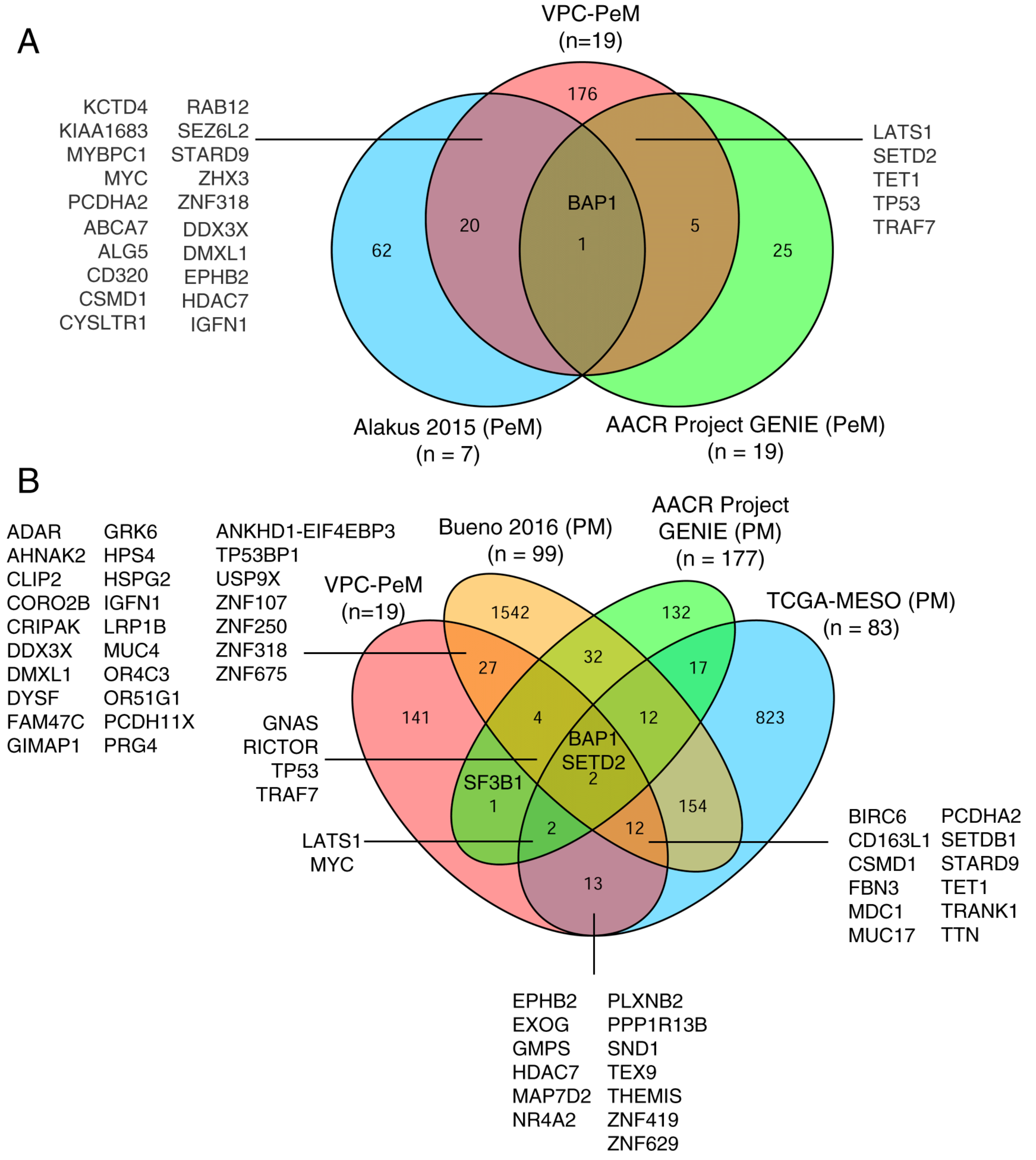

**Fig S8. Common somatic mutated genes in mesothelioma patients cohorts.** (A) Venn diagram showing mutated genes in three different PeM patient cohorts - VPC-PeM: includes tumors analyzed in this study, PeM reported in Alakus et al. 2015 and PeM reported in AACR Project GENIE. (B) Venn diagram showing common mutated genes in VPC-PeM cohort with pleural mesothelioma (PM) patient cohort. Here Bueno et al 2016, TCGA-MESO and AACR Project GENIE represent PM cohorts. The gene labels along the venn-diagram indicates the corresponding overlapped genes between different studies.

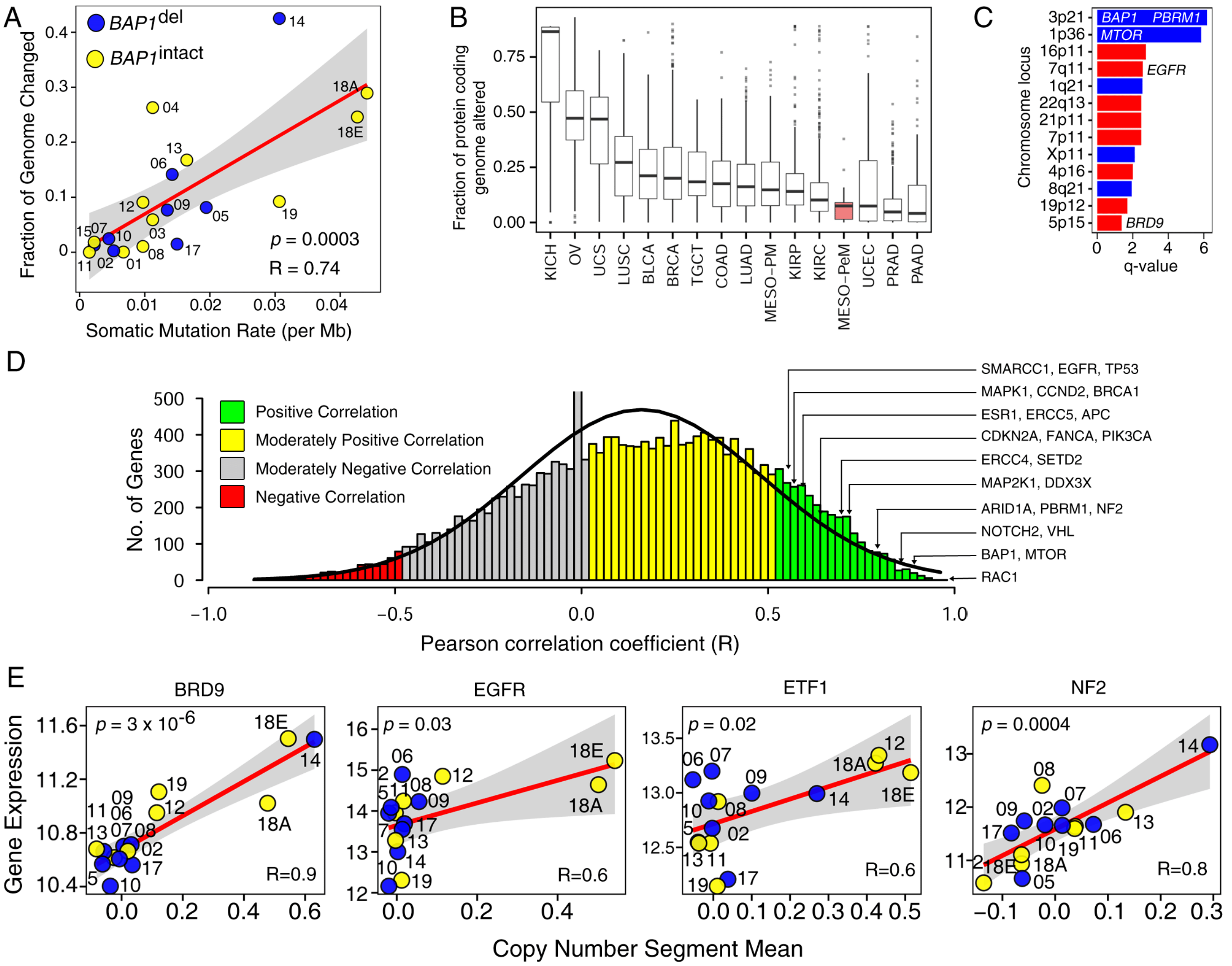

**Fig S9. Landscape of copy number aberrations in PeM.** (A)The somatic mutation rate per Mb and Fraction of copy-number genome changed were calculated for only protein-coding regions of the genome. Each dot represents a tumor sample. Blue line represents linear regression line with gray shades of 95% confidence intervals. The numbers alongside each dot represents the corresponding sample ids of the tumor. (B) Comparison of copy-number burden (considering protein-coding regions only) in PeM with respect to other adult cancers obtained from TCGA. (C) Highly aberrant genomic regions in PeM prioritized by GISTIC. (D) Correlation between copy number status and its corresponding mRNA expression. Pearson correlation coefficient (R) was used to measure the correlation between copy-number segment mean and its mRNA expression. The histogram represents the distribution R for all genes. (E) Correlation between copy-number segment mean and gene-expression of the cancer-associated genes with evidence of copy-number gain. LUSC: Lung Squamous Cell Carcinoma, LUAD: Lung adenocarcinoma, BLCA: Urothelial Bladder Carcinoma, COAD: Colorectal carcinoma, UCEC: Uterine Corpus Endometrial Carcinoma, OV: Ovarian cancer, KRIP: Kidney renal papillary cell carcinoma, KIRC: Kidney Renal Clear Cell Carcinoma, UCS: Uterine Carcinosarcoma, GBM: Glioblastoma Multiforme, BRCA: Breast Invasive Carcinoma, MESO-PM: Malignant Pleural Mesothelioma, MESO-PeM: Malignant Peritoneal Mesothelioma, PAAD: Pancreatic Adenocarcinoma, PRAD: Prostate Adenocarcinoma, KICH: Kidney Chromophobe, TGCT: Testicular Germ Cell Tumor.

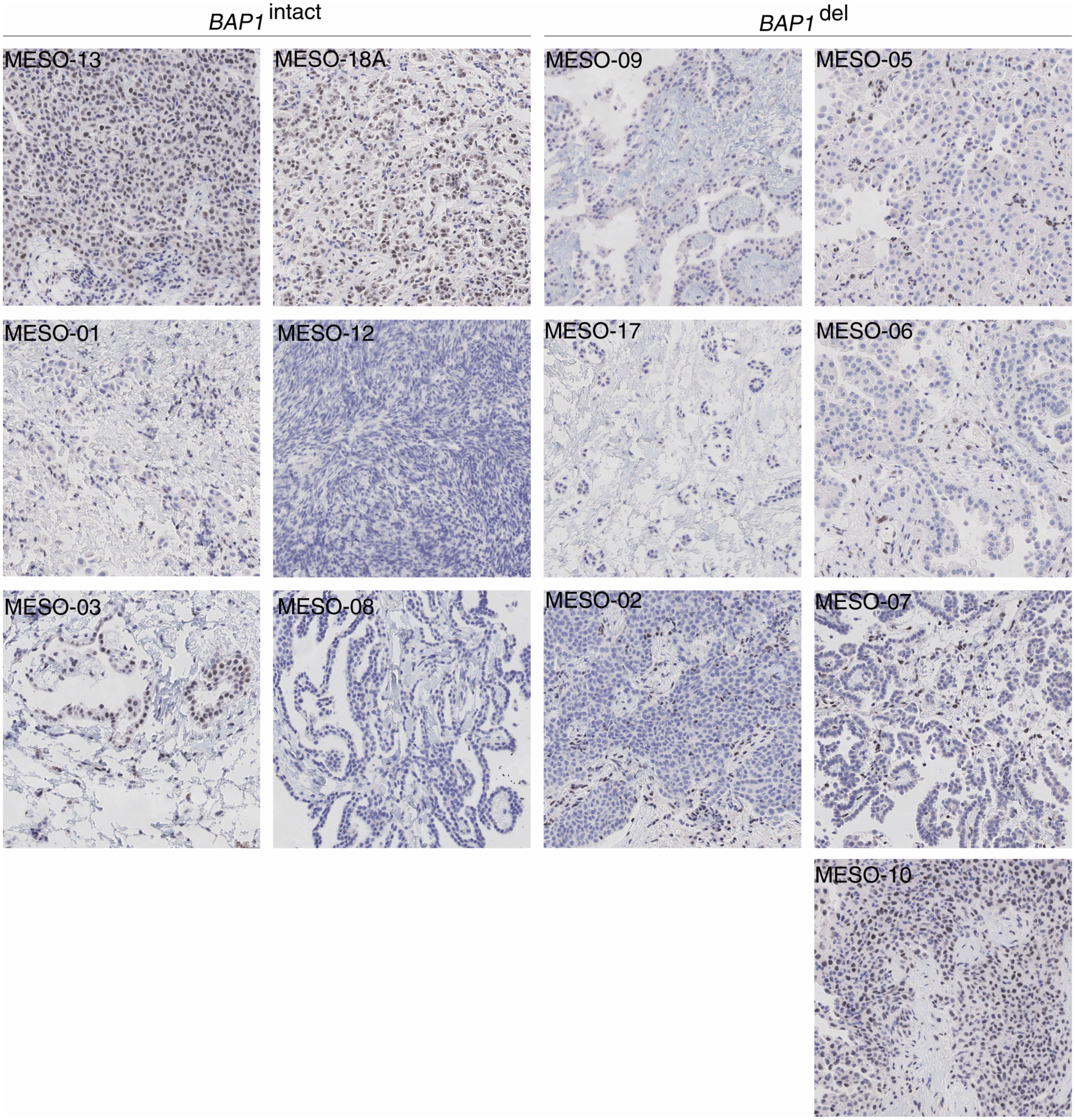

**Fig S10. TMA slides of PeM IHC stained for BAP1.** The left panel of TMA slides represents *BAP1*$^{\text{intact}}$ subtype while the right panel represents *BAP1*$^{\text{del}}$ subtype.

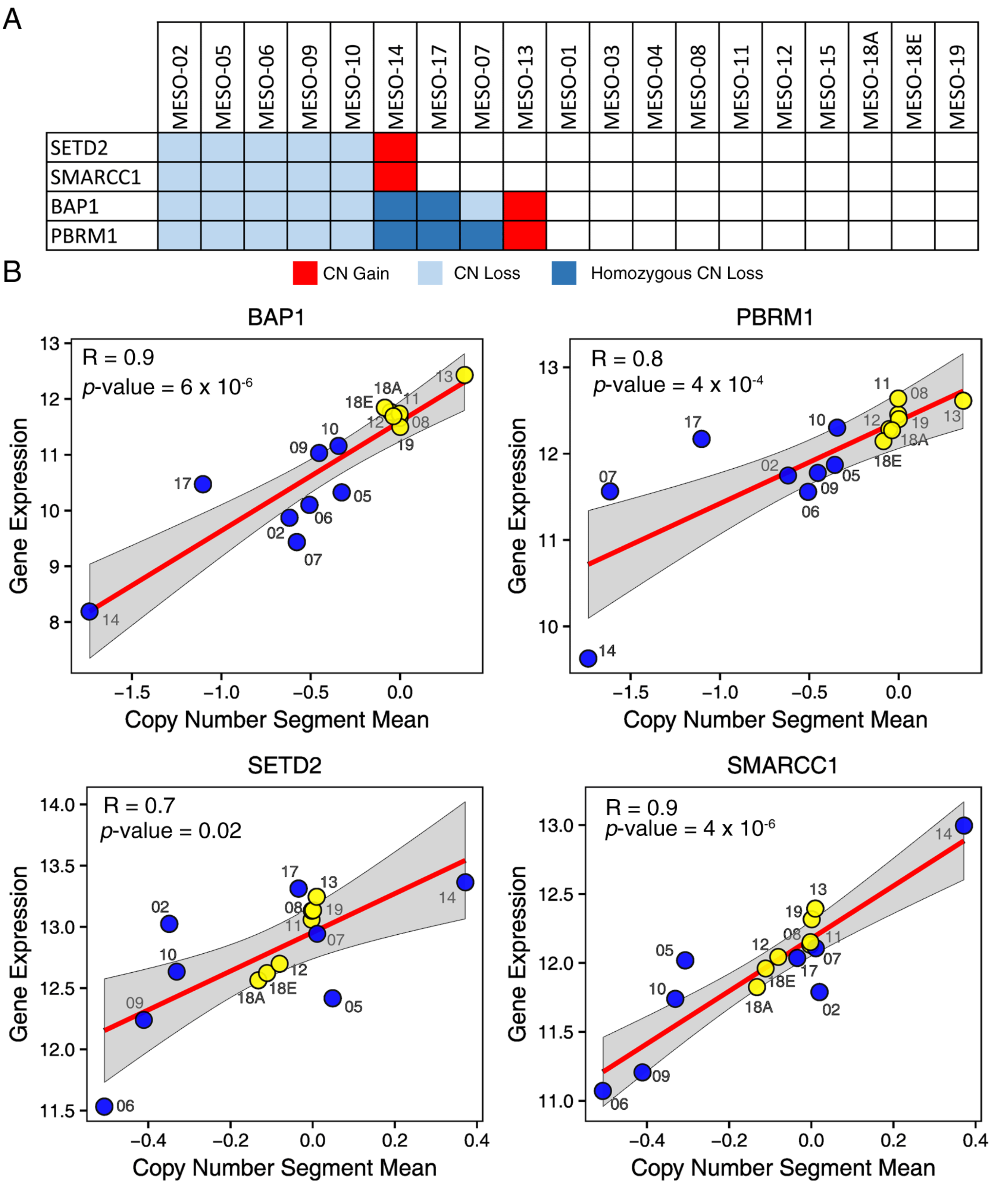

**Fig S11.** **Co-deletion of four cancer-associated genes in chromosomal region 3p21.** (A) Copy-number status of four cancer-associated in chromosomal region 3p21. (B) Correlation between copy-number segment mean and gene-expression of the above four genes. Each dot represents a tumor sample. Blue line represents linear regression line with gray shades of 95% confidence intervals. We calculated the pearson correlation coefficient (R) for each gene. The numbers alongside each dot represents the corresponding sample ids of the tumor.

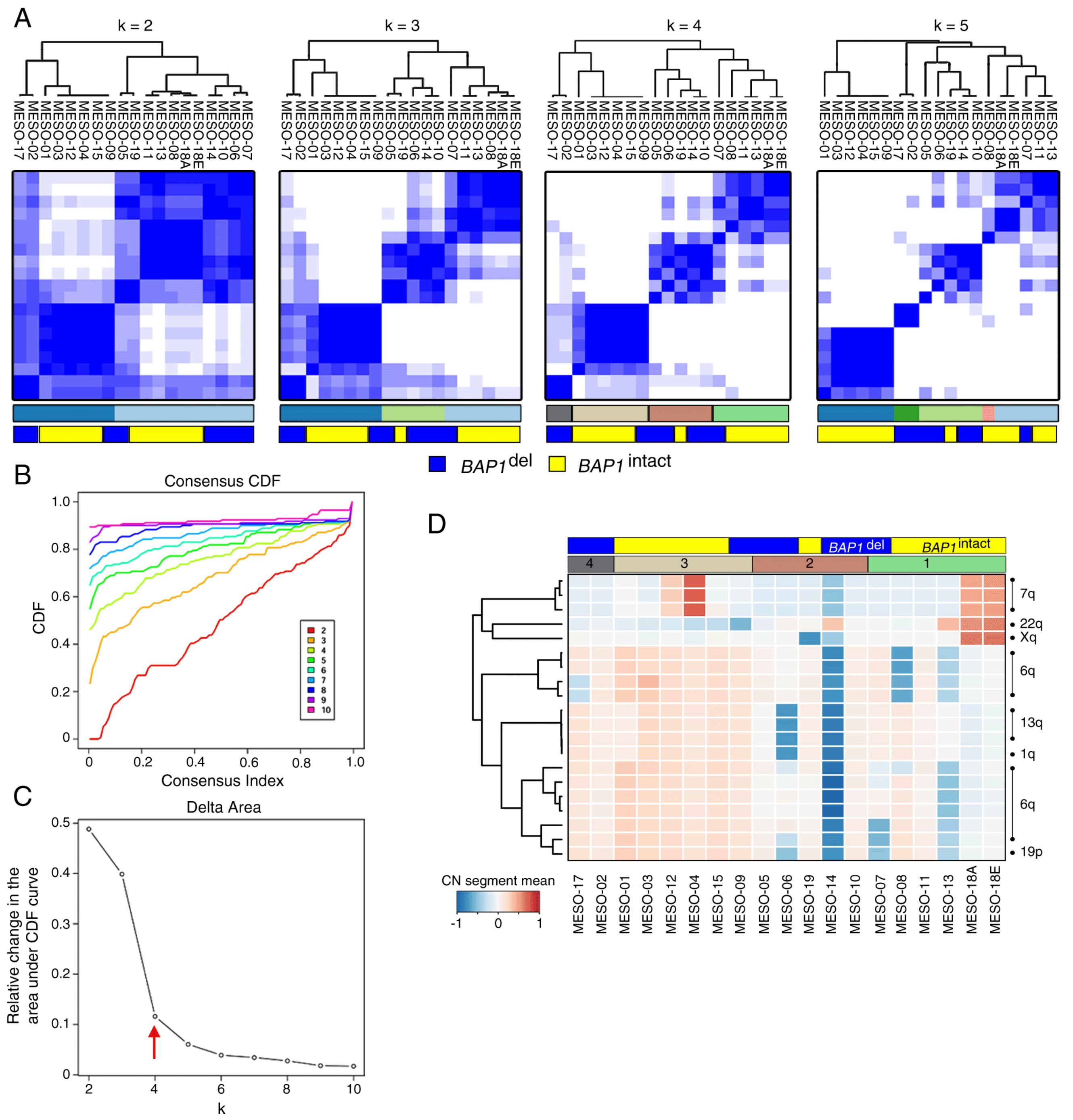

**Fig S12.** **Consensus clustering of copy-number segment mean profiles of PeM.** (A) Heatmap representing consensus clustering of copy-number segment mean profiles of PeM with different number of chosen clusters (k). (B) Consensus CDF curve and (C) Delta area curve. Optimum number of clusters (k=4) has been highlighted. (D) Differences in copy-number segments between four optimum consensus clusters.

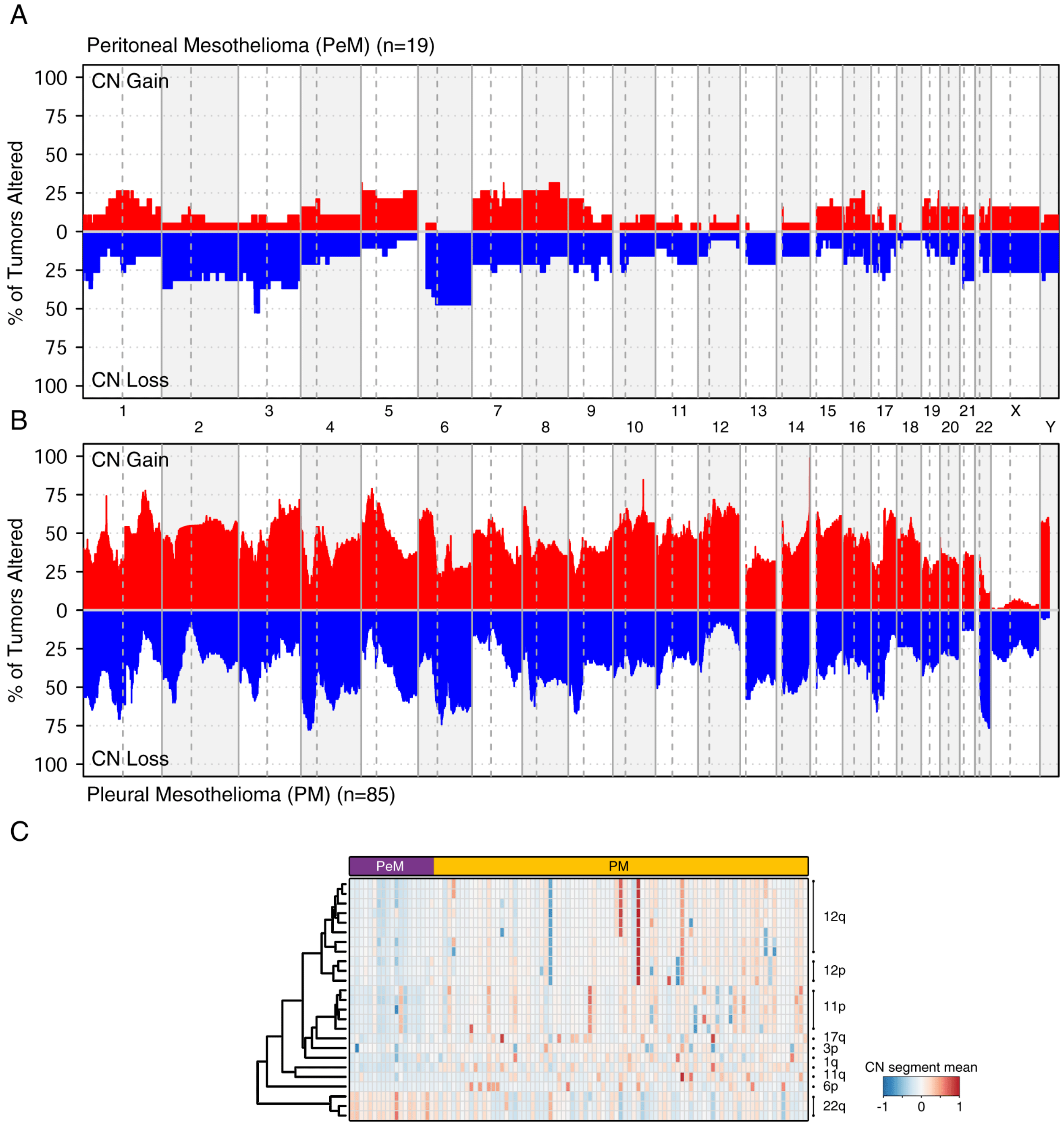

**Fig S13.** **Comparison of SCNA profile of peritoneal and pleural mesothelioma.** (A) CNA profile of peritoneal mesothelioma (PeM) and (B) CNA profile of pleural mesothelioma (PM) from TCGA. (C) Differential CNA regions between PeM and PM.

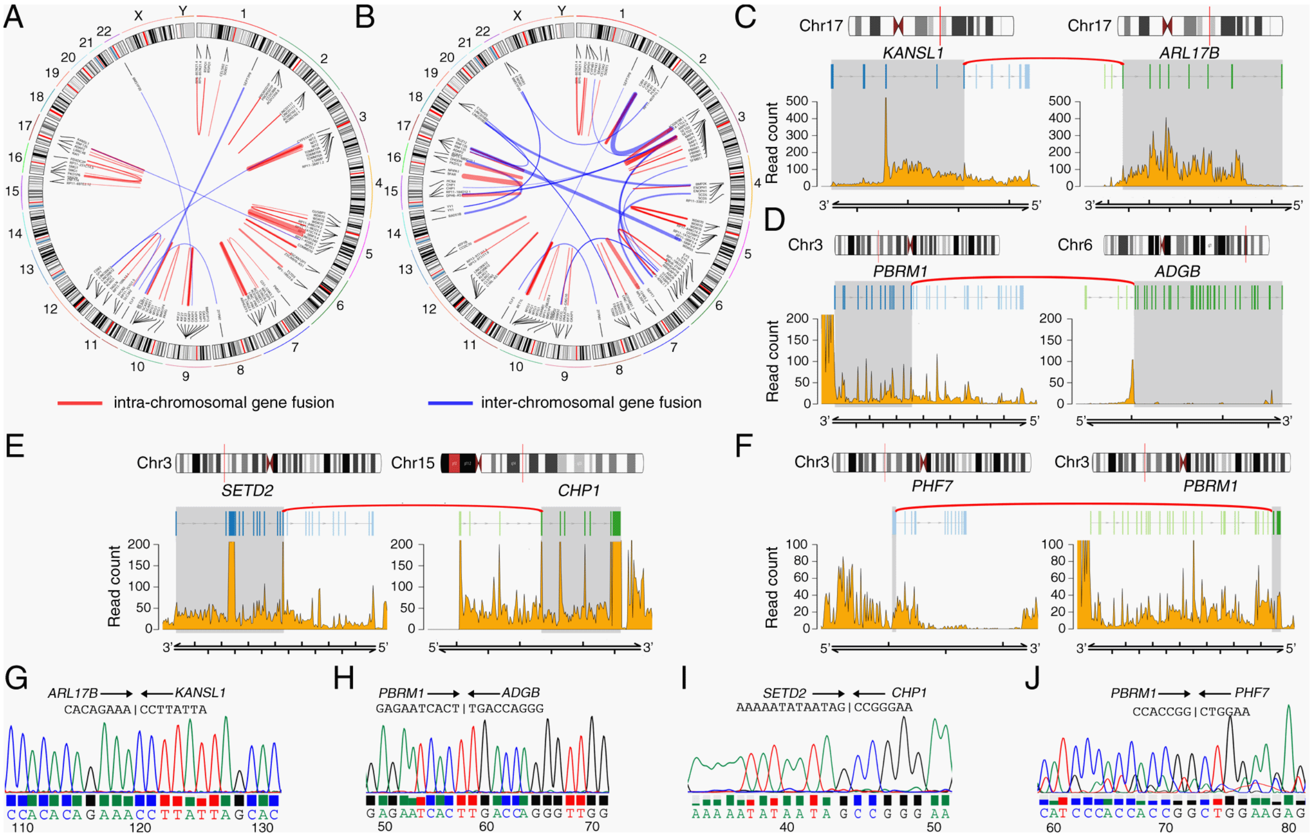

**Fig S14. Gene Fusions in PeM.** (A-B) Circos plot showing the gene fusion events identified in PeM tumors. The lines with red and blue color indicates intra-chromosomal and inter-chromosomal gene fusion events respectively. (A) *BAP1*$^{intact}$ subtype, (B) *BAP1*$^{del}$ subtype. (C-F) Selected gene fusion events identified in PeM that were validated using Sanger sequencing. The top and middle panel shows the chromosome and the transcripts involved in the gene fusion event. The bottom panel shows the RNA-seq read counts detected for the respective transcripts. (C) KANSL1-ARL17B fusion, (D) PBRM1-ADGB fusion, (E) SETD2-CHP1 fusion, and (F) PHF7-PBRM1 fusion. (G-J) The chromatogram showing the Sanger sequencing validation of the fusion-junction point.

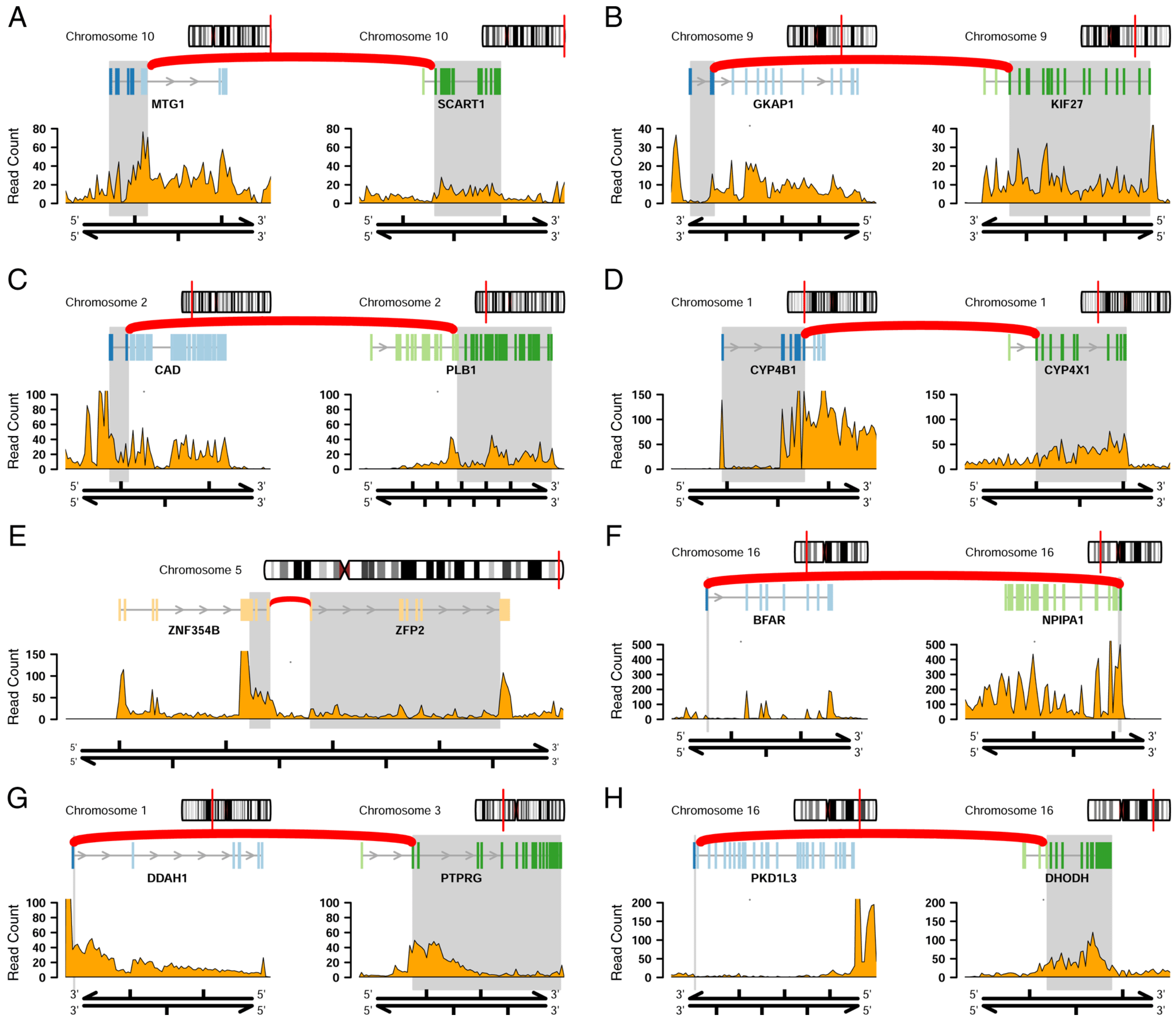

**Fig S15. Gene Fusions in PeM.** Gene fusion events identified in PeM that were validated using Sanger sequencing. The top and middle panel shows the chromosome and the transcripts involved in the gene fusion event. The bottom panel shows the RNA-seq read counts detected for the respective transcripts.

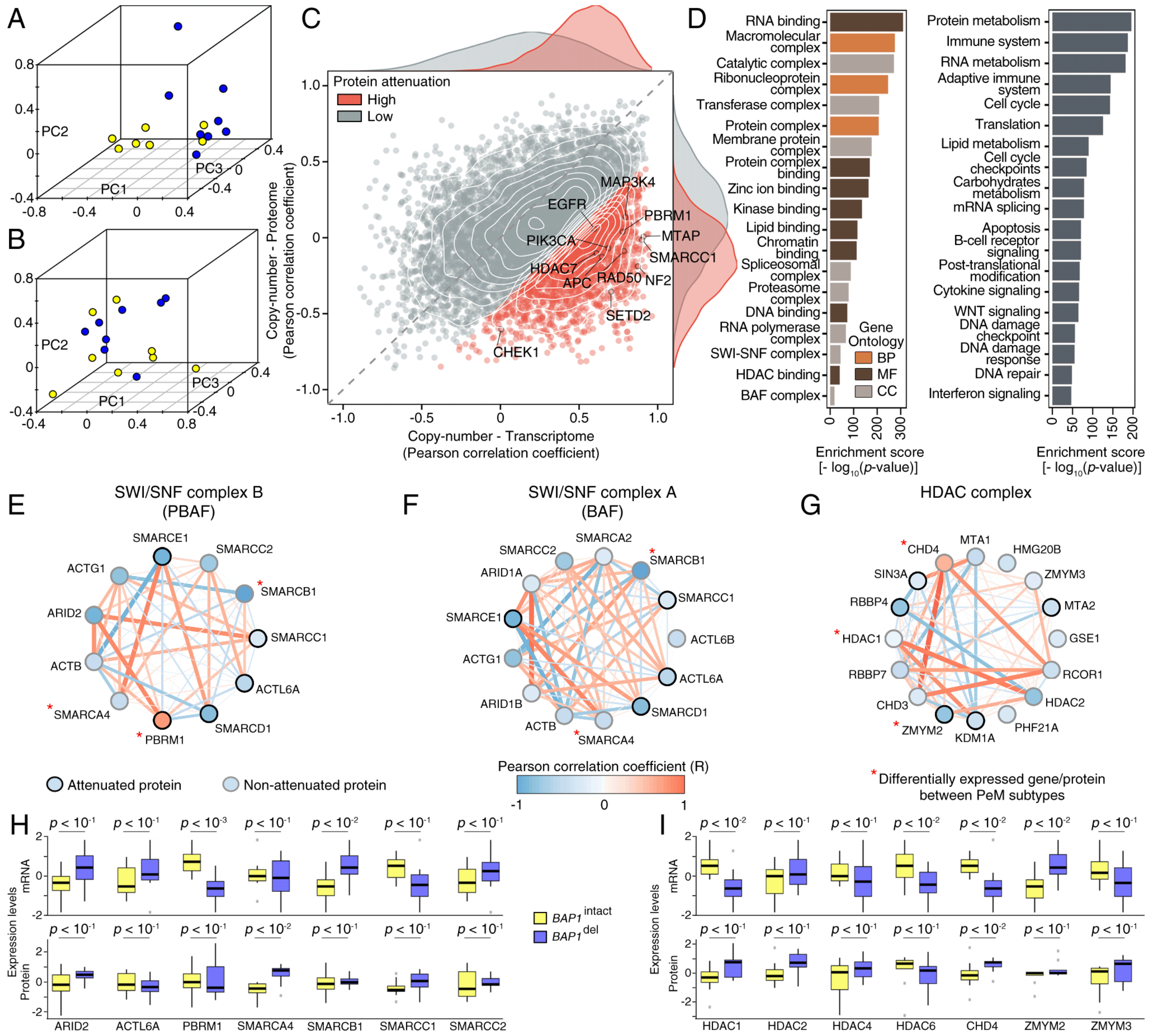

**Fig S16. Transcriptome and proteome profile of PeM.** (A-B) Principal component analysis of PeM using (A) transcriptome expression profiles and (B) proteome expression profiles. (C) Effects of CNA on transcriptome and proteome. In the scatterplot, each dot represents a gene/protein. The horizontal and vertical axes represent Pearson correlation coefficient between CNA-transcriptome and CNA-proteome respectively. Key cancer genes that undergo protein attenuation have been highlighted. (D) Geneset enrichment analysis of attenuated proteins against gene ontologies (left panel) and Reactome pathways (right panel). (E-G) CORUM core protein complexes regulated by PBRM1 and/or BAP1. The nodes represent individual protein subunit of the respective complex. The node color represents correlation of mRNA expression of respective gene with BAP1. The border color of the node indicates whether the respective protein is attenuated or not. The edge represents interaction between the protein subunits. The edge information were extracted from STRING v10 PPI network. The edge color (and edge thickness) represents correlation of protein expression between the respective interaction partners. (E) SWI/SNF complex B (PBAF), (F) SWI/SNF complex A (BAF), and (G) HDAC complex. (H-I) mRNA and protein expression level differences between PeM subtypes. (H) SWI/SNF complex and (I) HDAC complex. The expression levels are log2 transformed and mean normalized.

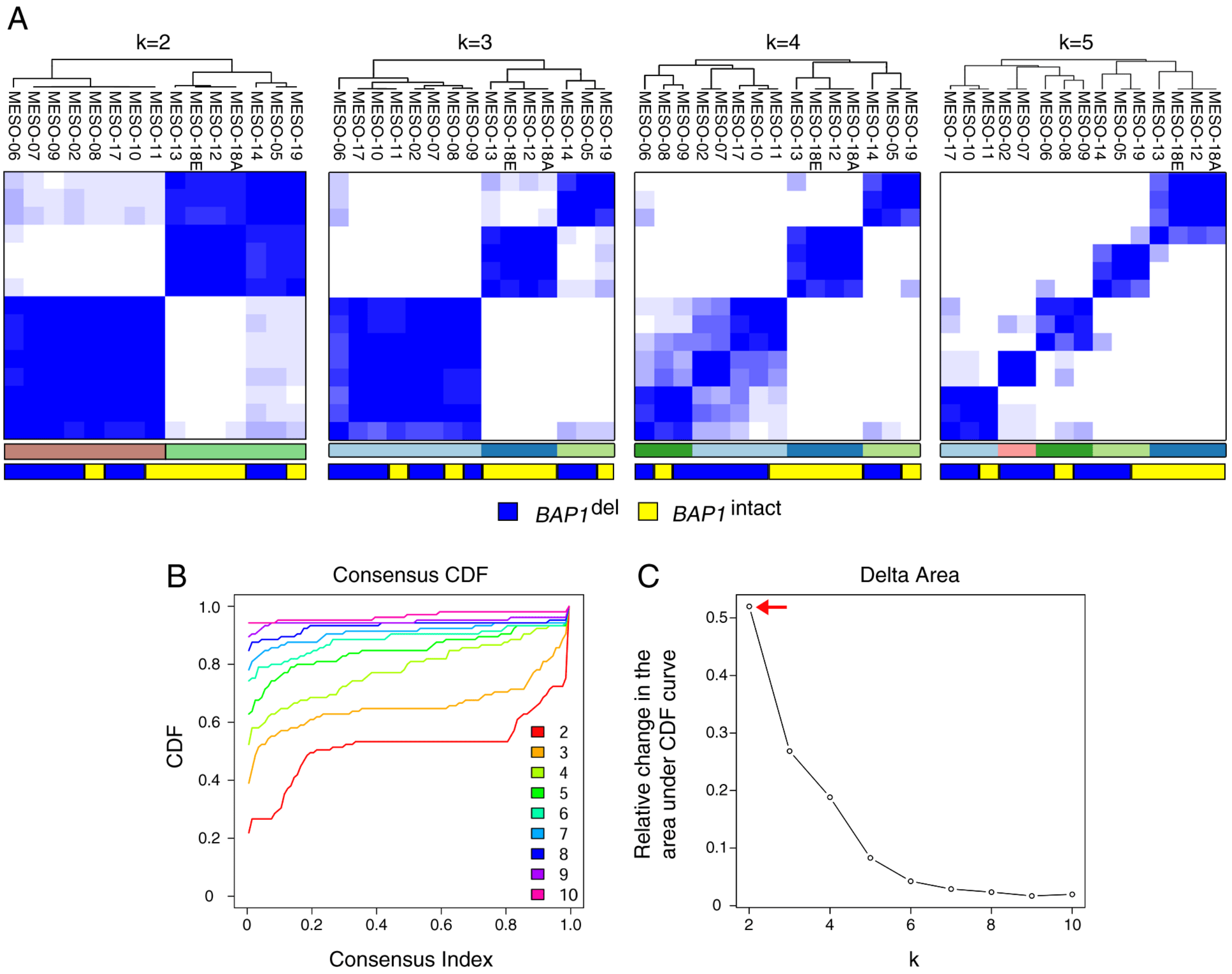

**Fig S17. Consensus clustering of mRNA expression profiles of PeM.** (A) Heatmap representing consensus clustering of mRNA expression profiles of PeM with different number of chosen clusters (k). (B) Consensus CDF curve and (C) Delta area curve. Chosen number of clusters (k=2) has been highlighted.

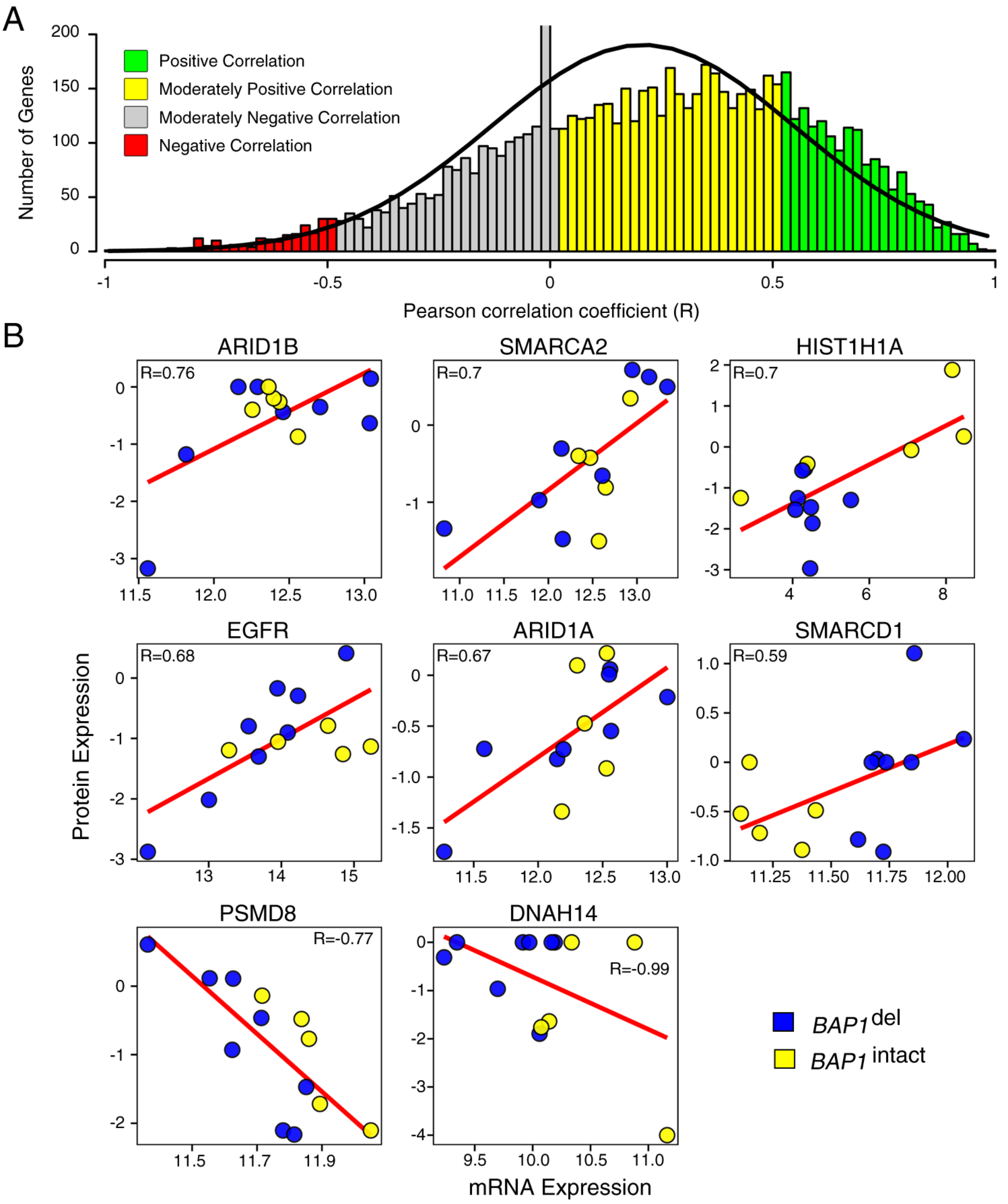

**Fig S18. Correlation between mRNA and protein expression profiles.** (A) Pearson correlation coefficient (R) was used to measure the correlation between mRNA and corresponding protein expression. The histogram represents the distribution R for all genes measured. (B) Correlation profiles of mRNA and corresponding protein expression of few selected cancer-associated genes.

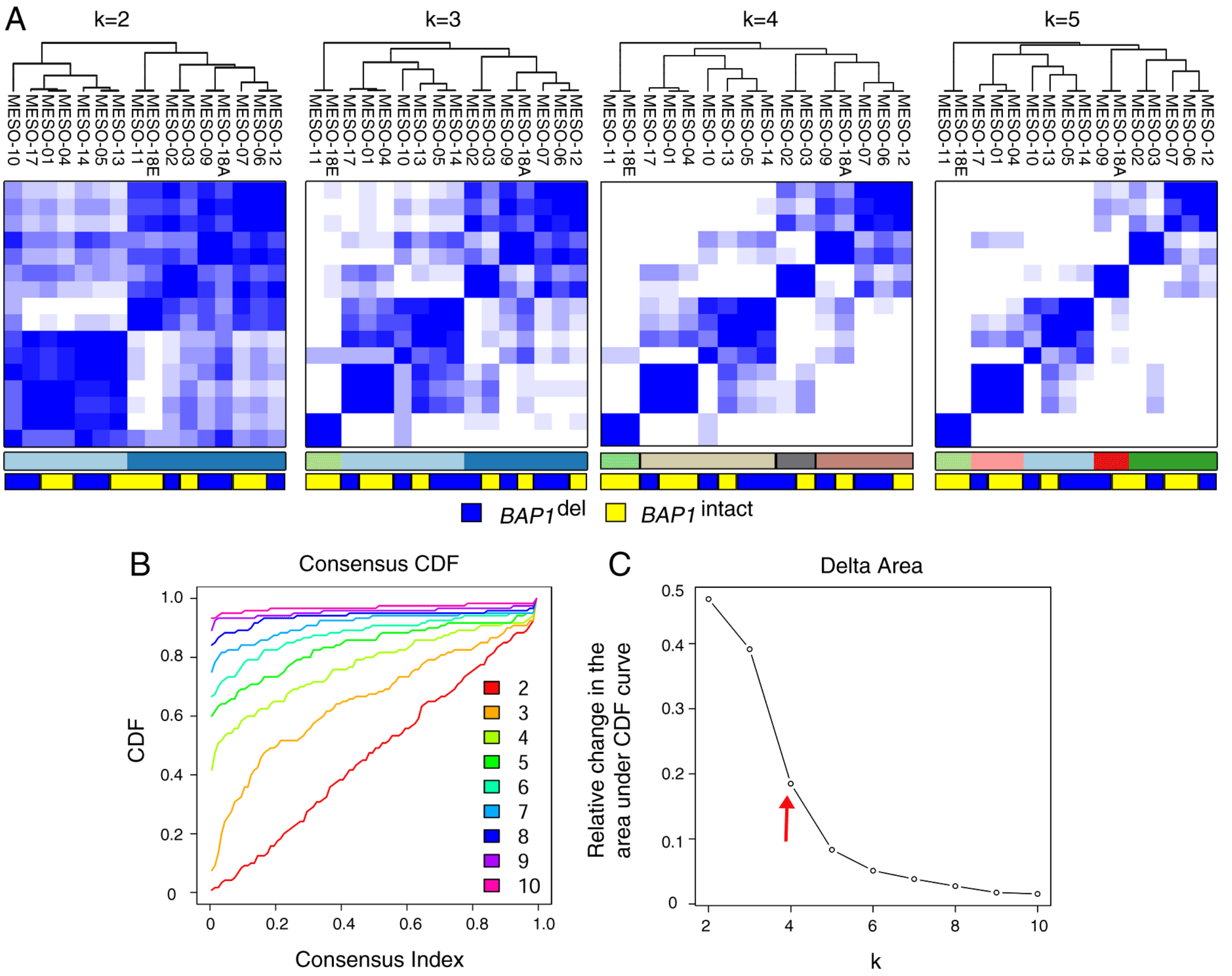

**Fig S19. Consensus clustering of mRNA expression profiles of PeM.** (A) Heatmap representing consensus clustering of mRNA expression profiles of PeM with different number of chosen clusters (k). (B) Consensus CDF curve and (C) Delta area curve. Chosen number of clusters (k=4) has been highlighted.

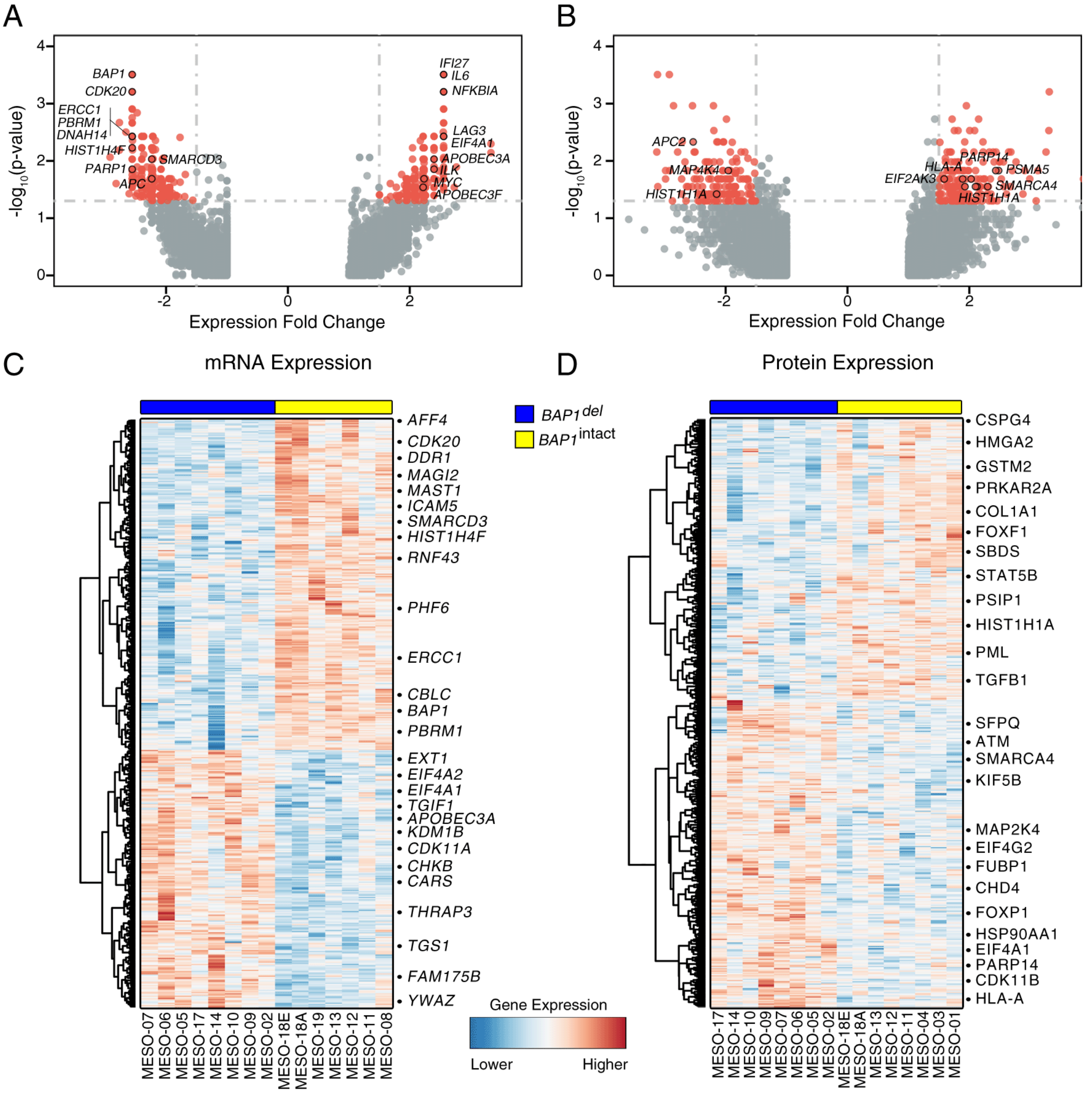

**Fig S20. Significant differentially expressed genes/proteins between molecular subtypes of PeM.** (A-B) Volcano plot showing the significantly differentially expressed genes/proteins. (A) mRNA expression, (B) protein expression. (C-D) Heatmap showing top-500 significantly differentially expressed genes/proteins. (C) mRNA expression, (D) protein expression.

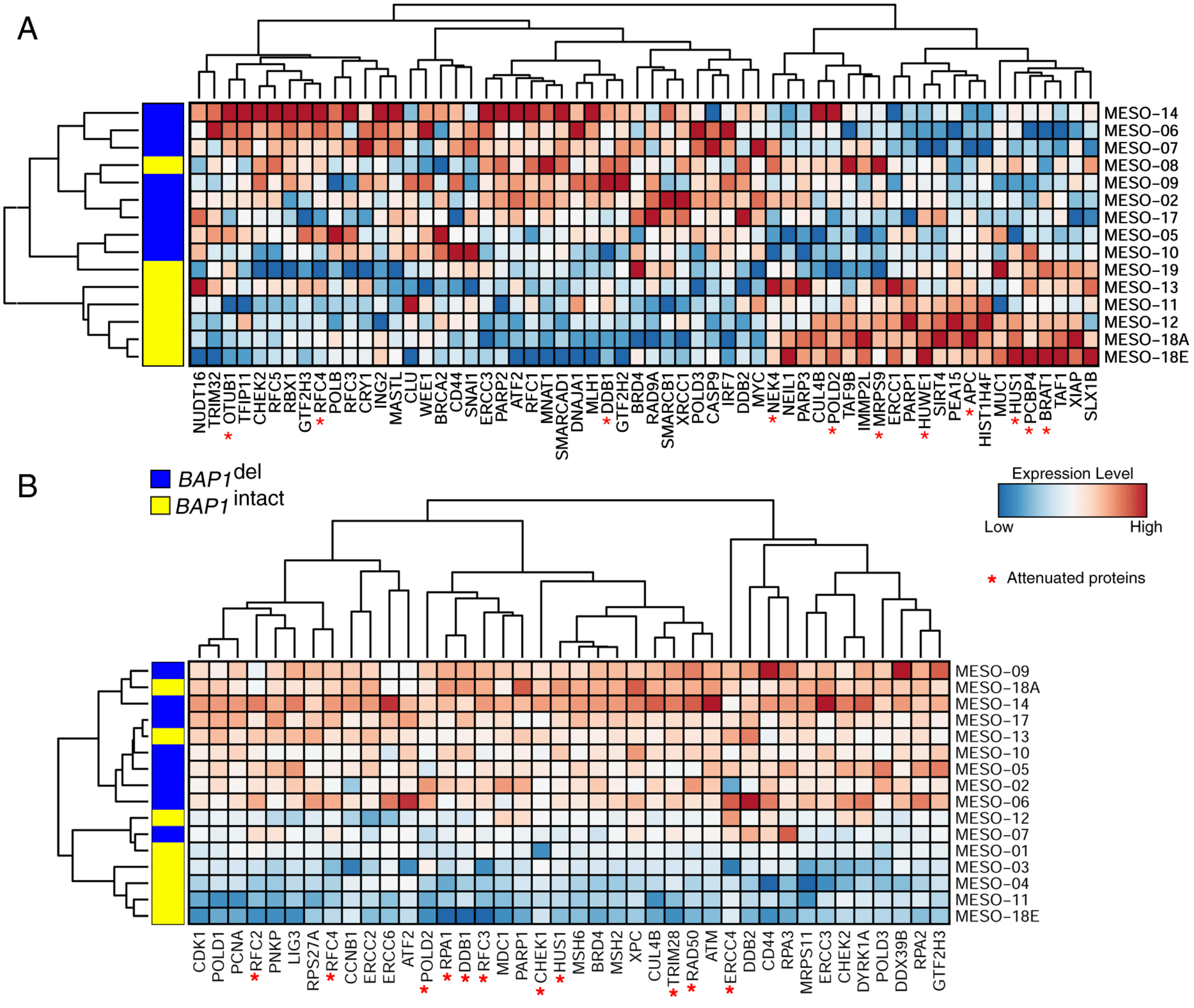

**Fig S21.** **Differentially expressed DNA-repair genes between PeM subtypes.** (A-B) Heatmap showing differentially expressed DNA-repair genes between PeM subtypes. (A) mRNA expression, and (B) protein expression.

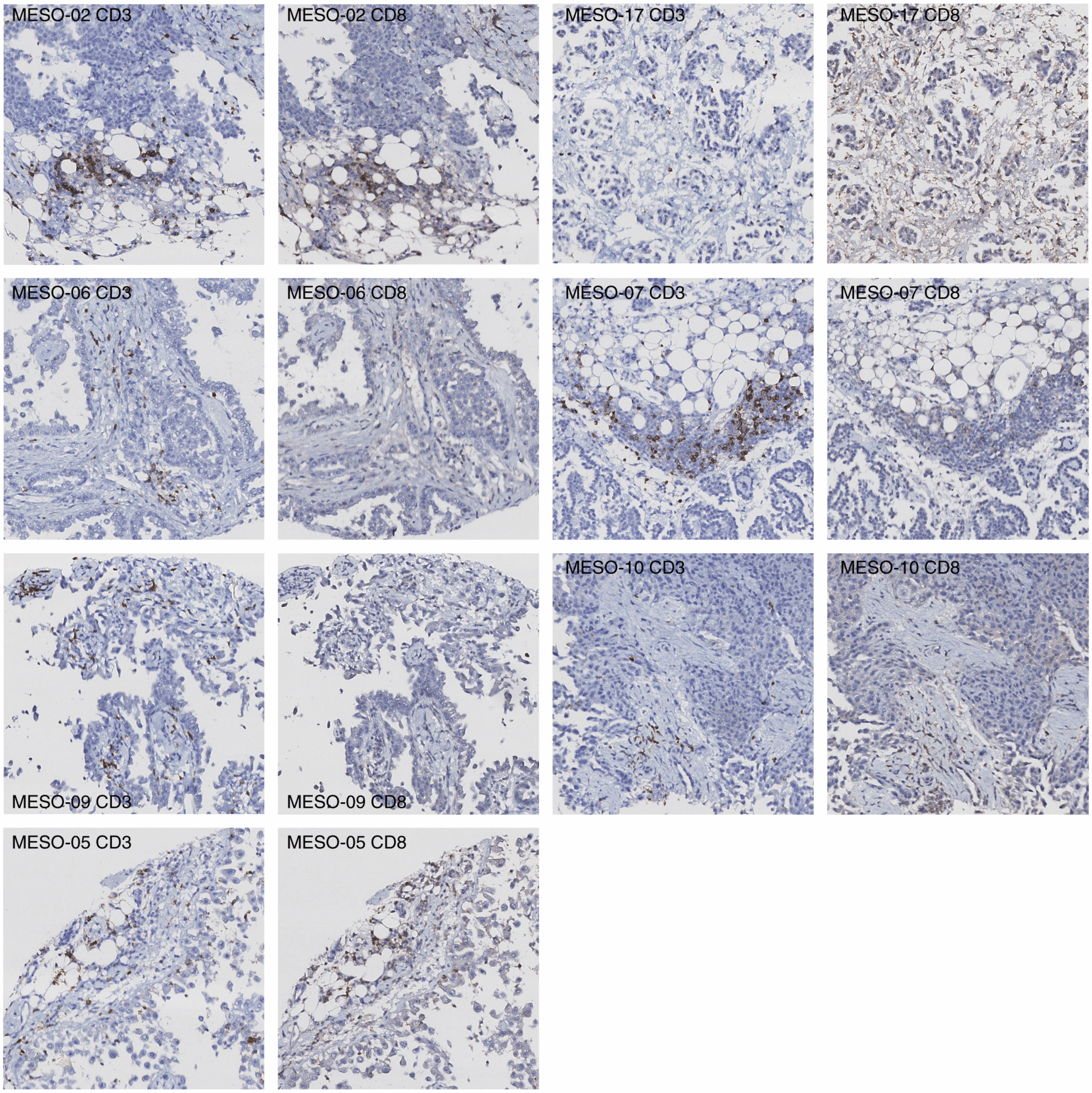

**Fig S22.** **TMA slides of BAP1$^{del}$ PeM IHC stained for CD3 and CD8.**

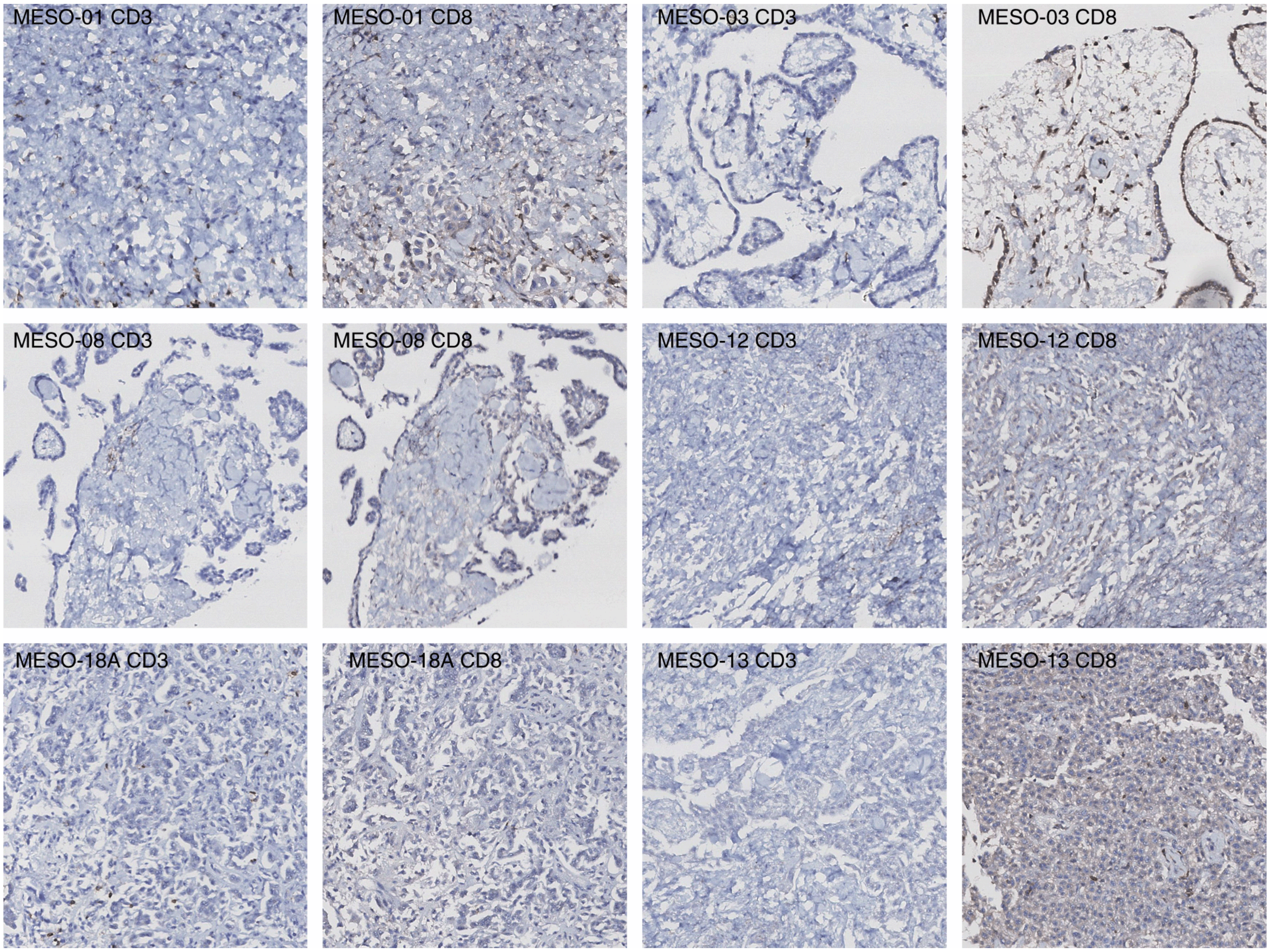

**Fig S23.** **TMA slides of $BAP1^{intact}$ PeM IHC stained for CD3 and CD8.**

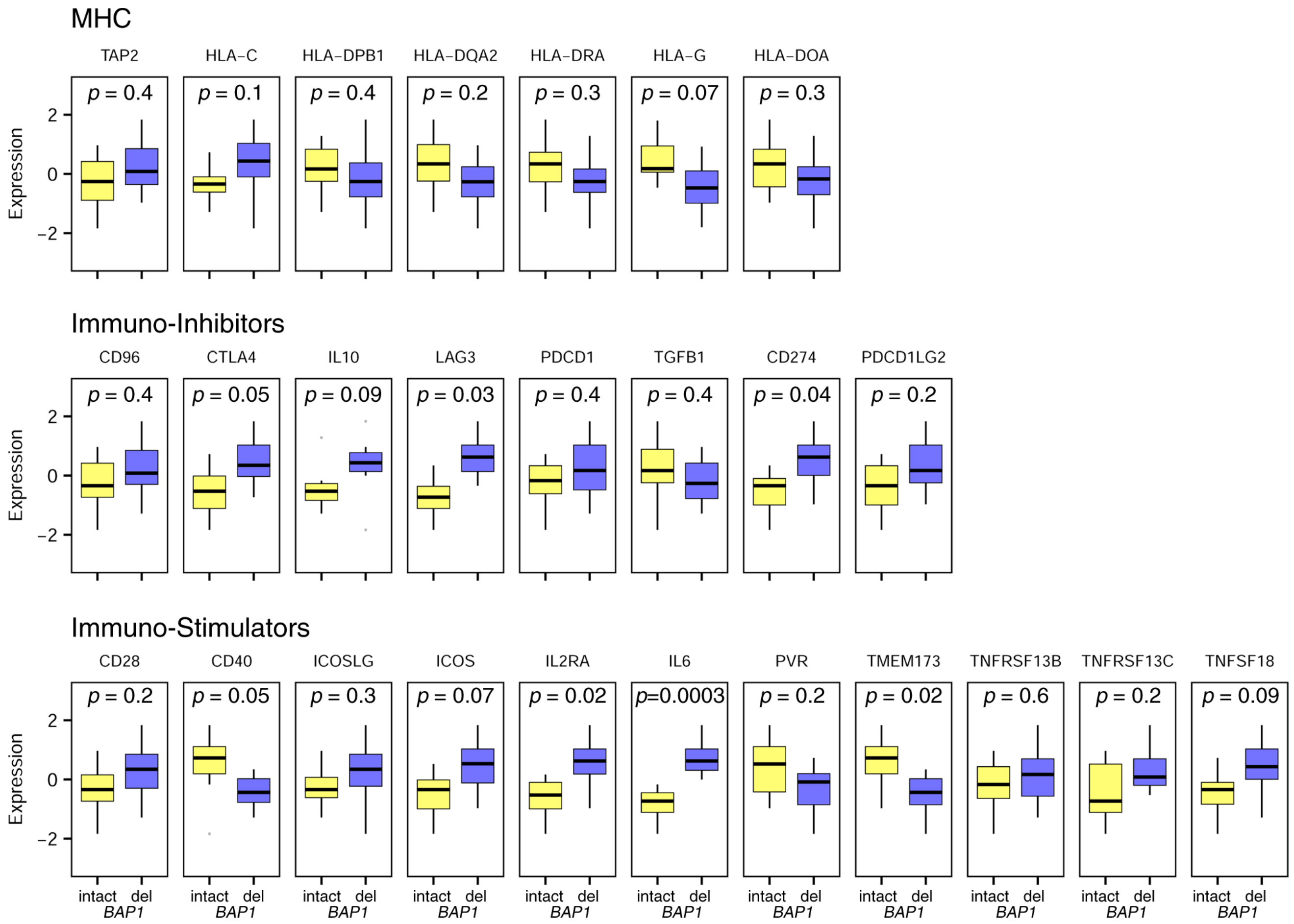

**Fig S24. mRNA expression pattern of genes involved in immune system between PeM subtypes.** The Immune system genes are grouped into three groups: MHC genes, immuno-inhibitors genes and immuno-stimulatory genes. The Wilcoxon signed-rank test p-values obtained comparing the PeM subtypes are mentioned above the plot.

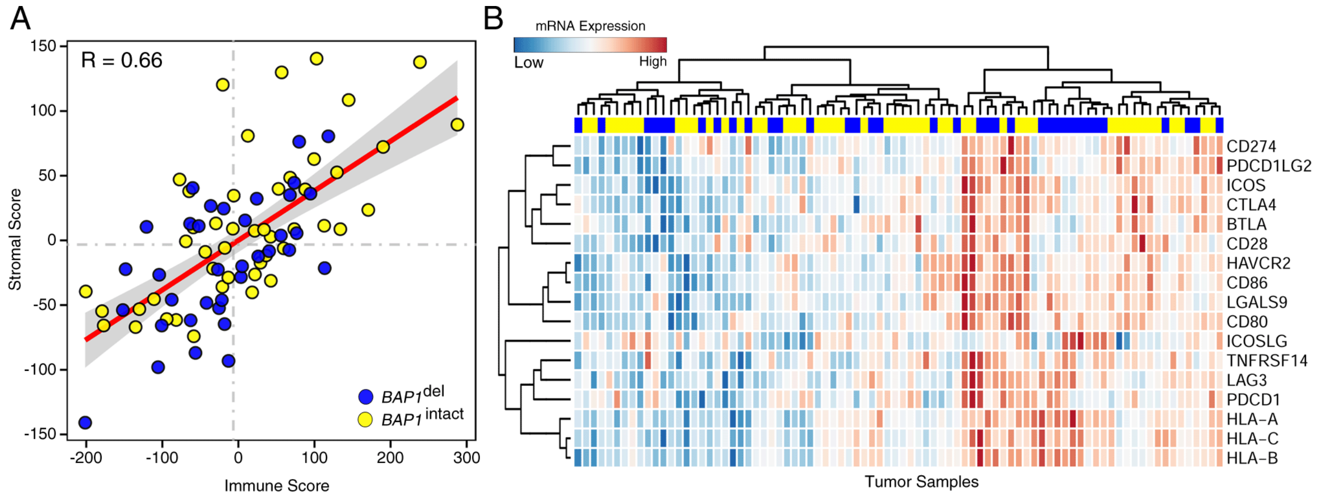

**Fig S25. Immune cell infiltration in Pleural Mesothelioma (TCGA-MESO).** (A) Correlation between immune score and stromal score derived for each tumor sample using mRNA expression. (B) Heatmap showing mRNA expression of different immune-checkpoint receptors in pleural mesothelioma tumors (TCGA-MESO)

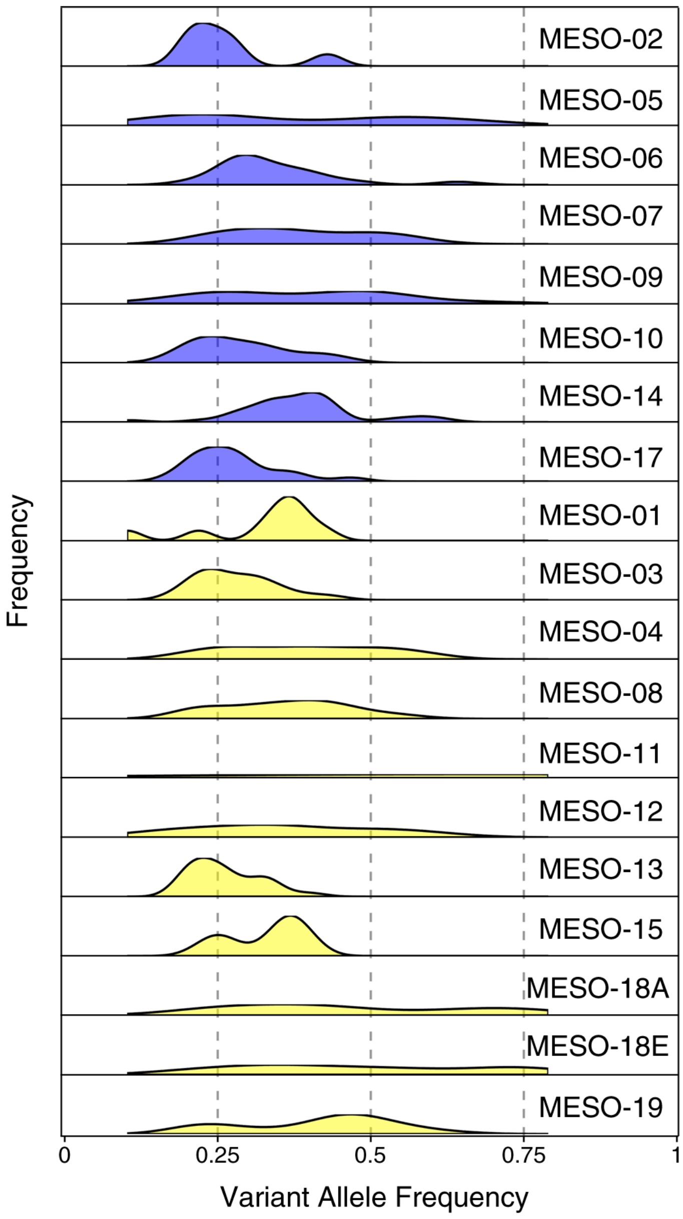

**Fig S26. Distribution of the variant allele frequency of somatic mutations in PeM.** Variant allele frequency distribution are colored by their corresponding PeM subtype BAP1$^{intact}$ (blue) and BAP1$^{intact}$ (yellow). Corresponding data is available in Supplementary Table S3.

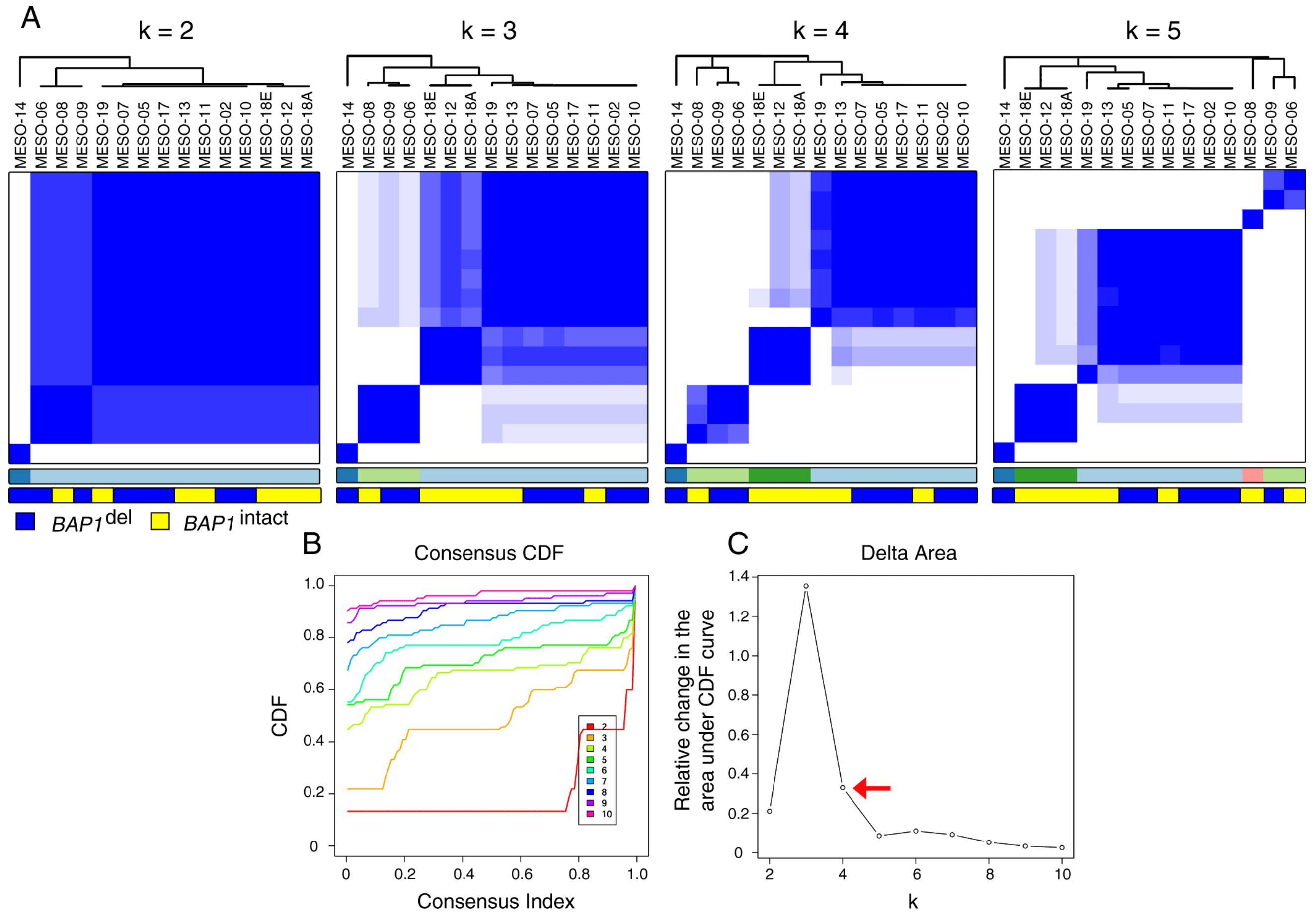

**Fig S27. Consensus clustering of long noncoding RNA (lncRNA) expression profiles of PeM.** (A) Heatmap representing consensus clustering of lncRNA expression profiles of PeM with different number of chosen clusters (k). (B) Consensus CDF curve and (C) Delta area curve. Optimum number of clusters (k=4) has been highlighted.

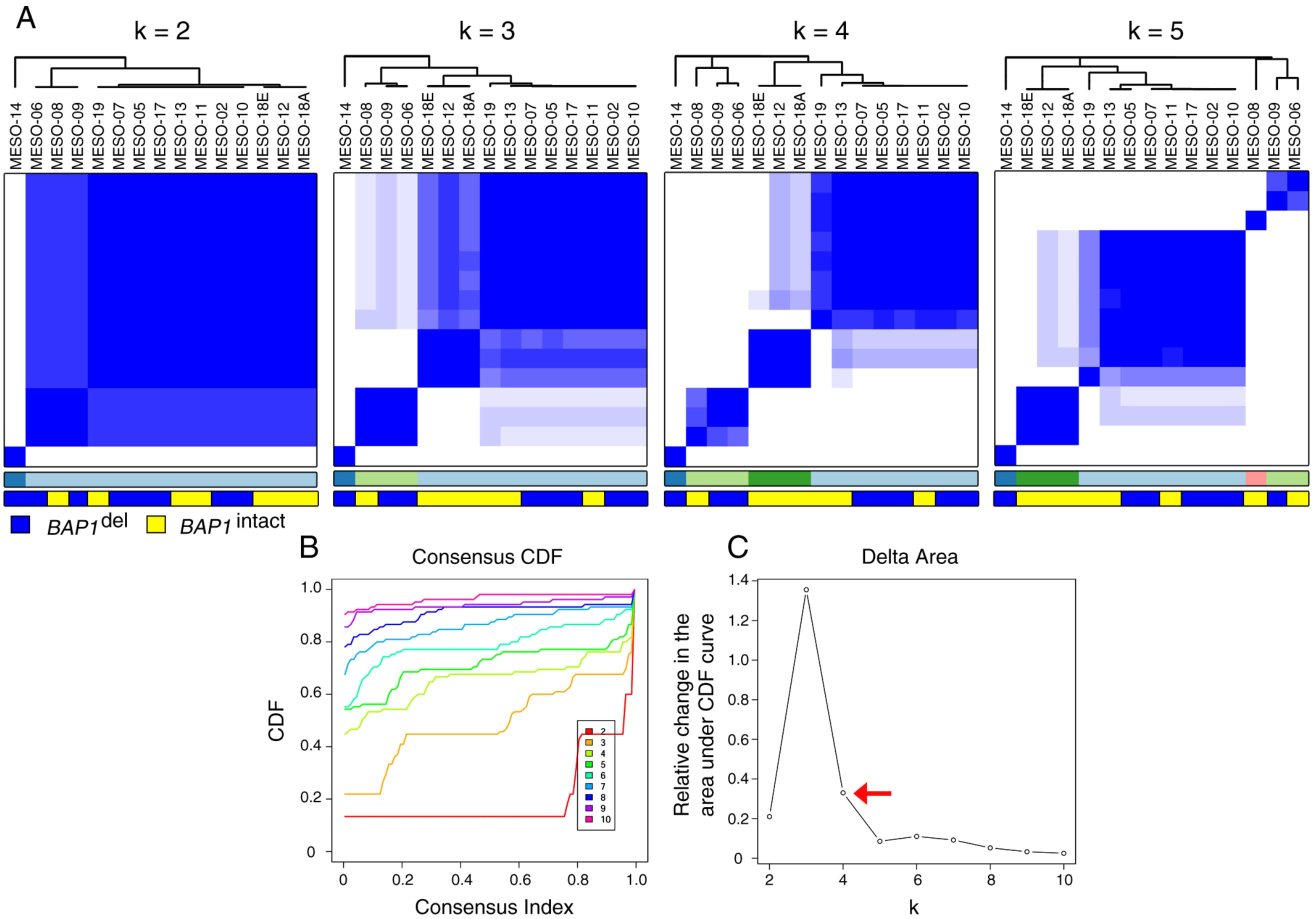

**Fig S28.** **Consensus clustering of micro RNA (miRNA) expression profiles of PeM.** (A) Heatmap representing consensus clustering of miRNA expression profiles of PeM with different number of chosen clusters (k). (B) Consensus CDF curve and (C) Delta area curve. Optimum number of clusters (k=4) has been highlighted.

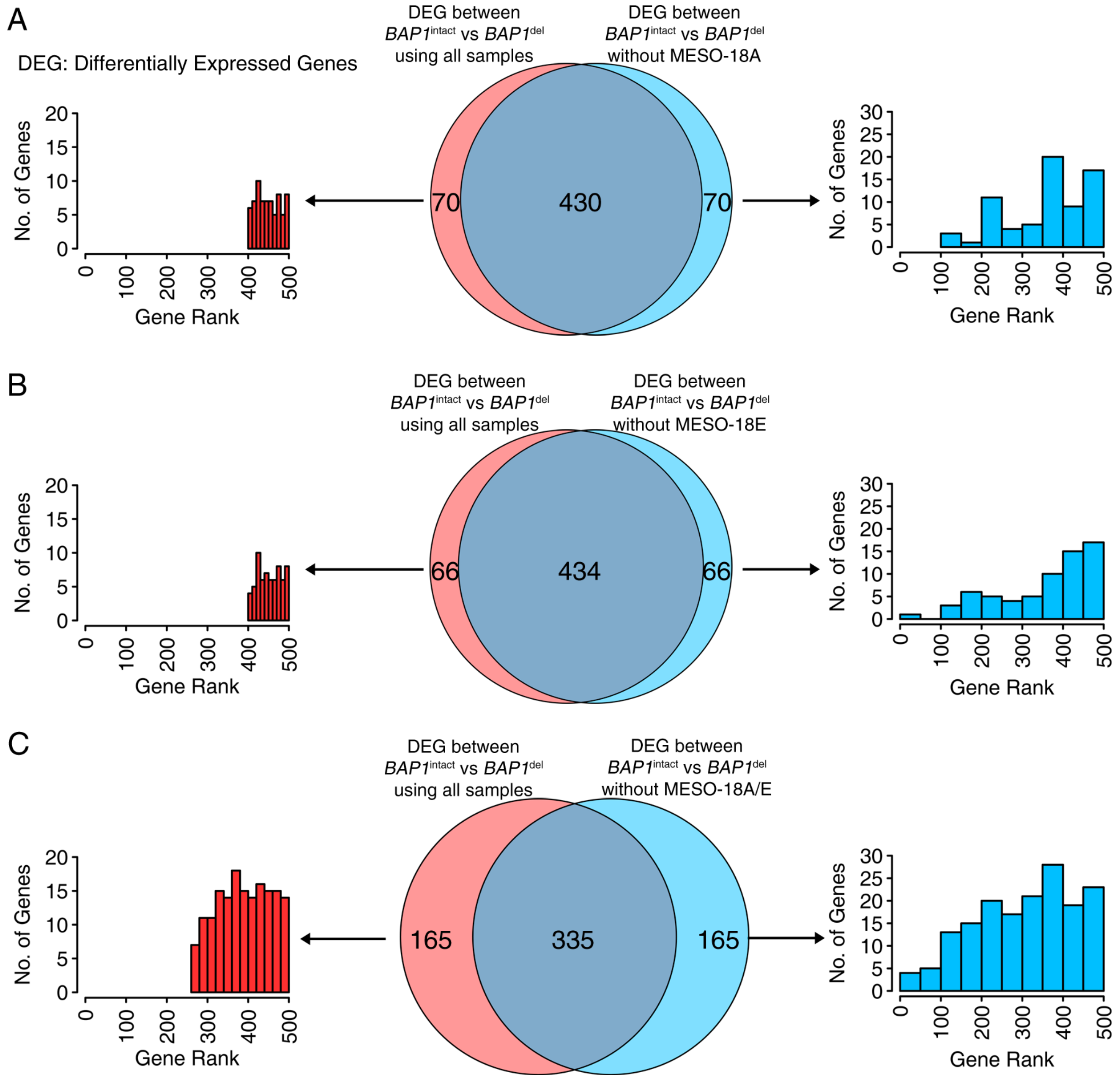

**Fig S29. Differentially expressed genes obtained when MESO-18A and/or MESO-18E are removed from the analysis.** We used Wilcoxon signed-rank test to identify the differentially expressed gene (DEG) between $BAP1^{del}$ and $BAP1^{intact}$ mesotheliomas in the mRNA expression dataset. We either used all samples or removed the sample(s) MESO-18A and/or MESO-18E for the analysis. We then selected top-500 DEG with p-value $< 0.05$. We then compared the DEG obtained using dataset (A) with all samples vs. that without MESO-18A, (B) with all samples vs. that without MESO-18E, and (C) with all samples vs. that without MESO-18A and MESO-18E. Histogram of ranked list of DEG are presented on the left and right panel.

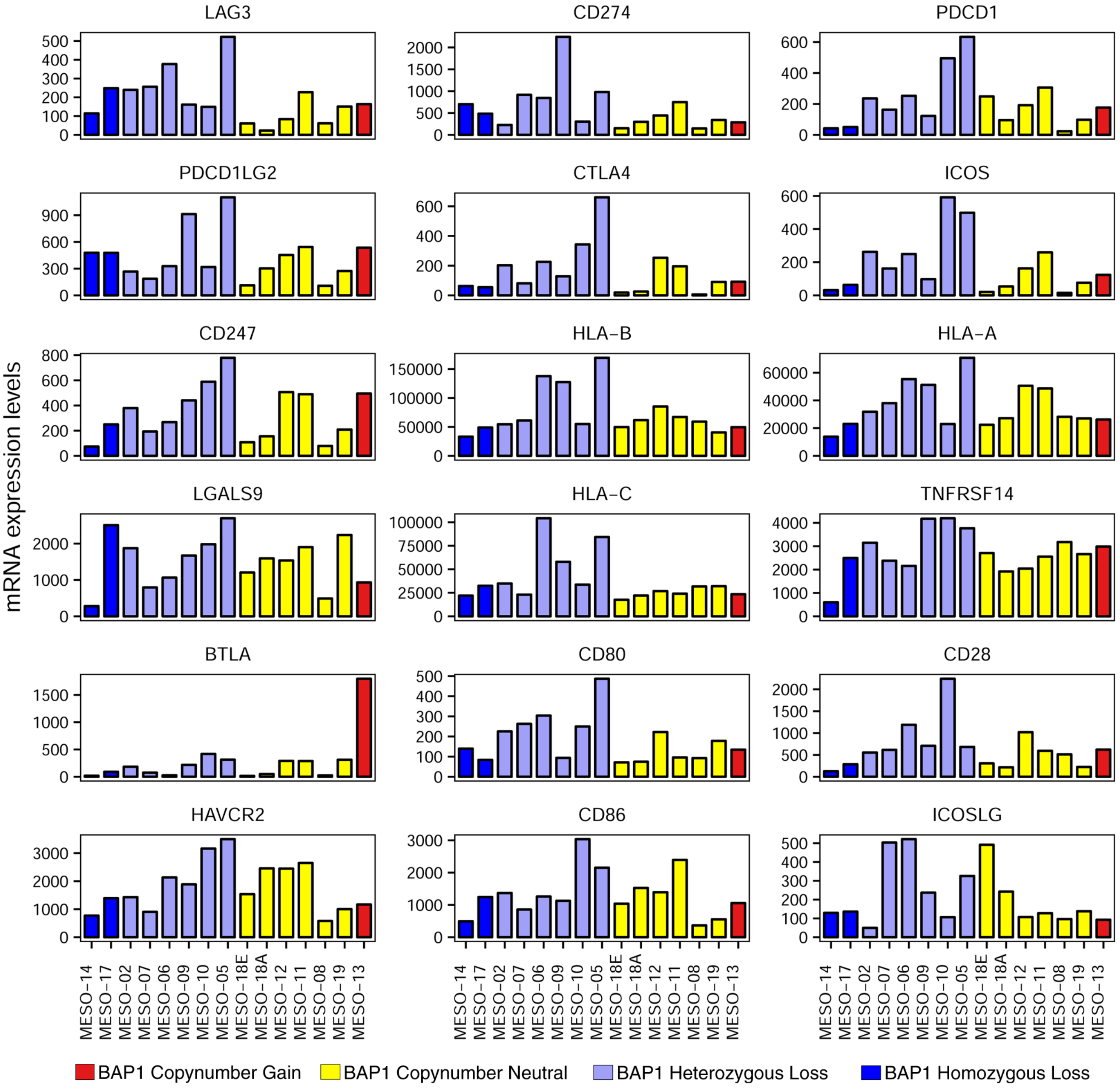

**Fig S30.** **Bar-plot of mRNA expression of different immune checkpoint receptors mentioned in Fig.4D.**
